# Plants and mammals once tracked climate closely, then mammal niches shifted under human pressure

**DOI:** 10.64898/2026.09.23.753753

**Authors:** Corentin Gibert-Bret, Benjamin R. Shipley, Julia A. Schap, Yue Wang, Silvia Pineda Munoz, Jenny L. McGuire

## Abstract

Despite recent climate change, many plants and animals are not yet tracking climate as predicted. We reconstruct realized climatic niches for 16 plants and 45 mammals across North America from the end of the last deglaciation to the present, calculating climate fidelity to measure temporal niche stability. Plants consistently exhibit higher climate fidelity than mammals, despite limited dispersal. During the deglaciation, which had lower human impact, some large mammals tracked warming climates as effectively as plants. Following European colonization, mammalian niches shifted markedly. Large mammals were displaced from warm, humid regions toward colder, drier areas, while some small mammals expanded into human-modified landscapes. In contrast, plant niches remained largely structured by climate, highlighting stronger anthropogenic constraints on mammalian responses to climate change thus far.

**Graphical Abstract:** 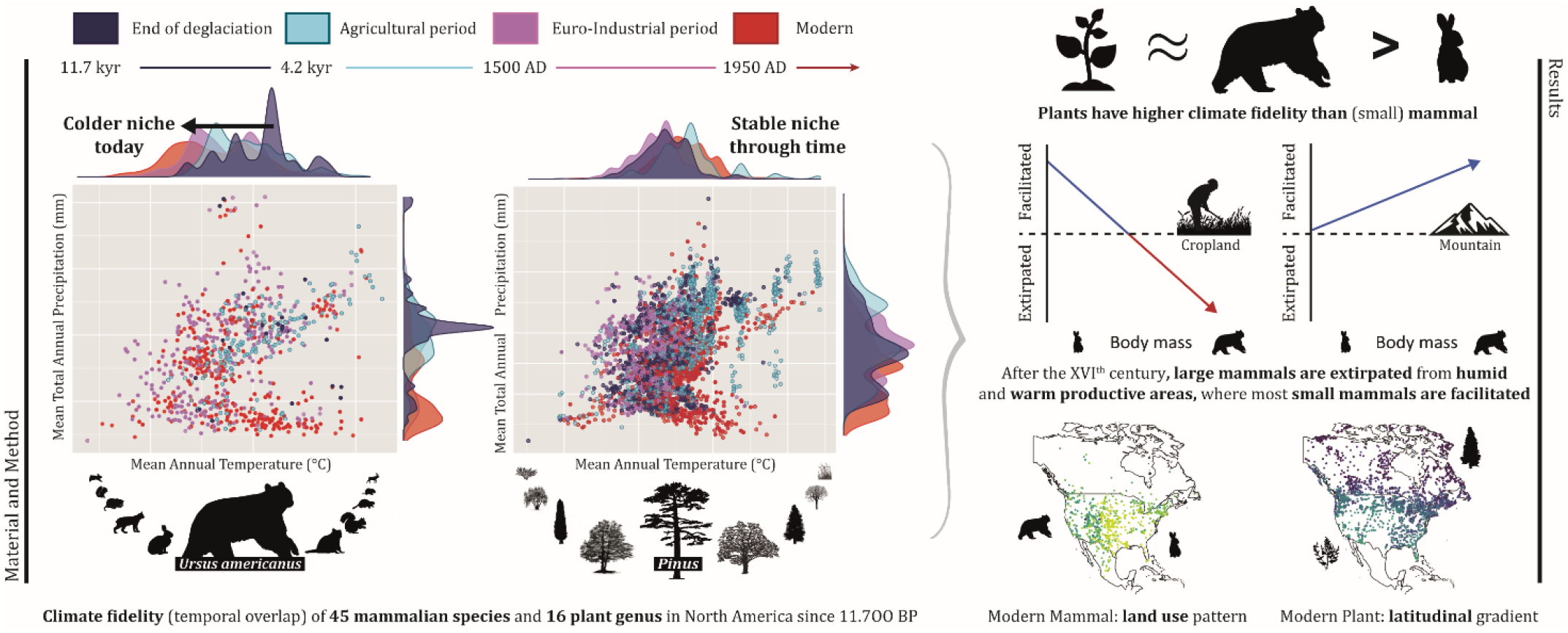

## Main Text

As climate change continues, we anticipate massive range shifts toward higher latitudes or elevations (1, 2). Forty to sixty percent of observed terrestrial species have begun to shift their distributions in manners consistent with predicted climate tracking trajectories (i.e., mainly toward high latitudes and altitudes; 3, 4, 5). Nevertheless, the dispersal of many of these species is likely to be outpaced by impending rates of climate change (6, 7). Notably, even though the 10 warmest years in the historical record have all occurred in the past decade (2012-2022; 8), half of terrestrial vertebrates have not yet begun to track changing climates (2, 9). Are these species unable to track climate, or are they unaffected by the magnitude of change they have experienced thus far?

The relationship between climate and species distributions is complex, especially on land. Given that plants are highly dependent on temperature and precipitation for biomass production, their success is more directly driven by bottom-up changes in climate than taxa at higher trophic levels (10). However, plants and animals are also highly codependent. Over half of plant dispersal is animal-mediated (11, 12), while herbivorous vertebrates depend on the identity and availability of plants to consume (13, 14). Beyond biotic constrain on dispersal, 70% of unglaciated terrestrial land has been converted for anthropogenic land use (15, 16). The resultant fragmentation reduces habitat connectivity, hinders species’ movements (17), and increases extinction risks by preventing terrestrial species from reaching potential climatic refugia (18, 19, 20). Body size, dietary generalization, and dispersal ability are all critical characteristics that decide the climate-tracking potential across today’s dynamically changing landscape (21).

Most evaluations of the effect of climate change on species distributions have been based on recent observations, limited to several decades (22). However, the cumulative effects of land use and climate change, and subsequent species’ responses, are long-term ecological processes (23, 24, 25, 26). To understand the dependence of species’ distributions on climate, we calculate the climate fidelity of North American plants and mammals over the past 11,700 years, that is, the extent to which each taxon maintains similar realized climatic niches during times of changing climates (27, 28, 29). We reconstruct the temporal dynamics of the climatic niches (mean annual temperature, MAT, and decadal mean total annual precipitation, MAP) of 16 plant taxa and 45 mammal species using fossil and modern occurrences paired with simulated and measured climate. We examine four time periods: End of Deglaciation (t_0_: 11,700 – 4,200 ybp), Agricultural (t_1_: 4,200 ybp – 1500 AD), Euro-Industrial (t_2_: 1500 – 1950 AD), and Modern (t_3_: 1950 – present) (**Fig. 1**). The first of these time periods experienced climate change, 1.15°C of warming and fluctuating precipitation regimes (30), with minimal human land uses, and the final two experienced weaker climate change but also increasing anthropogenic pressure (i.e., the arrival of Europeans and expanding urbanization and industrialization [31]). Reconstructed niches and climate fidelity are computed using the R package *ecospat* (32) following a subsampling procedure to ensure comparability of all time bins and taxa (see *supplementary methods* section for detailed methods and sensitivity analyses). Niche reconstructions account for available climate, and climate fidelity calculations are compared to null models (see *supplementary methods*). Using identical methods and data selection practices for plants and mammals (**Fig. S1**), we address the following questions: (**i**) Do plants track climate with higher fidelity than mammals or vice versa? (**ii**) Did increasing anthropogenic impacts affect climate fidelity? and (**iii**) How does the spatial distribution of climatically vulnerable communities change as anthropogenic pressure increases?

**Figure 1.**
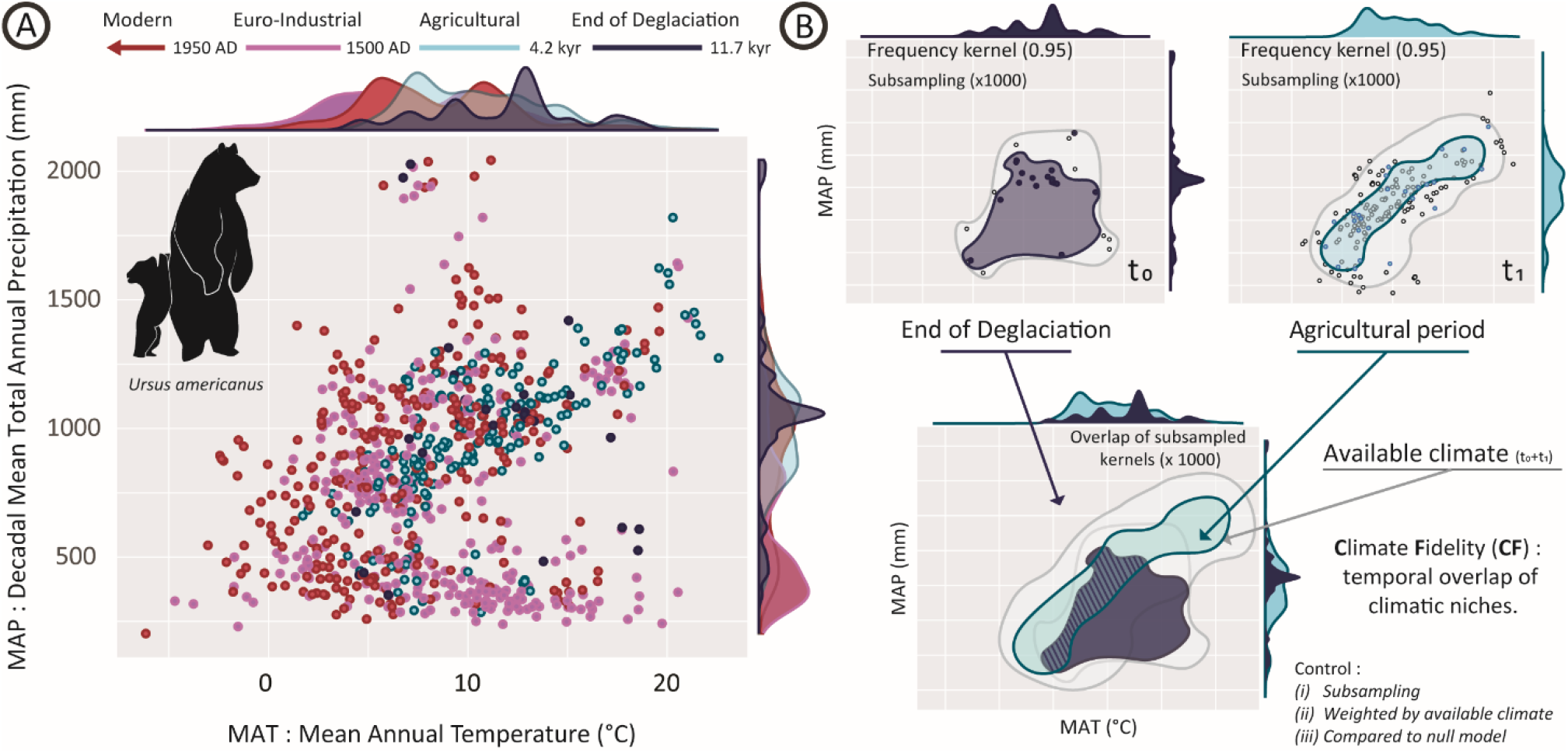
Illustration of Climate Fidelity (CF) approach using the temporal reconstructions of the climatic niche of the American black bear (*Ursus americanus*). (**1A**) The black bear’s climatic niche has shifted toward colder climates since 11,700 ybp (i.e., the kernel frequency shifts left). MAT and MAP are extracted from CCSM3 based on the mean ages of occurrences and sorted into four time periods. (**1B**) Climate fidelity is the temporal stability of climatic niches, computed as temporal niche overlap using Schoener’s *D*. The black bear’s deglaciation climatic niche (in dark blue, labeled t0) is compared with the climatic niche from the Agricultural period (4,200 ybp – 1500 AD; in light blue, labeled t1). Climate fidelity is the median of 1000 iterations of subsampling.

### Plants track climate better than mammals

Plants consistently exhibit higher climate fidelity than mammals (**Fig. 2**). We first calculated the climate fidelity of each plant and mammal based on the climatic niche temporal overlap (Schoener’s *D*) exhibited between the Deglaciation and Agricultural periods (**Fig. 2A**). By calculating climate fidelity at this time, we can evaluate the extent to which plants and animals tracked climate with the least obstruction from anthropogenic landscapes and when climates were changing the most (i.e., 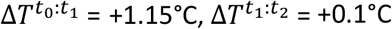). The mean climate fidelity of plants is 12% higher than for mammals 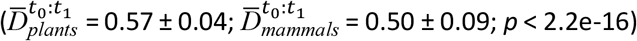 (**Fig. 2A, S2, Table S1-S2**). This difference in climate fidelity between plants and mammals becomes even more extreme (16%) when we compare modern climatic niches with older climatic 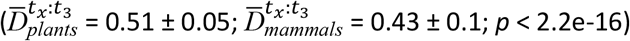 (**Fig. 2B, Table S3-S4**) or based on the comparison of any time-bin with the subsequent one 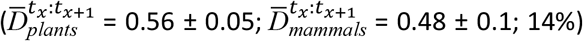 (**Fig. S2, Table S1-S2**). Although some mammals have higher climate fidelity than some plants (**Fig. 2A**), all plants are within the top 75% of climate fidelity values with low standard deviation 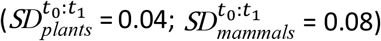 (**Tables S1-S2**). These findings are unaffected when we examine mammals at different taxonomic scales, comparable to those used for plants (**Figs. S3** and **S4**).

**Figure 2.**
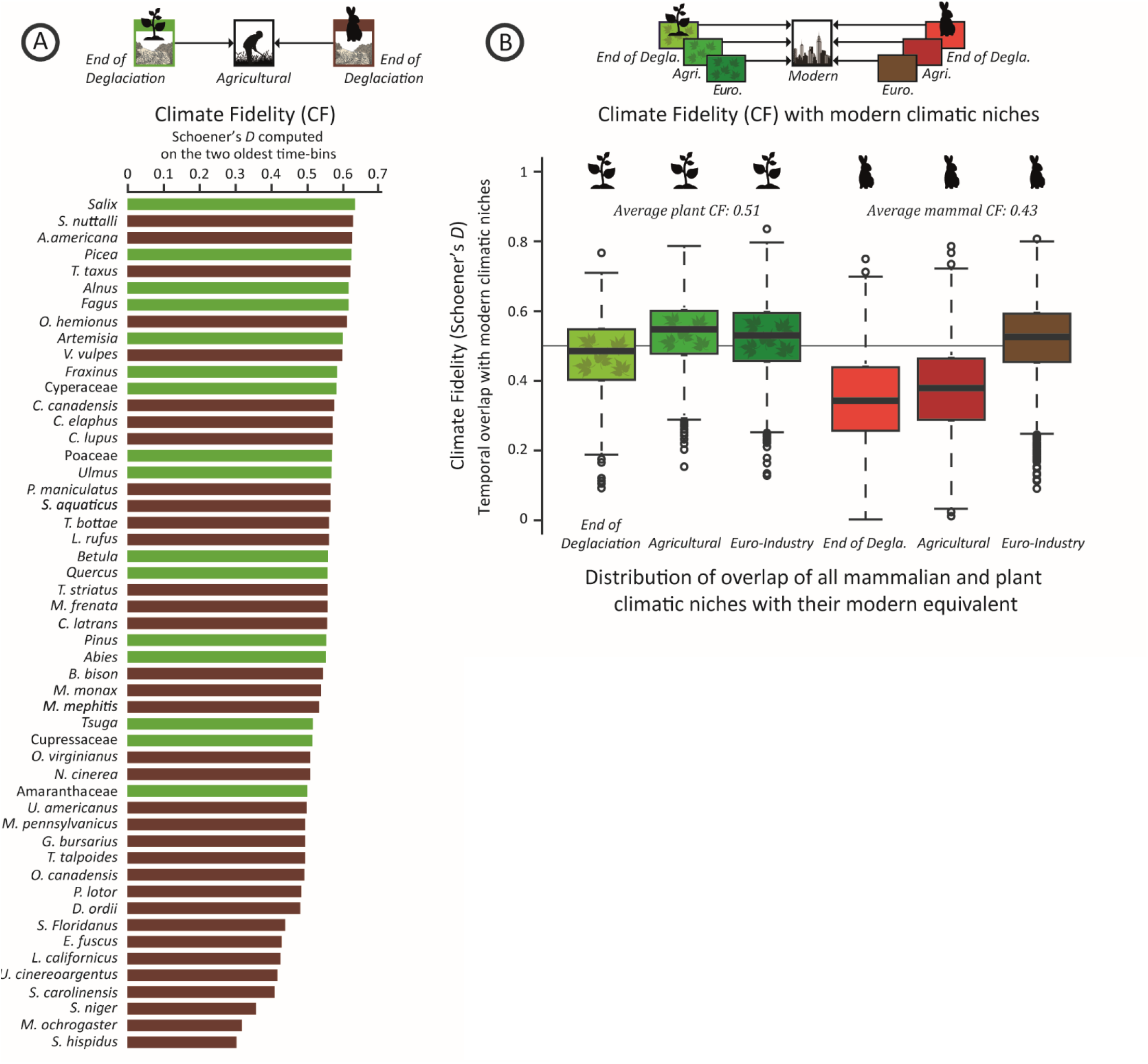
The climate fidelity of plants and mammals. (**A**) Climate fidelity computed before the arrival of Europeans in North America, comparing the Deglaciation (11,700– 4,200 ybp) and Agricultural (4,200 ybp – 1500 AD) periods (**Fig. 1)**. Plants exhibit higher overall climate fidelity than mammals (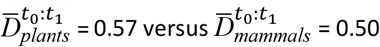, *p* < 0.01) (**B**) The climate fidelity of mammals and plants when comparing climatic niches from each time period to those of the Modern period. Mammalian climatic niches changed significantly more with respect to Modern after the arrival of Europeans and the industrialization of North America (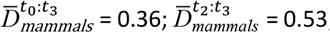, *p* < 0.01).

The plants with the highest climate fidelity, willow (*Salix*: 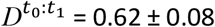), spruce (*Picea*: 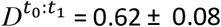), or beech (*Fagus*: 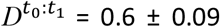), kept pace with quite mobile mammals, including the pronghorn antelope (*Antilocapra americana*: 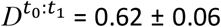), the American badger (*Taxidae taxus*: 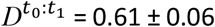), and the mule deer (*Odocoileus hemionus*: 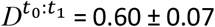) (**Fig. 2A, Table S1-S2**). Further, plants that exhibit long-distance dispersal do not have significantly higher climate fidelity (Δ*D* = 0.003, *p* = 0.61) (**Fig. S5**). Interestingly and unexpectedly, herbivores also do not exhibit higher climate fidelity than carnivores ( 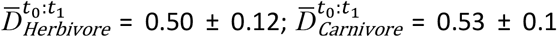 *p* = 2.2e-16) (**Fig. S6, S7**). Therefore, the intuitive perspective of mobile organisms tracking climate better than sessile organisms (12, 33, 34) is flawed for plants, and the hypothesis of herbivores tracking plants (13, 14) is deficient. These results suggest that, in general, environmental filtering is acting more strongly on plants than mammals.

When we consider only mammals, larger mammals (>5 kg) exhibited the highest climate fidelity (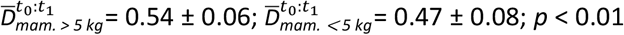 *p* < 0.01) (**Fig. S8**). Smaller mammals like the prairie vole (*Microtus ochrogaster*: 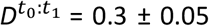) or the hispid cotton rat (*Sigmodon hispidus*: 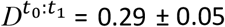), on the other end of the spectrum, possess much lower climate fidelities 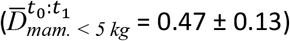 than larger mammals and lower climate fidelity than the lowest plants (Amaranthaceae: 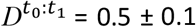). This relationship between body size and climate fidelity may be the direct consequence of the relationship between body size and velocity, where the larger the taxa, the faster it can move through the landscape to disperse toward favorable climates (35) (**Fig. S9**). Consequently, small mammals, known for their tight relationship with climate (36, 37, 38, 39, 40) are at risk of being unable to track climate in the near future, especially given the high rate of projected change (33) (**Fig. S10-S12**). However, the low climate fidelity of small mammals could also be the consequence of their ability to buffer climate disturbance by moving between microhabitats (41, 42) and adjusting their behavior (e.g., feeding habits, burrowing). It must be noted that these local adaptations to environmental change have already been observed in generalist species like the bank vole when specialist species become restricted to their shrinking optimal habitat (43). Moreover, these adaptations could be insufficient to mitigate the consequences of the ongoing climate change, especially as modern spatial patterns of diversity of burrowing species are inversely correlated to annual temperature (44).

### The growing impact of human activities on climate fidelity

By comparing climatic niches from each time period with the Modern period (**Fig. 2B)**, we find that the mammalian climatic niches have been more strongly impacted by the growing transformation of the landscape by human activities than by climate change. The arrival of Europeans and the industrialization of North America profoundly changed the climatic niches of mammals, while only slightly modifying plants’ climatic niches. The climatic niches of mammals during the Deglaciation (11,700-4,200 ybp) and Agricultural (4,200 ybp – 1500 AD) periods strongly diverge from their Modern period’s niches 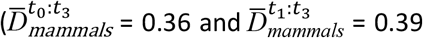 *p* < 0.01) (**Fig. 2B, Table S3**). This is opposed to mammalian climatic niches during the Euro-Industrial period (1500 – 1950 AD), which strongly overlap with those of the Modern period 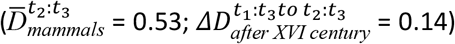 (**Fig. 2B, Table S3**). In contrast, plants were only slightly affected by the arrival of Europeans and the onset of Industrialization 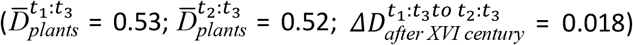 (**Fig. 2B, Table S4**). However, grasses, forbs, and sedges (Amaranthaceae, *Artemisia*, Cyperaceae) and certain trees (*Cupressaceae, Fagus, Fraxinus*) are now characterized by drier climatic niches than before (**Fig. S13**).

Given that we observed a major shift in mammalian climatic niches at the Euro-Industrial period, we sought to determine whether these changes correspond with geographic regions that contain climates now dominated by anthropogenic landscapes. First, we computed the modern climatic niches of five landscapes: cropland, urban, forest, grassland, and mountain. Then, we evaluated the overlap of the climatic niches of each taxon (plants and mammal) relative to the modern climatic niches of each of these five landscapes. We calculated the change in overlap before and after the advent of the Euro-Industrial period (1500 AD). Finally, the differences in overlap create a metric (**Fig. 3B**), called the climate exclusion index (CEI), that tells us the extent to which a plant or animal has expanded into (i.e., positive CEI) or been extirpated from (i.e., negative CEI) the portion of climate space now occupied by a particular landscape (based on Pineda-Munoz 2021 [45]; see also Methods).

**Figure 3.**
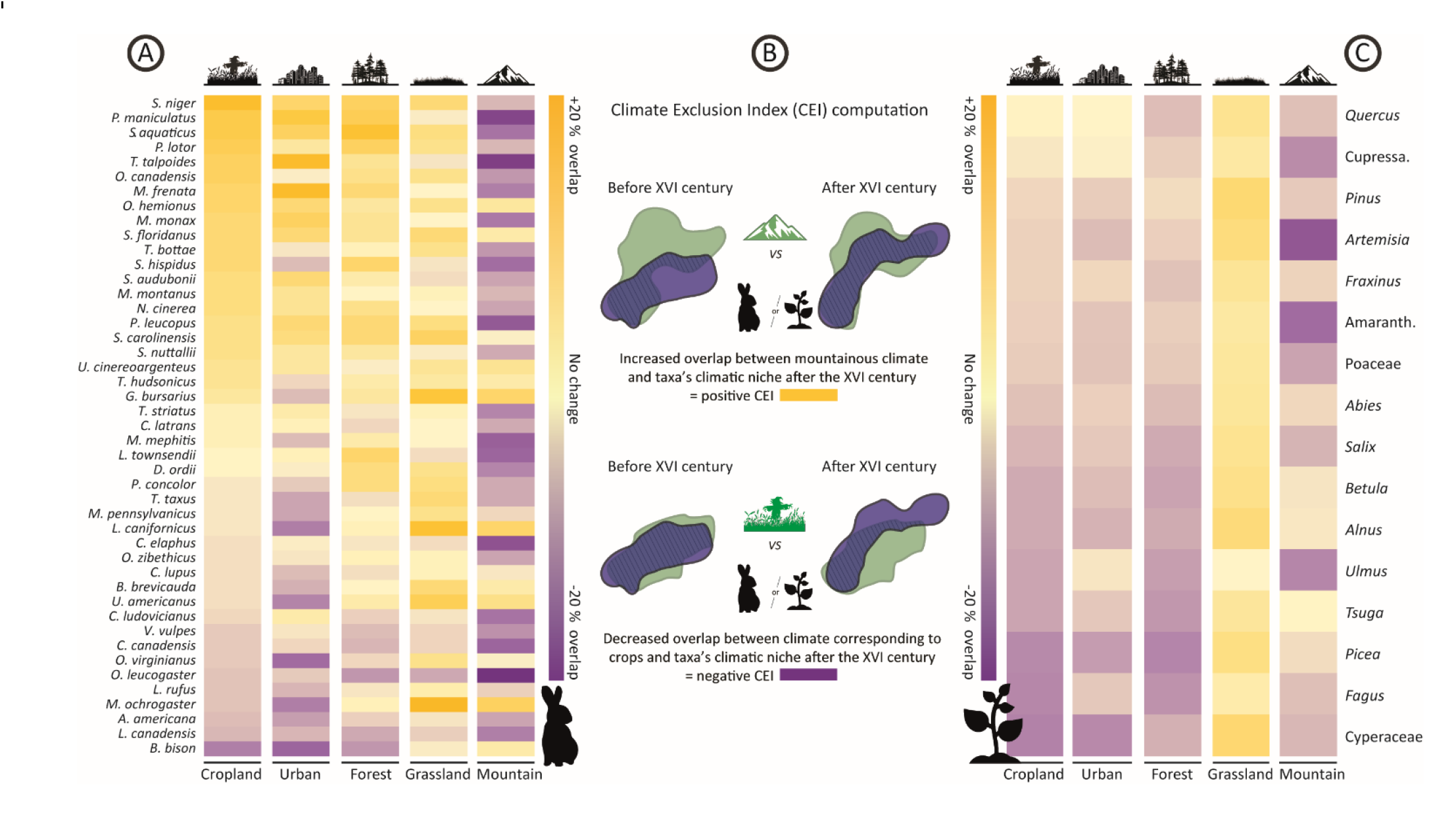
Climate exclusion index for mammals (CEI). The climate exclusion index (CEI) is calculated as the difference between the overlap of the modern climatic niches of five landscapes and the mammal and plant climatic niches before and after the arrival of Europeans (1500 AD). The CEI is sorted by changes in (**A**) mammals’ and (**C**) plants’ climatic niches relative to the crop climatic niche. Gold indicates increased overlap with a landscape’s climate after the arrival of Europeans (i.e., facilitation or commensalism) and purple refers to decreased overlap with landscape’s climate (i.e., extirpation). (**B**) A diagram explaining CEI index calculation. The pink area corresponds to mountain climate (top), green is crop climate (bottom), blue is a taxon’s climatic niche, and the hatched area indicates the overlap between landscape and taxon’s climate (niches) in environmental space. If the overlap increases after the XVI century, the CEI will be positive, if overlap decreases, the CEI will be negative. CEI is corrected for available climate before versus after the XVI century (see more detailed explanation in supplementary method section).

In general, we found that mammals and plants (e.g., bison and spruce) have experienced contractions of their climatic niches (**Table S5-S6**) in parts corresponding to the climates found in anthropogenic landscapes (**Fig. 3**). This contraction was stronger within mammals (min CEI_Urban-Cropland_ = -0.2) than plants (min CEI_Urban-Cropland_ = -0.17). However, some mammals (*urban*: 41% of mammals, *cropland*: 54% of mammals; but no plants) have expanded their climatic niches into these anthropogenic landscapes (**Fig. 3A**). Mammals that are contracting out of urban climates, such as the eastern meadow vole (*Microtus pennsylvanicus*), the American black bear (*Ursus americanus*) or the Castor (*Castor canadensis*), are expanding their climatic niches into montane regions, and vice versa. For many mammals, forests, grasslands, and occasionally mountains can act as land-use refugia, with 61%, 59%, and 22% of mammals, respectively, expanding their climatic niches into those of these landscapes. For plants, contrary to mammals, climates associated with cities, croplands, forests, or mountains do not facilitate any taxa (**Fig. 3C**). Grasslands are the sole landscape whose climates now host more taxa than before industrialization and the arrival of Europeans (**Fig. 3C**). This last observation could be the consequence of reduced grazing of tree shoots associated with large mammal extirpation (46, 47) from the most anthropized landscapes of North America (45). Forests and crops (more than cities) depict similar trends shows that forest, do not consistently act as refugia at the continental scale.

Even though large and well-moving mammals had higher climate fidelity between Deglaciation and Agricultural periods (**Fig. S8-S9**), they were preferentially extirpated from climates occupied by urban areas and croplands since the arrival of Europeans 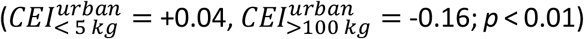 (**Figs. 4A-4B, S14, S15**). Small- and intermediate-sized mammals have offset these losses by expanding into climates associated with forests (**Fig. S15**), all mammals have expanded into grasslands, and the largest mammals (>100 kg) have especially expanded into montane climates (**Fig 4C**). In fact, the climate niches of large and intermediate-sized mammals like the black bear (*U. americanus*, ΔT_after XVI century_ = - 2.3°C, ΔP_after XVI century_ = - 281 mm) (**Fig. 1**), the river otter (*L. canadensis*, Δ_after XVI century_ = - 1.8°C, ΔP_after XVI century_ = - 50.4 mm) (**Fig. S12F**), and the elk (*C. elaphus*, Δ_after XVI century_ = - 0.3°C, ΔP_after XVI century_ = - 334.5 mm) (**Fig. S10D**) reflect their geographic shift away from warmer, wetter areas favorable to cities and agriculture, like the temperate plains or the Atlantic and Gulf Coasts (**Fig. S16**). Taken together, this implies that these large species were relegated to some of the less productive portions of their climatic niches (48). Conversely, 41% (toward Urban) and 54% (toward Cropland) of small mammals, including the valley gopher (*T. bottae*) (**Fig. S10M**) or the fox squirrel (*S. niger*) (**Fig. S11L**), strongly extended their climate niche since the arrival of Europeans, likely because they have found favorable microclimates, habitats, and abundant, supplemented food sources (49, 50, 51, 52).

**Figure 4.**
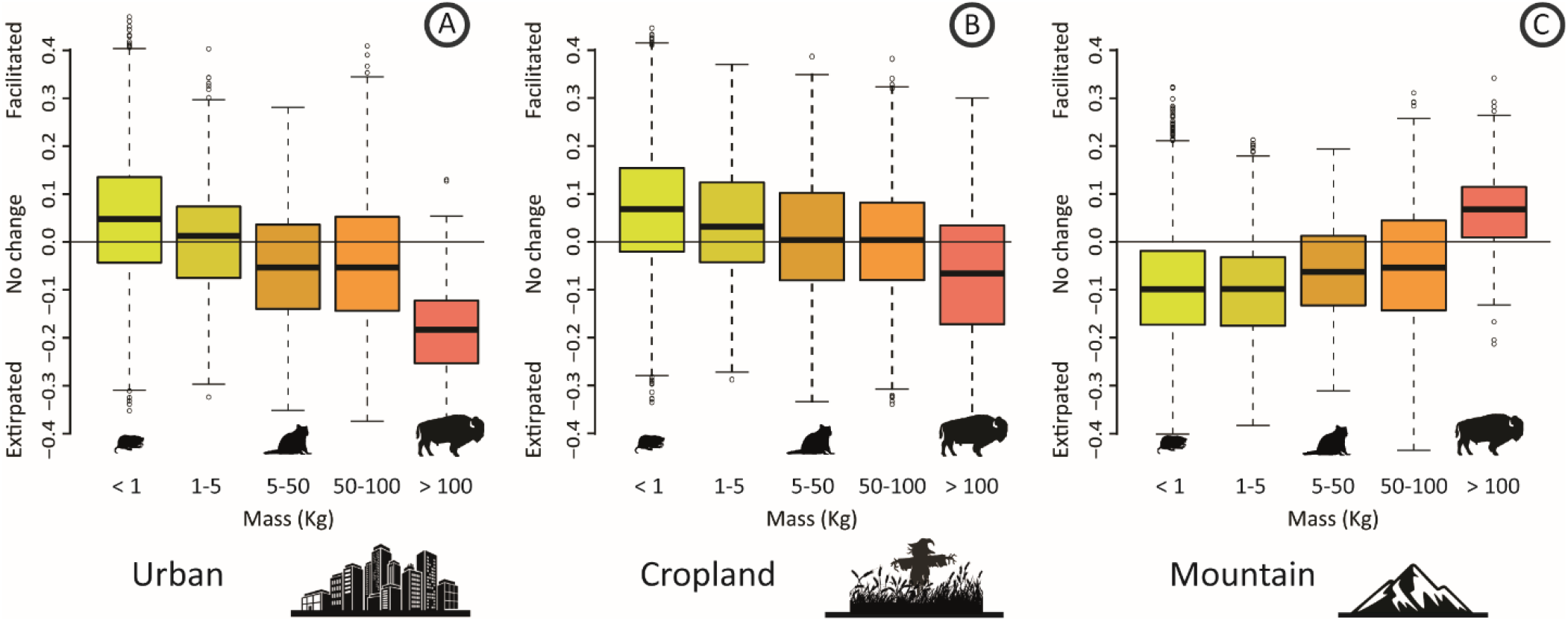
Relationship between five classes of mammalian body mass (in kg) and climate exclusion index (CEI) for three landscapes. See more landscapes (i.e., grasslands and forests) in **Fig. S14**. Positive CEI values are related to facilitation after the XVI^th^ century. Negative CEI values are related to extirpations from the targeted landscape’s climate after the XVI^th^ century. Natural areas transformed today into (**A**) urban zones or (**B**) crops. (**C**) mountain areas.

### The impact of human activities at continental and community scale

Because plants and mammals are not affected to the same extent by land use change through time, modern plant and mammal communities depict very dissimilar spatial patterns of climatic fidelity (**Fig. 5**). We averaged the climate fidelity score 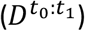 of each plant or animal taxon at the community scale (see **Fig. 2A** and **Table S1-S2**). The spatial distribution of mammalian climate fidelity has changed considerably over time (**Fig S17**). Before 1500 AD, mammalian communities in vast parts of the Great Lakes region as well as in the temperate and Northern Plains were composed of a higher proportion of taxa with higher climate fidelity than today (**Figs. S16, S17B**). Modern mammal communities were profoundly affected by anthropogenic landscapes, to the point where extirpations lead to the disappearance of latitudinal gradients of climate fidelity that were present during the End of the Deglaciation (**Fig. S17D**) and the emergence of spatial patterns matching the distribution of human impacts (**Fig. 5**: *R* = -0.38, *p* < 0.01). After the 16^th^ century, mountains acted as refugia for mammalian species with high climate fidelity, i.e., the largest mammals (**Fig. 5**: *R* = 0.3, *p* < 0.05).

**Figure 5.**
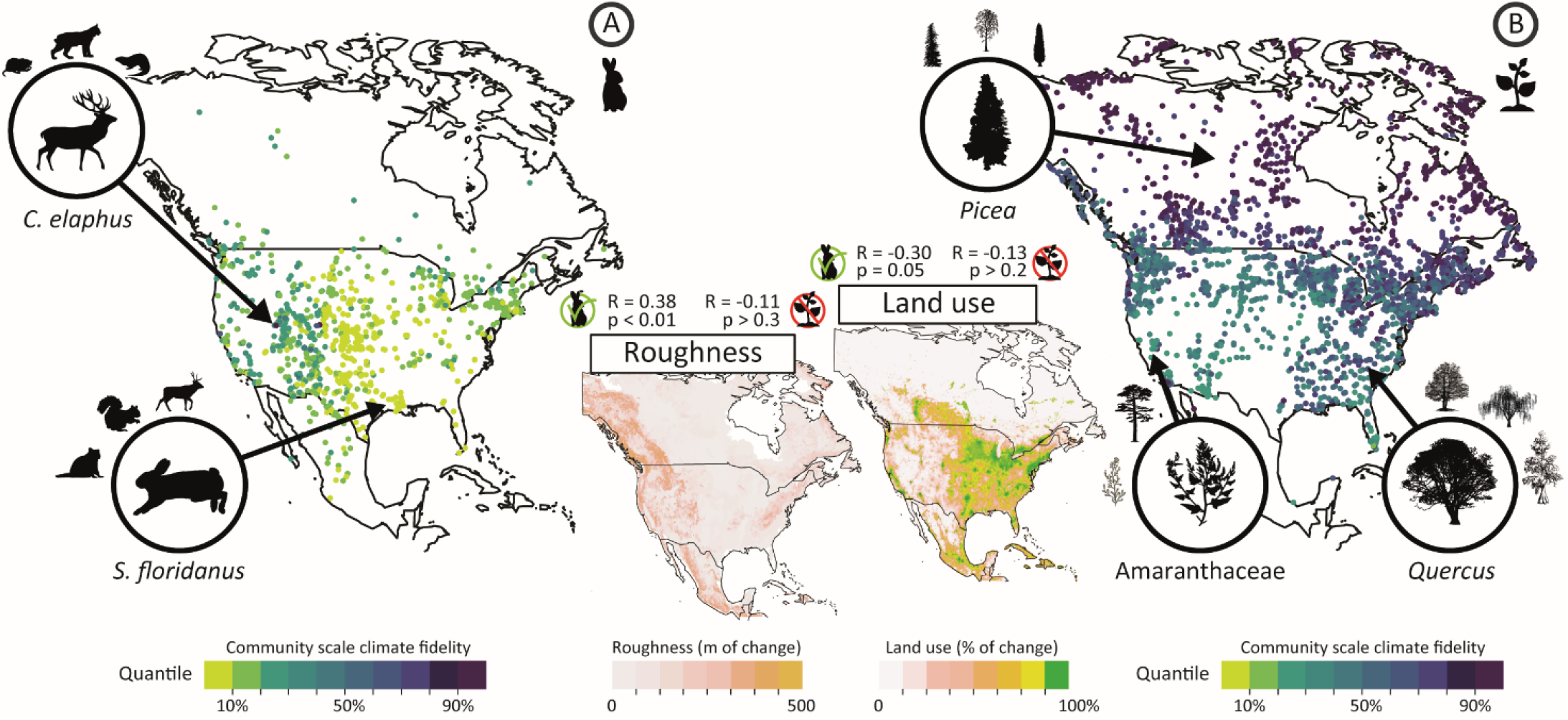
The spatial distribution of modern community climate fidelity for mammals (A) and plants (B). The climate fidelity of individual taxa, based on the comparison of the end of the Deglaciation and Agricultural periods (see **Fig. 2A**), are averaged at the community scale based on the composition of modern communities. Mammal and plant silhouettes depict examples of common taxa in the region. In the central maps, roughness (from Danielson & Gesch 2011 [58]) and land use (from Theobald et al. 2020 [59]) both correlate with mammal community climate fidelity, but not plant community climate fidelity.

The spatial distribution of climate fidelity for plants communities, however, is more consistent over time (**Fig. S16A**), and neither land use nor mountains constrain the distribution of modern plant climate fidelity. The climate fidelity of plants communities is primarily shaped by the latitudinal distribution of biomes and climates in North America (**Fig S16C**), a pattern found in mammal climate fidelity only during the end of the Deglaciation (**Fig. S16D**). Climate fidelity patterns across North America suggests that all biomes are not equally sensitive to climate change (53). Boreal, coastal, deciduous, and mixed forests could struggle to find future climate refugia in the light of the unprecedent rate of warming happening today. Counterintuitively, mammal communities characterized by the presence of higher climate fidelity mammals in the west and north may be more vulnerable than central and eastern communities to ongoing climate change, especially if they are associated with mountains, which can become isolated islands limiting the ability of species to disperse towards climatic refugia (54, 55, 56, 57).

### Climate fidelity and conservation

Plants and many larger mammals (>5 kg) exhibited relatively high climate fidelity (*D* > 0.50) between the End of the Deglaciation (11,700 – 4,2000 ybp) and the Agricultural periods (4,200 ybp – 1500 AD) while many small mammals exhibited lower climate fidelity (**Fig. 2A**). Climate fidelity can therefore reveal the relative innate vulnerability of taxa to the ongoing climate change. Many large mammals will be at risk, especially those that exhibited high climate fidelity but that are very sensitive to anthropogenic landscapes (**Figs. 2A, 3, 4**), such as the American bison and river otter. Conversely, some small mammals have become commensal species that benefit from crops and cities (e.g.the fox squirrel, *Sciurus niger*, or the northern pocket gopher, *Thomomys talpoides*). They exhibit relatively low climate fidelity, and they may find food and microhabitats suitable for them under climate change in human-impacted areas (43, 60, 61). However, the taxa most at risk in this study are the mammals that are not facilitated by anthropogenic landscapes (**Fig. 3**), that exhibited high climate fidelity for warm and humid habitats (**table S1**) and/or strong climatic niche change after the arrival of Europeans (**table S7, S10E-S10O**), such as most large mammals tested here like *Canis lupus, Lynx rufus, Ursus americanus, Antilocapra americana* and *Bison bison* or small mammal like the California jackrabbit (*Lepus californicus*) and the prairie vole (*Microtus ochrogaster*) that already have been extirpated from Texas and Louisiana (62).

When compared to the projected climate connectivity of the contiguous U.S. (18) with the climate fidelity of modern plant and mammal communities (**Fig. 5**, **Fig. S18**), we find that many of the mammals that exhibit high climate fidelity are also in areas of high climate connectivity, notably the Rocky Mountains and Pacific Northwest. However, some communities, including those in the plains and around the Great Lakes, will need increased climate connectivity to thrive. For plants, will be most critical along the Appalachian Mountains, a highly fragmented landscape in dire need of increased habitat connectivity (63).

## Supporting information

Extended version of the method section

## Authors contributions

CGB performed all the analysis and created all figures, CGB is the main contributor to the text. BRS, YW, SPM and JLM contributed to the analysis pipeline. BRS, YW, SPM, JAS, and JLM performed interpretations of the databases and climate fidelity changes through time. All authors have contributed to rewriting, figure polishing and interpretations.

## Data, code, and materials availability

All data and codes are available in the supplementary materials, on (REQUEST TO CORRESPONDING AUTHOR) and CGB GitHub (*private before acceptance*) repository named Corentin-Gibert-Paleontology/Plants-and-mammals-once-tracked-climate-closely--then-mammal-niches-shifted-under-human-pressure.

## List of Supplementary Materials

Materials and Methods (7 pages)

Figs. S1 to S40

Tables S1 to S7

References (64 - 82)

Data are stored on DRYAD -*reviewer sharing link*- : (ASK the corresponding author please) and first author GitHub repository

