## Extended version of the method section for "Plants and mammals once tracked climate closely, then mammal niches shifted under human pressure"

*Attached to*

### Methods

*Workflow and codes*

The structure of the study, materials and methods is illustrated as a workflow in Supplementary Figure 1. An example of R code to reproduce the computation of climate fidelity and climate exclusion index is available (request to corresponding author), as well as a R Markdown script and its associated HTML file made to reproduce all the main figures of the manuscript.

*Datasets*

Fossil and modern occurrence data of North American terrestrial mammals and plant pollens were downloaded from open databases. We restricted all analyses to the Holocene from 11,700 years BP to modern (1950 AD – 2022 AD). Mammalian fossil occurrences were downloaded from NEOTOMA database (constituted of FAUNMAP database for Quaternary fossil vertebrate records) and all fossil ages were estimated from radiocarbon dating using R package Bchron and the calibration curve intCal13 (64). The ages of occurrences are computed as the average age between minimum and maximum age of the strata. Modern mammalian occurrences were downloaded from GBIF (all downloaded occurrences are available : request to corresponding author) and cleaned of observations outside human observation and preserved specimen in museum and zoo. Surface and fossil pollen samples and ages are extracted from NEOTOMA database, constituted of the North American surface pollen and the North American Bayesian-aged fossil pollen datasets (65). We compiled all pollen samples and terrestrial mammal occurrences in North America between 10°- 48°N and 48°- 140°W, excluding the Alaskan occurrences/samples due to possible inaccuracies in paleoclimate reconstructions in the region during deglaciation (66, 67) and to increase comparability between plant and mammal datasets as plants are more commonly sampled in high latitude than mammals.

In total, 16,318 pollen samples were compiled, including 4,493 surface/modern pollen samples and 11,825 fossil pollen samples from 337 sites. A total of 131,711 mammalian occurrences were included in our study, with 84,809 modern and 46,902 fossil occurrences coming from 78,102 sites.

To correct for different dispersal capabilities of pollen, reflected in abundance measurements, plant taxa are considered present or absent in the pollen samples using a set of abundance thresholds based on established taphonomic processes (68, 69). *Pinus* and *Quercus* are considered present at 5% of relative abundance, *Tsuga* at 2.5%, and all other taxa above 1%.

Mammalian traits like diet, body mass, and dispersal capabilities were extracted from Schloss et al. 2012 (33).

*Similar time bins for plants and mammalian datasets*

To be able to observe the effect of major climatic and historical events since 11,700 ybp on mammal and plant climatic niches, and more specifically, to produce comparable measurement of climate fidelity through time for both mammals and plants, we have analyzed both datasets through the same four time bins. The first time bin, named here "End of the Deglaciation" ($t^0$), goes from the Pleistocene-Holocene transition to the end of the mid-Holocene, 11,700 - 4,2000 ybp; The second, named here Agricultural period ($t^1$), goes from the widespread aridification that marks the mid-Holocene transition (70) to 1500 AD and the arrival of European settlers in North America. The third, named here Euro-Industrial ($t^3$), extends to 1950 AD, corresponding to the establishment of modern cities, industrialized factories, mechanized crop harvesting, and modern road network, including the Interstate Highway System of North America (71, 72). The last time bin corresponds to modern times ($t^4$), after 1950 AD to 2022 AD. These four time-bins represent two gradients. From the Pleistocene to the present, the last deglaciation ($t^0$) experiences the highest climate fluctuations and the mid-Holocene aridification ($t^1$) experiences the next most, with little overall climate change in the final two ($t^2$ and $t^3$). While the second gradient, is associated with increasing human impact across the time-bins corresponding to the early expansion of early human settlements ($t^0$), increased land use by agrarian communities ($t^1$), European arrival and the beginning of the Industrial Revolution in North America ($t^2$), and finally modern capitalist society with highly mechanized crop harvesting, large cities and suburbs, and a complete interstate network ($t^3$). Deglaciation [between the Last Glacial Maximum (18,000 ybp), and early Holocene (11,700 ybp)] depicts the strongest mismatches between fossils occurrences and their associated environments because of time-averaging and age uncertainty (73). Because of these uncertainties and the scarce record of mammalian fossil before 11,700 ybp, our analysis is limited to the period between early Holocene and modern days.

*Climate estimates at fossil sites*

Mean annual temperature (MAT) and decadal mean total annual precipitation (MAP) are extracted for each mammal and plant occurrence from the CCSM3 debiased and downscaled paleoclimate model (74, 75, 76). For surface pollen samples and modern mammalian occurrences, we have used climate data from 1980 AD in the same CCSM3 paleoclimate model for data consistency. MAT and MAP were selected over other climate variables because MAT and MAP are strongly correlated ($R > 0.7$) with all other potential climate variables (see Wang et al. 2020 [53] supplementary information for details). Furthermore, the SynTrace CCSM3 precipitation and climate paleoseasonality (or min/max variable) reconstructions have been demonstrated to be less accurate than MAT or MAP (76). In brief, MAT and MAP were selected to reduce autocorrelation between variables, facilitate climatic niche interpretations, and minimize error in climate and climatic niche reconstruction.

*Climatic niches and time averaging*

The reconstruction of climatic niches through time in this study represent the realized climatic niche of taxa (77), a fraction of their fundamental climatic niche based on the available environment at any given time and after the resolutions of intra- and interspecific interactions (Hutchinson 1957). The climatic niches of individual taxa are reconstructed in two dimensions by extracting MAT and MAP values for each occurrence, based on average site age (**Fig. 1A**). Sites associated with erroneous climate estimations were discarded. For any given time-bins, all sites with a MAT range higher than 3°C and a MAP range higher than 25 mm were not considered (*see* Pineda-Munoz et al. 2021 [45]). Climatic niches are computed as the kernel-density function of MAT and MAP values based on *ecospat* package (31).

*Computing climate fidelity, available environment and subsampling*

Climate fidelity is the temporal stability of climatic niches, here computed as the temporal overlap of taxa's realized climatic niches. We compared the climatic niches of plants and mammalian taxa between all time-bins and the modern one (**Fig. 2B**), as well as between all time-bins and the next one (**Fig. S2**). The quality and quantity of sampling being highly variable through time, the computation of niche displacement or overlap (i.e., niche similarity or equivalency tests) only consider the available environment between the two analyzed periods (78). The tests for niche conservatism and the temporal overlap of climatic niches as Schoener's *D* index were computed with *ecospat* R package (32). Schoener's *D* index compute the overlap of two probability distributions (78, 79).

$$D(p_{t_n}, p_{t_{n+1}}) = 1 - \frac{1}{2}\sum_i |p_{t_n,i} - p_{t_{n+1},i}|$$

Where $p$ is the probability distribution of the occurrences of taxa ($i$), in the time-bin ($t_n$ or $t_{n+1}$). In this study, $p$ corresponds to the kernel-density of MAT and MAP of plant and mammal climatic niches for $t_n$ and $t_{n+1}$ time-bins. *D* ranges from 0 (no overlap) to 1 (identical niches). We used the function *ecospat.niche.overlap* from the ecospat package to compute temporal Schoener's *D* or climate fidelity (32). Because the number of occurrences can greatly vary from one time bins to another, climate fidelity is computed only in the light of available environment. To do so, we used the MAT/MAP values associated with all occurrences of plant or mammal of $t_n$ and $t_{n+1}$ time-bins to build the kernel-density of available environment. Secondly, in order to remove the potential climatic outlier or wrongly attributed occurrences, the overlaps are computed on 95 % of the kernel-density distribution. Finally, because our goal is to compare Schoener's D, or climate fidelity, of different taxa of plants and mammals through space and time, the figures and manuscript are based on median climate fidelity values and climate fidelity distributions of 1000 subsampling of 30 occurrences per taxa and time-bin. The sensitivity of climate fidelity to sample size is estimated by computing Schoener's *D* for all taxa and time-bins with bootstrap of *N* = 20, 30, 40, 50 (**Fig. S19**). As demonstrated before (29, 45), climate fidelity (and Scheoner's *D*) is sensitive to sample size, with a higher *D* when sample sizes increase, but the patterns illustrated and discussed in this study remain constant through varying sample size. We choose N = 30 as subsampling size in order to maximize both the number of included mammal (N = 40 remove 6 mammal taxa from the first time-bin) in our analysis and the relationship between plants and mammalian Schoener's *D* (**Fig. S3-S4**).

*Niche similarity and equivalency*

Given the available environment in the two compared time-bins (either with modern period or $t_0$ to $t_1$), we can use the *ecospat.niche.similarity.test* designed to evaluate the niche conservatism of modern sister taxa to evaluate the significance of climate fidelity (78, 80). The observed niche overlap, or

Schoener's *D*, computed between two time-bins will be compared to the null distribution of 1000 simulated niche overlaps. These simulated overlaps are based on simulated climatic niches build from random sampling of the available climates during $t_0$ and $t_1$. If the observed Schoener's *D* fall outside the envelope made of 950 of these simulations (i.e., 95% threshold), we can assume that this taxon exhibits significant niche conservatism interpreted here as significant climate fidelity. These plots comparing observed and simulated temporal overlap are illustrated in **Supplementary Fig. S20 to S39**. Because this study focuses on the quantification of the temporal change of overlap, niche volume and location in the environmental space as well as comparison of Schoener's *D* value between plants and mammals through time and space, the results of similarity or equivalency tests are not necessary to our interpretation. Especially because these two tests are intended to assess niche conservatism between species or through times (see Wang et al. 2023). The results of these tests were added to supplementary information if readers want to know more about niche conservatism through time for plant and mammal species in North America, see Pineda-Munoz et al. (2021) et Wang et al. (2023) for more details (29, 45).

*Sensitivity of climate fidelity to taxonomic level*

The ecological niche, and consequently, the climatic niche is a characteristic of species. In the NEOTOMA database, mammals are identified to the species level, however and as in most paleontological records, plant occurrences are based on pollen data and therefore identified to the genus level or above. To compare plant and mammal climate fidelity, we first have to evaluate how climate fidelity is affected by taxonomic level and how climate fidelity is preserved across taxonomic levels. To do so, we computed the climate fidelity of mammalian taxa at the genus and family levels and compare it with species-level climate fidelity. We combined mammalian species within genera and families and reproduce the exact procedure detailed above to compute climate fidelity (i.e., subsampling, bootstrapping, overlap in the light of available climate) with modern periods as well as between any time-bin and the next one. When we illustrate the relationship between mammalian climate fidelity at species and genus scale (**see Supplementary Fig. 3-4**), we found very strong correlation ($R^2$ = 0.93). The climate fidelity of mammalian species are preserved at the genus scale (**Supplementary Figs. 3-4**). Although we found that climate fidelity is less preserved at family scale, it is still relatively well-preserved. The climate fidelity of mammalian species and family depicted lower correlation than mammalian species and genus climate fidelity ($R^2$ = 0.90). Climate Fidelity is lower at family level ($D_{family}$ = 0.49 ± 0.1) than at species level ($D_{species}$ =0.50 ± 0.09); therefore the climate fidelity computed for the four plant families included in this study (Cupressaceae, Poaceae, Cyperaceae, Amaranthaceae) may be lower than the climate fidelity of the individual species that compose them. The direction of this effect (lower climate fidelity at higher taxonomic level) reinforces our findings (i.e., that mammals have lower climate fidelity than plants, plant climate fidelity is high, and the standard deviation of plants' climate fidelity is low).

*Traits and functional groups*

To assess the effect of traits and functional groups on plant and mammal climate fidelity, we sorted both datasets based on functional and morphological information extracted from the literature. The plant dataset has been divided in cold/wet and warm/dry taxa based on their niche centroids from the last 18,000 years. Following (29) Wang et al. (2023), *Tsuga*, *Abies*, *Alnus*, *Picea*, *Betula,* and *Fagus* form the cold/wet group, while *Fraxinus*, *Quercus*, *Pinus*, Cupressaceae, *Ulmus*, Cyperaceae, *Salix*, Poaceae, *Artemisia* and *Amaranthaceae* form the warm/arid group (**Fig. S40**). The plant dataset is divided based on their dispersion potential following Wang et al. (2023), as well, *Pinus*, *Quercus*, *Betula*, *Alnus*, *Fagus*, *Ulmus*, *Abies*, *Fraxinus* form the short-distance dispersal group, while *Picea*, *Tsuga*, Cupressaceae, *Salix*, Poaceae, Cyperaceae, *Artemisia* and Amaranthaceae form the long-distance dispersal group (29).

The mammalian dataset has been split following Schloss et al. 2012 into 6 body-size categories (0.1 – 1 kg, 1-5 kg, 5-50 kg, 50-100 kg, 100-1000 kg, > 1000 kg), 3 diets (herbivorous, carnivorous, omnivorous) and 4 velocity categories (< 3 km/y, 3 – 10 km/y, 10 – 20 km/y, > 20 km/y) (33). The two heaviest categories of body-size are merged for the figure depicted in the main manuscript, split in supplementary figures. Body-size is the best predictor for mammalian climate fidelity. Therefore, this morphological trait is included in the main part of the manuscript, while all others categorial results are detailed in supplementary information (**see Supplementary Figs. S5 to S9**). We have evaluated the difference in climate fidelity between these different groups with *t*-tests.

*Computing the overlap between the climates of North American landscapes with plant and mammal climatic niches*

To understand the effect of land use cover and specific landscapes on the history of climatic niches of North American mammals and plants, we have computed the overlap of the individual taxon niches with the climates of five common landscape (crops, cities, prairies, mountains and forests) through time. We obtained U.S. Geological Survey land cover data from 1970 because the average age of modern occurrence data is 1969 (81). We turn each land use cover polygon into one occurrence point, located in polygons' centroids. Land use categories were restricted to five by reclassifying complex nomenclature (e.g. cropland with grazing land became cropland) in our list of five landscapes (**Fig. S14, S15**) and removing land use cover categories that had fewer than 15 occurrences or that were irrelevant to this study (e.g., lakes) or too ambiguous (e.g., Forest and woodland mostly grazed).

Since the main climatic niche shifts happened after the arrival of Europeans, the overlap between pre-Europeans and post-Europeans climatic niches of mammal and plant taxa are compared to the five landscape's modern climates. Thanks to the **C**limate **E**xclusion **I**ndex (CEI), we could identify which taxa are facilitated by or extirpated from specific landscapes' climates after the arrival of Europeans and the industrialization of the continent. Finally, because we want to assess the increase of decrease of overlap between a landscape's climate and a taxon's niche, this index must not be obscured by trends coming from changing available climate through time. Consequently, the effect of climate change between pre- and post-Europeans periods has been integrated to the calculation of this index.

$$CEI = [D(N_L^{t_1}, N_T^{t_1}) - D(N_L^{t_1}, N_T^{t_2})] - \overline{D}(N_L^{t_1}, N_L^{t_2})$$

where $D(N_L^{t_1}, N_T^{t_1})$ is the overlap of modern landscape's climate ($N_L^{t_1}$) with one plant or one mammal climatic niche ($N_T^{t_1}$) after 1500 AD. $D(N_L^{t_1}, N_T^{t_2})$ is the overlap of modern landscape's climate with one plant or one mammal climatic niche ($N_T^{t_2}$) before 1500 AD. Based on these first two expressions in the bracket, the more negative the index, the more a taxon ($T$) niche shifted outside the landscape's ($L$) climate. The more positive the index, the more the taxon ($T$) has experienced a niche shift toward the landscape's ($L$) climate. The third expression of this equation $D(N_L^{t_1}, N_L^{t_2})$ is the correction for background climate change. All plants and mammals' occurrences points used to compute the overlap between modern landscape's climate and pre- or post-European climate niches are re-used to compute the overall change toward each habitat. For Urban areas for example, the mean of every $D(N_{Urban}^{t_1}, N_T^{t_1})$ and $D(N_{Urban}^{t_1}, N_T^{t_2})$ values computed between the Urban area's modern climate and every (mammal or plant) taxa climate niche is subtracted from the individual taxa's CEI value.

*Computing the geographical climate fidelity of plants and mammals' communities*

To identify the spatial patterns and distributions of climate fidelity, we have computed climate fidelity at community scale for both plants and mammalian communities. To do so, we used the individual climate fidelity of taxa computed on the first two time-bins (to minimize the effect of human impact) and computed the mean climate fidelity at community scale based on the composition of modern and fossil communities. Only sites with 5 taxa or more were considered as communities. For example, based on individual climate fidelity value depicted in Figure 2, a community composed of wolves (D = 0.57), racoons (D = 0.47), muskrats (D = 0.51), voles (D = 0.31), and squirrels (D = 0.35) would have an average climate fidelity of 0.44 ± 0.11. In the main part of the manuscript, we have only depicted the climate fidelity at community scale for modern plants and mammalian assemblages (**Fig. 5**). For the sake of clarity, we have divided the range of climate fidelity computed at community scale into four quartiles to highlight climate fidelity spatial patterns, with low-climate fidelity-community (0 – 25%), low-medium-climate fidelity-community (25% – 50%), medium-high-climate fidelity-community (50% – 75%) and finally high-climate fidelity-community (75% – 100%). The climate fidelity at community scale has been computed for the three previous time-bins but only illustrated in supplementary data (**see Fig. S17**). The comparison of mammalian and plants scale (**Fig. 5, Fig. S17**) confirm how plants are overall better at tracking climate than mammals and how plant's climate fidelity variance is lower than mammalian climate fidelity variance, as all plants communities in North America are part of the high-climate fidelity-community group defined for mammals.

*Comparing geographic patterns climate fidelity with land use and roughness indexes*

To estimate the effect of geographical features (like mountains or land use) on climate fidelity, computed at community scale, we have compared the spatial patterns of community-scale climate fidelity with *roughness* and *land use* patterns extracted respectively from Danielson & Gesch (2011) and Theobald et al. (2020) (58, 59). *Roughness* and *land use* values are extracted at each occurrence point corresponding to the mammal and plant communities. The correlation between *roughness*, *land use* and climate fidelity at the community scale were computed with function *modified.ttest* from R package SpatialPack (82) to correct for the effect of autocorrelation. Finally, the relationship between the distribution of community scale climate fidelity and latitude is computed for both mammal and plant communities during the four time-bins. These correlations are depicted in supplementary data (**Fig. S1**) for the all four time-bins, and the modern correlation (R and p-value) are illustrated in figure 5.

*Spatial and temporal autocorrelation*

In this study, we have reconstructed the climatic niches of plants and mammals during four time-bins by using their spatial distribution in association with paleoclimate estimates. Because all our methods are based on occurrence data, and the reconstruction of the temporal dynamic (stability or displacement) of climatic niches, we needed to consider temporal and spatial autocorrelation. In our analyses, temporal and spatial autocorrelations should reinforce the temporal stability of climatic niches, i.e., result in increased climate fidelity. Especially for the comparison of recent periods (i.e., Modern, Euro-industrial, Agricultural) that are shorter when combined than the oldest time-bins (i.e., End of Deglaciation). In this way, if there were temporal autocorrelation, it would make our finding of a strong niche displacement in mammals between Agricultural and Euro-Industrial time-bins (and stability during End of Deglaciation and Agricultural) more robust. Spatial autocorrelation acts in the same direction, but may be decreased by the strong subsampling used in all niche reconstructions.
