## Supplementary tables and figures for "Plants and mammals once tracked climate closely, then mammal niches shifted under human pressure"

**Supplementary Table 1**. Climate fidelity (Schoener's *D* ) computed for mammals between **Tx and Tx+1**. Schoener's *D* values depicted here are median values based on 1000 subsampled climatic niches and 1000 computed overlap.

| | Comparison of Tx to Tx+1 | | |
|---|---|---|---|
| Species | End of Deglaciation to Agricultural | Agricultural to Euro-Industrial | Euro-Industrial to Modern |
| Lepus californicus | 0,421 | 0,425 | 0,505 |
| Procyon lotor | 0,477 | 0,475 | 0,543 |
| Lynx rufus | 0,559 | 0,424 | 0,460 |
| Ursus americanus | 0,496 | 0,345 | 0,491 |
| Cervus elaphus | 0,573 | 0,378 | 0,529 |
| Urocyon cinereoargenteus | 0,419 | 0,377 | 0,528 |
| Odocoileus virginianus | 0,506 | 0,349 | 0,312 |
| Mephitis mephitis | 0,524 | 0,440 | 0,416 |
| Ondatra zibethicus | 0,516 | 0,459 | 0,595 |
| Castor canadensis | 0,578 | 0,370 | 0,576 |
| Marmota monax | 0,539 | 0,418 | 0,566 |
| Sigmodon hispidus | 0,290 | 0,401 | 0,482 |
| Eptesicus fuscus | 0,427 | 0,243 | 0,562 |
| Neotoma cinerea | 0,506 | 0,322 | 0,549 |
| Thomomys bottae | 0,562 | 0,210 | 0,587 |
| Antilocapra americana | 0,620 | 0,528 | 0,597 |
| Microtus ochrogaster | 0,313 | 0,366 | 0,616 |
| Onychomys leucogaster | 0,379 | 0,250 | 0,498 |
| Scalopus aquaticus | 0,562 | 0,417 | 0,600 |
| Mustela frenata | 0,555 | 0,425 | 0,591 |
| Vulpes vulpes | 0,590 | 0,347 | 0,512 |
| Bison bison | 0,541 | 0,517 | 0,481 |
| Sylvilagus floridanus | 0,437 | 0,527 | 0,518 |
| Sciurus carolinensis | 0,411 | 0,458 | 0,512 |
| Canis lupus | 0,570 | 0,281 | 0,539 |
| Tamias striatus | 0,556 | 0,404 | 0,580 |
| Blarina brevicauda | Low sampling | 0,377 | 0,587 |
| Sciurus niger | 0,349 | 0,469 | 0,541 |
| Puma concolor | Low sampling | 0,387 | 0,437 |
| Microtus pennsylvanicus | 0,490 | 0,369 | 0,591 |
| Taxidea taxus | 0,613 | 0,463 | 0,539 |
| Canis latrans | 0,547 | 0,418 | 0,420 |
| Geomys bursarius | 0,490 | 0,576 | 0,444 |
| Tamiasciurus hudsonicus | Low sampling | 0,337 | 0,634 |
| Thomomys talpoides | 0,486 | 0,431 | 0,638 |
| Lontra canadensis | Low sampling | 0,324 | 0,523 |
| Cynomys ludovicianus | Low sampling | 0,487 | 0,599 |
| Microtus montanus | Low sampling | 0,433 | 0,627 |

| | | | |
|---|---|---|---|
| Peromyscus maniculatus | 0,565 | 0,310 | 0,597 |
| Lepus townsendii | Low sampling | 0,577 | 0,548 |
| Ovis canadensis | 0,483 | 0,242 | 0,480 |
| Odocoileus hemionus | 0,601 | 0,327 | 0,497 |
| Sylvilagus audubonii | Low sampling | 0,281 | 0,547 |
| Dipodomys ordii | 0,463 | 0,389 | 0,533 |
| Sylvilagus nuttallii | 0,624 | 0,553 | 0,610 |
| **Average Schoener's D** | 0,504 | 0,398 | 0,536 |
| **Standard deviation's D** | 0,084 | 0,089 | 0,067 |
| **Global** | | | |
| Minimum D value | **0,210** | | |
| Maximum D value | **0,638** | | |
| **All average** | **0,478** | | |
| **Standard deviation** | **0,100** | | |

**Supplementary Table 2**. Climate fidelity (Schoener's *D* ) computed for plant genus between **Tx and Tx+1**. Schoener's *D* values depicted here are median values based on 1000 subsampled climatic niches and 1000 computed overlap.

| | Comparison of **Tx** to **Tx+1** | | |
|---|---|---|---|
| Plant taxa | End of Deglaciation to Agricultural | Agricultural to Euro-Industrial | Euro-Industrial to Modern |
| Pinus | 0,544 | 0,592 | 0,515 |
| Quercus | 0,556 | 0,610 | 0,544 |
| Picea | 0,619 | 0,556 | 0,542 |
| Betula | 0,556 | 0,464 | 0,447 |
| Alnus | 0,603 | 0,566 | 0,531 |
| Tsuga | 0,514 | 0,602 | 0,558 |
| Cupressaceae | 0,511 | 0,604 | 0,461 |
| Fagus | 0,603 | 0,623 | 0,523 |
| Ulmus | 0,567 | 0,600 | 0,587 |
| Abies | 0,541 | 0,574 | 0,573 |
| Fraxinus | 0,582 | 0,631 | 0,540 |
| Salix | 0,624 | 0,546 | 0,528 |
| Poaceae | 0,568 | 0,585 | 0,542 |
| Cyperaceae | 0,574 | 0,598 | 0,498 |
| Artemisia | 0,592 | 0,617 | 0,456 |
| Amaranthaceae | 0,494 | 0,590 | 0,440 |
| **Average Schoener's D** | **0,565** | **0,585** | **0,518** |
| **Standard deviation's D** | **0,038** | **0,040** | **0,045** |
| **Global** | | | |
| Minimum D value | **0,440** | | |
| Maximum D value | **0,631** | | |
| **All average** | **0,556** | | |
| **Standard deviation** | **0,049** | | |

**Supplementary Table 3**. Climate fidelity (Schoener's *D* ) computed for mammals between **Modern (t3) and $T_0$, $T_1$, $T_2$**. Schoener's *D* values depicted here are median values based on 1000 subsampled climatic niches and 1000 computed overlap.

| | Comparison of tx to **Modern** ($T_3$) | | |
|---|---|---|---|
| Species | End of Deglaciation to Modern | Agricultural to Modern | Euro-Industrial to Modern |
| Lepus californicus | 0,508 | 0,340 | 0,499 |
| Procyon lotor | 0,300 | 0,426 | 0,544 |
| Lynx rufus | 0,381 | 0,425 | 0,466 |
| Ursus americanus | 0,267 | 0,360 | 0,469 |
| Cervus elaphus | 0,399 | 0,432 | 0,517 |
| Urocyon cinereoargenteus | 0,330 | 0,332 | 0,531 |
| Odocoileus virginianus | 0,322 | 0,404 | 0,290 |
| Mephitis mephitis | 0,285 | 0,223 | 0,391 |
| Ondatra zibethicus | 0,454 | 0,500 | 0,586 |
| Castor canadensis | 0,434 | 0,425 | 0,582 |
| Marmota monax | 0,401 | 0,509 | 0,571 |
| Sigmodon hispidus | 0,206 | 0,325 | 0,467 |
| Eptesicus fuscus | 0,350 | 0,200 | 0,565 |
| Neotoma cinerea | 0,535 | 0,480 | 0,541 |
| Thomomys bottae | 0,374 | 0,284 | 0,583 |
| Antilocapra americana | 0,426 | 0,455 | 0,577 |
| Microtus ochrogaster | 0,169 | 0,424 | 0,623 |
| Onychomys leucogaster | 0,278 | 0,235 | 0,516 |
| Scalopus aquaticus | 0,343 | 0,371 | 0,609 |
| Mustela frenata | 0,373 | 0,494 | 0,585 |
| Vulpes vulpes | 0,437 | 0,378 | 0,508 |
| Bison bison | 0,370 | 0,359 | 0,471 |
| Sylvilagus floridanus | 0,318 | 0,414 | 0,515 |
| Sciurus carolinensis | 0,235 | 0,405 | 0,518 |
| Canis lupus | 0,276 | 0,264 | 0,521 |
| Tamias striatus | 0,490 | 0,492 | 0,578 |
| Blarina brevicauda | Low sampling | 0,401 | 0,583 |
| Sciurus niger | 0,098 | 0,308 | 0,532 |
| Puma concolor | Low sampling | 0,297 | 0,448 |
| Microtus pennsylvanicus | 0,443 | 0,401 | 0,608 |
| Taxidea taxus | 0,439 | 0,501 | 0,518 |
| Canis latrans | 0,308 | 0,241 | 0,427 |
| Geomys bursarius | 0,337 | 0,427 | 0,441 |
| Tamiasciurus hudsonicus | Low sampling | 0,362 | 0,606 |
| Thomomys talpoides | 0,526 | 0,468 | 0,630 |
| Lontra canadensis | Low sampling | 0,491 | 0,507 |
| Cynomys ludovicianus | Low sampling | 0,534 | 0,601 |
| Microtus montanus | Low sampling | 0,411 | 0,621 |
| Peromyscus maniculatus | Low sampling | 0,371 | 0,586 |
| Lepus townsendii | Low sampling | 0,522 | 0,538 |
| Ovis canadensis | 0,136 | 0,221 | 0,484 |
| Odocoileus hemionus | 0,365 | 0,448 | 0,501 |

| | | | |
|---|---|---|---|
| Sylvilagus audubonii | Low sampling | 0,416 | 0,525 |
| Dipodomys ordii | 0,356 | 0,321 | 0,528 |
| Sylvilagus nuttallii | 0,474 | 0,519 | 0,600 |
| Peromyscus leucopus | Low sampling | 0,275 | 0,481 |
| **Average Schoener's D** | **0,354** | **0,389** | **0,530** |
| **Standard deviation's D** | **0,105** | **0,091** | **0,068** |
| **Global** | | | |
| Minimum D value | **0,098** | | |
| Maximum D value | **0,630** | | |
| **Average All** | **0,430** | | |
| **Standard deviation** | **0,116** | | |

**Supplementary Table 4**. Climate fidelity (Schoener's *D* ) computed for plants between **modern and $T_0$, $T_1$ and $T_2$**. Schoener's *D* values depicted here are median values based on 1000 subsampled climatic niches and 1000 computed overlap.

| | Comparison of tx to **Modern** ($T_3$) | | |
|---|---|---|---|
| Plant taxa | End of Deglaciation to Modern | Agricultural to Modern | Euro-Industrial to Modern |
| Pinus | 0,5104 | 0,5237 | 0,4952 |
| Quercus | 0,4519 | 0,5468 | 0,5553 |
| Picea | 0,5338 | 0,5975 | 0,5356 |
| Betula | 0,5030 | 0,5615 | 0,4535 |
| Alnus | 0,4821 | 0,5879 | 0,5610 |
| Tsuga | 0,3295 | 0,5318 | 0,5819 |
| Cupressaceae | 0,3780 | 0,4663 | 0,4687 |
| Fagus | 0,4768 | 0,5186 | 0,5161 |
| Ulmus | 0,4493 | 0,5385 | 0,5851 |
| Abies | 0,4164 | 0,5055 | 0,5642 |
| Fraxinus | 0,4890 | 0,5318 | 0,5373 |
| Salix | 0,5661 | 0,5988 | 0,5339 |
| Poaceae | 0,4892 | 0,5208 | 0,5316 |
| Cyperaceae | 0,5350 | 0,5615 | 0,4905 |
| Artemisia | 0,4626 | 0,5043 | 0,4382 |
| Amaranthaceae | 0,4173 | 0,4642 | 0,4350 |
| **Average Schoener's D** | 0,4682 | 0,5350 | 0,5177 |
| **Standard deviation's D** | 0,0610 | 0,0404 | 0,0491 |
| **Global** | | | |
| Minimum D value | **0,3295** | | |
| Maximum D value | **0,5988** | | |
| **Average All** | **0,5069** | | |
| **Standard deviation** | **0,0574** | | |

**Supplementary Table 5**. Climate exclusion index (**CEI**) computed for mammals. It is calculated as the difference between the overlap of the modern (after 1950 AD) climatic niches of five landscapes and the mammals' climatic niches before and after the arrival of Europeans (1500 AD).

| SPECIES/HABITAT | **Mountain** | **Urban** | **Cropland** | **Grassland** | **Forest** |
|---|---|---|---|---|---|
| **Sciurus niger** | -0,06488 | 0,12174 | 0,19901 | 0,09653 | 0,1274 |
| **Scalopus aquaticus** | -0,09488 | 0,07174 | 0,11901 | 0,00653 | 0,0674 |
| **Peromyscus maniculatus** | -0,15488 | 0,11174 | 0,11901 | 0,09653 | 0,1574 |
| **Procyon lotor** | -0,10488 | 0,02174 | 0,10901 | 0,07653 | 0,1174 |
| **Thomomys talpoides** | 0,03512 | 0,08174 | 0,10901 | 0,07653 | 0,0474 |
| **Ovis canadensis** | -0,19488 | 0,09174 | 0,10901 | 0,05653 | 0,1074 |
| **Marmota monax** | 0,04512 | 0,08174 | 0,09901 | 0,07653 | 0,0774 |
| **Sigmodon hispidus** | -0,09488 | 0,09174 | 0,09901 | -0,00347 | 0,0574 |
| **Thomomys bottae** | -0,22488 | 0,24174 | 0,09901 | 0,02653 | 0,0974 |
| **Mustela frenata** | -0,16488 | 0,17174 | 0,09901 | 0,01653 | 0,1174 |
| **Sylvilagus floridanus** | -0,06488 | 0,07174 | 0,09901 | 0,06653 | 0,0974 |
| **Odocoileus hemionus** | -0,24488 | 0,15174 | 0,09901 | 0,02653 | 0,1474 |
| **Sylvilagus audubonii** | -0,15488 | 0,14174 | 0,07901 | 0,03653 | 0,0774 |
| **Neotoma cinerea** | -0,00488 | 0,08174 | 0,06901 | 0,07653 | 0,0974 |
| **Microtus montanus** | 0,06512 | 0,05174 | 0,06901 | 0,05653 | 0,0274 |
| **Peromyscus leucopus** | -0,18488 | -0,03826 | 0,06901 | 0,01653 | 0,1074 |
| **Sciurus carolinensis** | -0,08488 | 0,03174 | 0,05901 | 0,00653 | 0,0474 |
| **Sylvilagus nuttallii** | -0,05488 | 0,06174 | 0,05901 | 0,01653 | 0,0274 |
| **Urocyon cinereoargenteus** | -0,15488 | 0,03174 | 0,04901 | 0,02653 | 0,0974 |
| **Tamiasciurus hudsonicus** | 0,09512 | -0,06826 | 0,04901 | 0,12653 | 0,0974 |
| **Geomys bursarius** | -0,11488 | -0,00826 | 0,03901 | 0,05653 | 0,0174 |
| **Tamias striatus** | 0,04512 | 0,02174 | 0,02901 | 0,05653 | 0,0374 |
| **Mephitis mephitis** | -0,19488 | -0,04826 | 0,01901 | 0,02653 | 0,0474 |
| **Canis latrans** | -0,18488 | 0,04174 | 0,01901 | -0,02347 | 0,1074 |
| **Lepus townsendii** | -0,07488 | 0,03174 | 0,01901 | -0,00347 | -0,0126 |
| **Puma concolor** | -0,07488 | -0,10826 | 0,00901 | 0,06653 | 0,0074 |
| **Taxidea taxus** | -0,15488 | 0,04174 | 0,00901 | 0,04653 | 0,0374 |
| **Dipodomys ordii** | -0,21488 | 0,05174 | 0,00901 | 0,00653 | -0,0126 |
| **Lepus californicus** | -0,18488 | 0,04174 | -0,00099 | -0,01347 | -0,0226 |
| **Cervus elaphus** | 0,08512 | -0,15826 | -0,00099 | 0,15653 | 0,0574 |
| **Ondatra zibethicus** | -0,01488 | -0,11826 | -0,00099 | 0,06653 | 0,0174 |
| **Microtus pennsylvanicus** | 0,06512 | -0,08826 | -0,00099 | 0,09653 | 0,0274 |
| **Canis lupus** | -0,09488 | -0,05826 | -0,01099 | 0,10653 | 0,1074 |
| **Ursus americanus** | 0,11512 | -0,17826 | -0,02099 | 0,16653 | 0,0674 |
| **Blarina brevicauda** | -0,00488 | -0,06826 | -0,02099 | 0,01653 | -0,0126 |
| **Vulpes vulpes** | -0,11488 | -0,00826 | -0,03099 | 0,03653 | 0,0474 |
| **Cynomys ludovicianus** | -0,12488 | -0,03826 | -0,03099 | -0,03347 | -0,0626 |
| **Castor canadensis** | 0,06512 | -0,14826 | -0,04099 | 0,10653 | 0,0574 |
| **Lynx rufus** | -0,00488 | -0,06826 | -0,05099 | 0,03653 | -0,0026 |
| **Odocoileus virginianus** | -0,09488 | -0,11826 | -0,05099 | 0,00653 | -0,0326 |
| **Onychomys leucogaster** | -0,19488 | -0,01826 | -0,05099 | -0,00347 | -0,0126 |
| **Microtus ochrogaster** | -0,27488 | -0,03826 | -0,06099 | -0,12347 | -0,1026 |
| **Antilocapra americana** | -0,12488 | -0,07826 | -0,07099 | -0,05347 | -0,0626 |
| **Lontra canadensis** | 0,02512 | -0,16826 | -0,08099 | 0,04653 | -0,0326 |

| | | | | | |
|---|---|---|---|---|---|
| **Bison bison** | 0,04512 | -0,19826 | -0,10099 | -0,00347 | -0,0526 |
| **Minimum** | -0,2749 | -0,1983 | -0,1010 | -0,1235 | -0,1026 |
| **Maximum** | 0,11512 | 0,24174 | 0,19901 | 0,16653 | 0,1574 |

**Supplementary Table 6**. Climate exclusion index (**CEI**) computed for mammals. It is calculated as the difference between the overlap of the modern (after 1950 AD) climatic niches of five landscapes and the plants' climatic niches before and after the arrival of Europeans (1500 AD).

| GENUS/HABITAT | Mountain | Urban | Cropland | Grassland | Forest |
|---|---|---|---|---|---|
| **Quercus** | -0,07 | 0,00 | 0,01 | 0,08 | -0,07 |
| **Cupressaceae** | -0,15 | -0,01 | -0,02 | 0,05 | -0,05 |
| **Pinus** | -0,06 | -0,05 | -0,04 | 0,13 | -0,03 |
| **Artemisia** | -0,23 | -0,08 | -0,04 | 0,11 | -0,05 |
| **Fraxinus** | -0,04 | -0,04 | -0,04 | 0,07 | -0,07 |
| **Amaranthaceae** | -0,20 | -0,06 | -0,05 | 0,05 | -0,05 |
| **Poaceae** | -0,12 | -0,07 | -0,05 | 0,06 | -0,06 |
| **Abies** | -0,04 | -0,05 | -0,07 | 0,06 | -0,08 |
| **Salix** | -0,09 | -0,06 | -0,09 | 0,08 | -0,10 |
| **Betula** | -0,02 | -0,07 | -0,10 | 0,09 | -0,12 |
| **Alnus** | -0,01 | -0,09 | -0,10 | 0,12 | -0,10 |
| **Ulmus** | -0,16 | -0,02 | -0,11 | 0,00 | -0,12 |
| **Tsuga** | 0,01 | -0,06 | -0,11 | 0,06 | -0,13 |
| **Picea** | -0,03 | -0,13 | -0,16 | 0,10 | -0,16 |
| **Fagus** | -0,07 | -0,06 | -0,16 | 0,04 | -0,14 |
| **Cyperaceae** | -0,08 | -0,15 | -0,17 | 0,13 | -0,10 |
| **Minimum** | -0,23 | -0,15 | -0,17 | 0,00 | -0,16 |
| **Maximum** | 0,01 | 0,00 | 0,01 | 0,13 | -0,03 |

**Supplementary Table 7**. Climate fidelity (Schoener's *D* ) computed for mammals between the End of Deglaciation (see Fig. 2B or Table S1) and the Modern periods compared to climate fidelity score obtained between the End of Deglaciation and the Agricultural periods (see Fig. 2A and Table S3). This table shows the intensity of change related to the arrival of Europeans and the anthropisation of North America

| Species | End of Deglaciation to Agricultural (Table S1) | End of Deglaciation to Modern (Table S3) | Intensity of change |
|---|---|---|---|
| Lepus californicus | 0,421 | 0,508 | -0,087 |
| Procyon lotor | 0,477 | 0,300 | 0,177 |
| Lynx rufus | 0,559 | 0,381 | 0,178 |
| Ursus americanus | 0,496 | 0,267 | 0,229 |
| Cervus elaphus | 0,573 | 0,399 | 0,175 |
| Urocyon cinereoargenteus | 0,419 | 0,330 | 0,089 |
| Odocoileus virginianus | 0,506 | 0,322 | 0,185 |
| Mephitis mephitis | 0,524 | 0,285 | 0,238 |
| Ondatra zibethicus | 0,516 | 0,454 | 0,062 |
| Castor canadensis | 0,578 | 0,434 | 0,144 |
| Marmota monax | 0,539 | 0,401 | 0,138 |
| Sigmodon hispidus | 0,290 | 0,206 | 0,084 |
| Eptesicus fuscus | 0,427 | 0,350 | 0,076 |
| Neotoma cinerea | 0,506 | 0,535 | -0,029 |
| Thomomys bottae | 0,562 | 0,374 | 0,188 |
| Antilocapra americana | 0,620 | 0,426 | 0,194 |
| Microtus ochrogaster | 0,313 | 0,169 | 0,144 |
| Onychomys leucogaster | 0,379 | 0,278 | 0,101 |
| Scalopus aquaticus | 0,562 | 0,343 | 0,219 |
| Mustela frenata | 0,555 | 0,373 | 0,181 |
| Vulpes vulpes | 0,590 | 0,437 | 0,152 |
| Bison bison | 0,541 | 0,370 | 0,171 |
| Sylvilagus floridanus | 0,437 | 0,318 | 0,119 |
| Sciurus carolinensis | 0,411 | 0,235 | 0,176 |
| Canis lupus | 0,570 | 0,276 | 0,294 |
| Tamias striatus | 0,556 | 0,490 | 0,066 |
| Blarina brevicauda | Low sampling | Low sampling | Low sampling |
| Sciurus niger | 0,349 | 0,098 | 0,252 |
| Puma concolor | Low sampling | Low sampling | Low sampling |
| Microtus pennsylvanicus | 0,490 | 0,443 | 0,047 |
| Taxidea taxus | 0,613 | 0,439 | 0,174 |
| Canis latrans | 0,547 | 0,308 | 0,239 |
| Geomys bursarius | 0,490 | 0,337 | 0,154 |
| Tamiasciurus hudsonicus | Low sampling | Low sampling | Low sampling |
| Thomomys talpoides | 0,486 | 0,526 | -0,040 |
| Lontra canadensis | Low sampling | Low sampling | Low sampling |
| Cynomys ludovicianus | Low sampling | Low sampling | Low sampling |
| Microtus montanus | Low sampling | Low sampling | Low sampling |
| Peromyscus maniculatus | 0,565 | Low sampling | Low sampling |
| Lepus townsendii | Low sampling | Low sampling | Low sampling |
| Ovis canadensis | 0,483 | 0,136 | 0,347 |
| Odocoileus hemionus | 0,601 | 0,365 | 0,236 |
| Sylvilagus audubonii | Low sampling | Low sampling | Low sampling |

| | | | |
|---|---|---|---|
| Dipodomys ordii | 0,463 | 0,356 | 0,108 |
| Sylvilagus nuttallii | 0,624 | 0,474 | 0,149 |
| **Average Schoener's D** | **0,504** | **0,354** | **0,148** |
| **Standard deviation's D** | **0,084** | **0,105** | **0,090** |

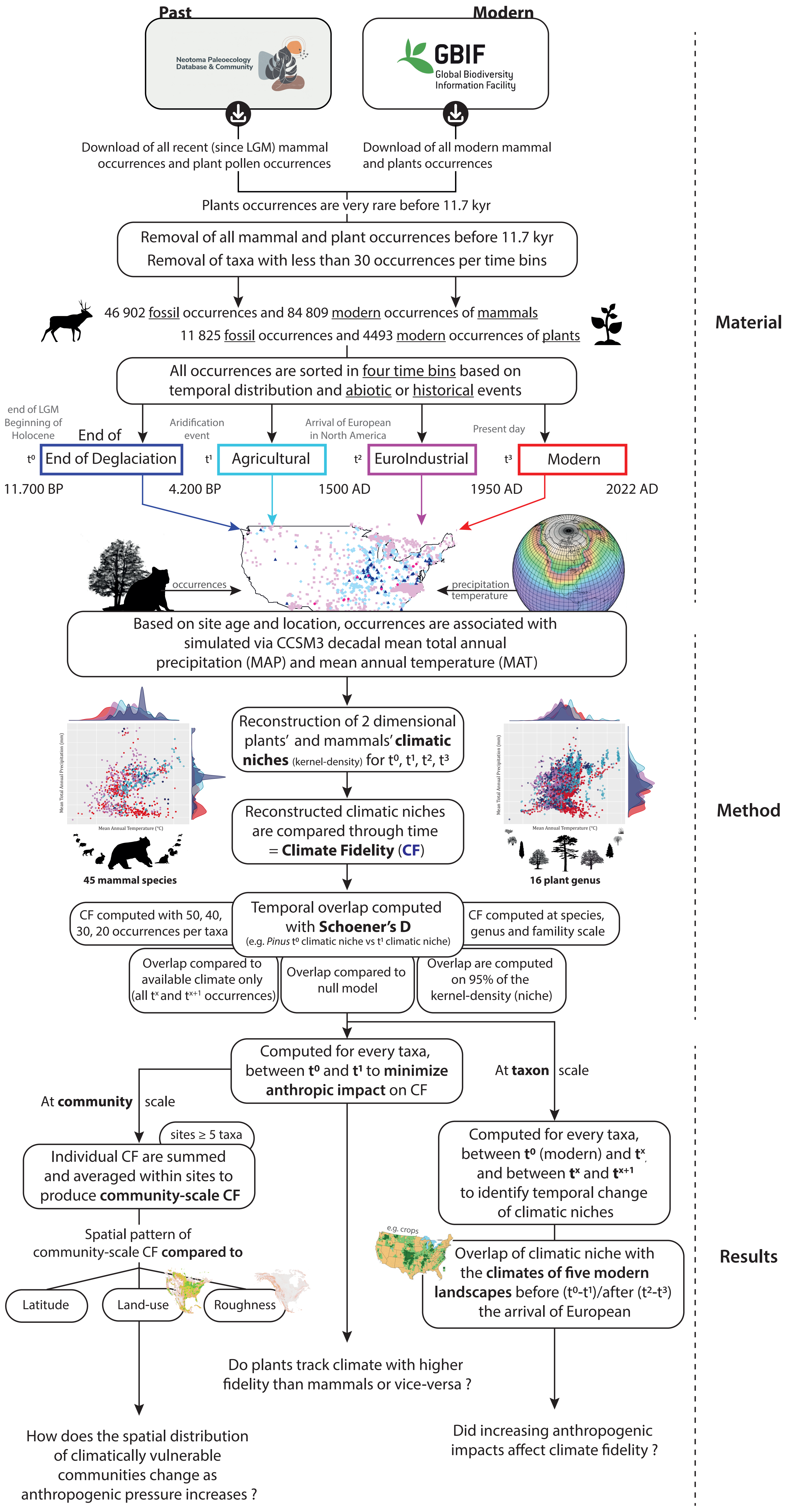

**Supplementary Figure 1**. Study workflow. Datasets of plant and mammal occurrences have been cleaned and analysed in identical ways. Final arrows point toward the three main hypothesis tested in this study.

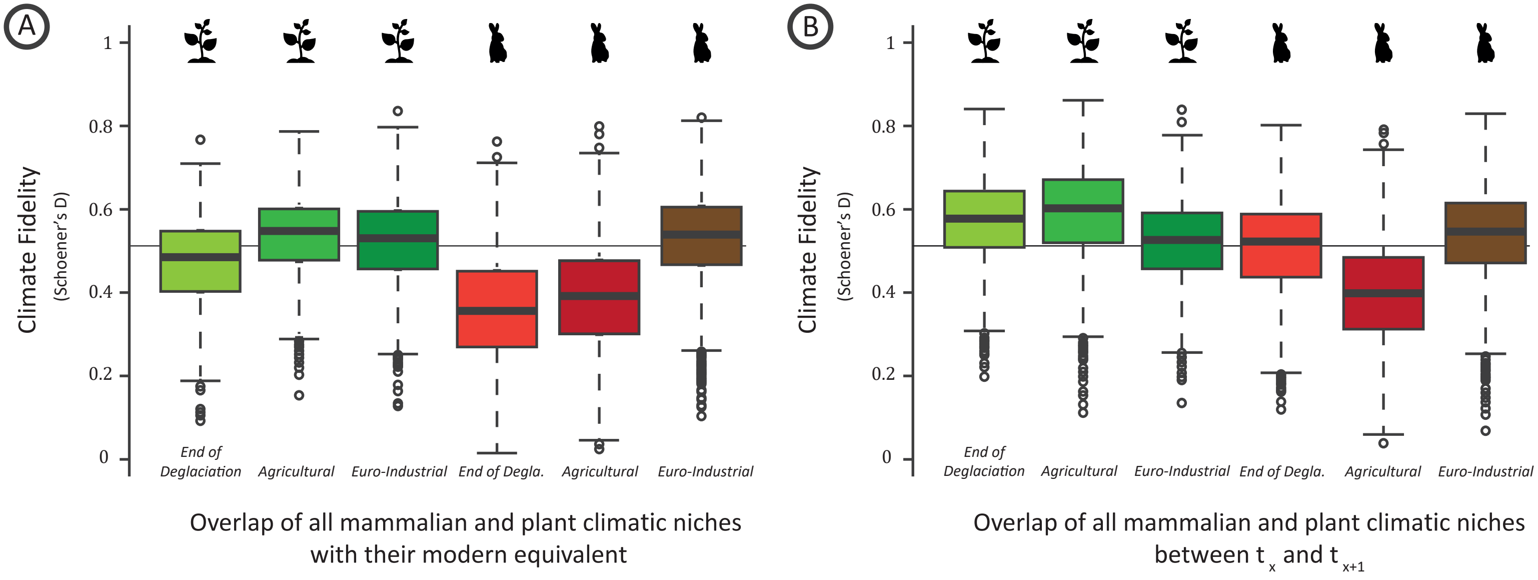

**Supplementary Figure 2**. A) Distribution of climate fidelity of mammals and plants computed from each time period to the Modern period. B) Distribution of climate fidelity of mammals and plants computed from each time period ($t^x$) to the next period ($t^{x+1}$).

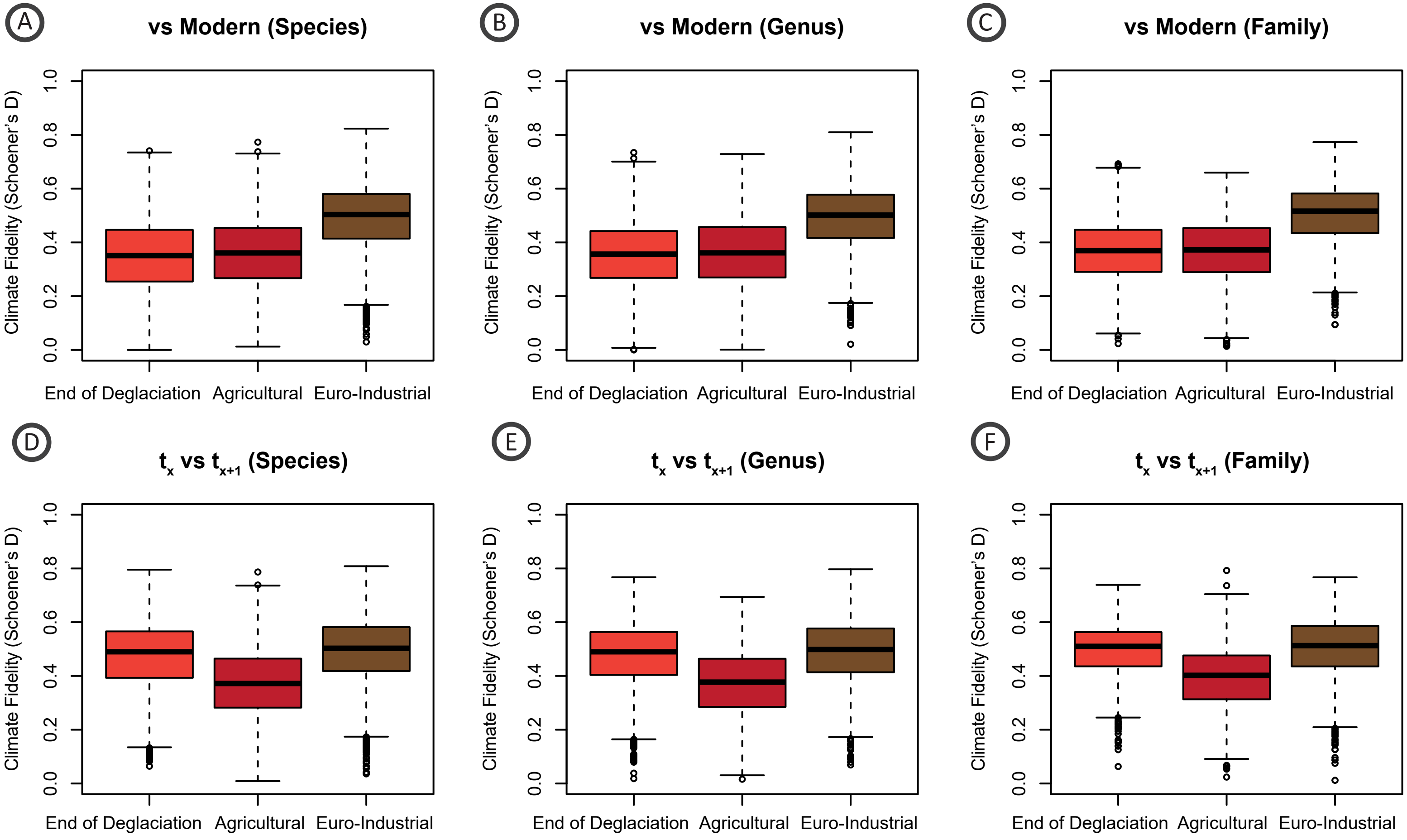

**Supplementary Figure 3**. Distribution of climate fidelity for all mammals species (A, D), genera (B, E) and families (C, F). Top row is climate fidelity computed between modern climatic niches and previous ones ($t_x$); bottom row boxplots depict climate fidelity computed between $t_x$ and $t_{x+1}$.

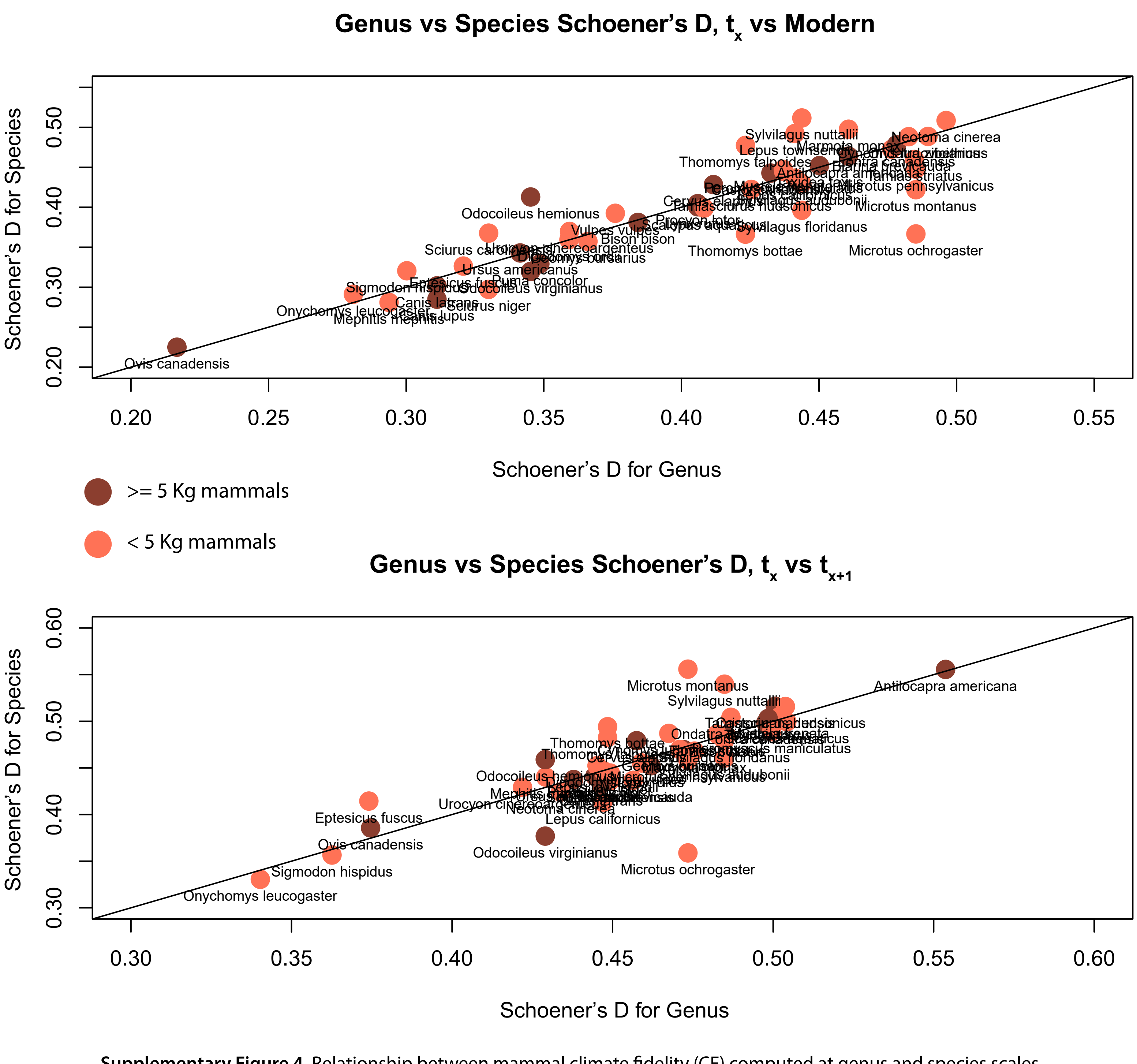

**Supplementary Figure 4**. Relationship between mammal climate fidelity (CF) computed at genus and species scales. Top plot illustrates CF computed between $t_x$ and $t_3$ (modern period). Bottom top illustrates CF computed between $t_x$ and $t_{x+1}$.

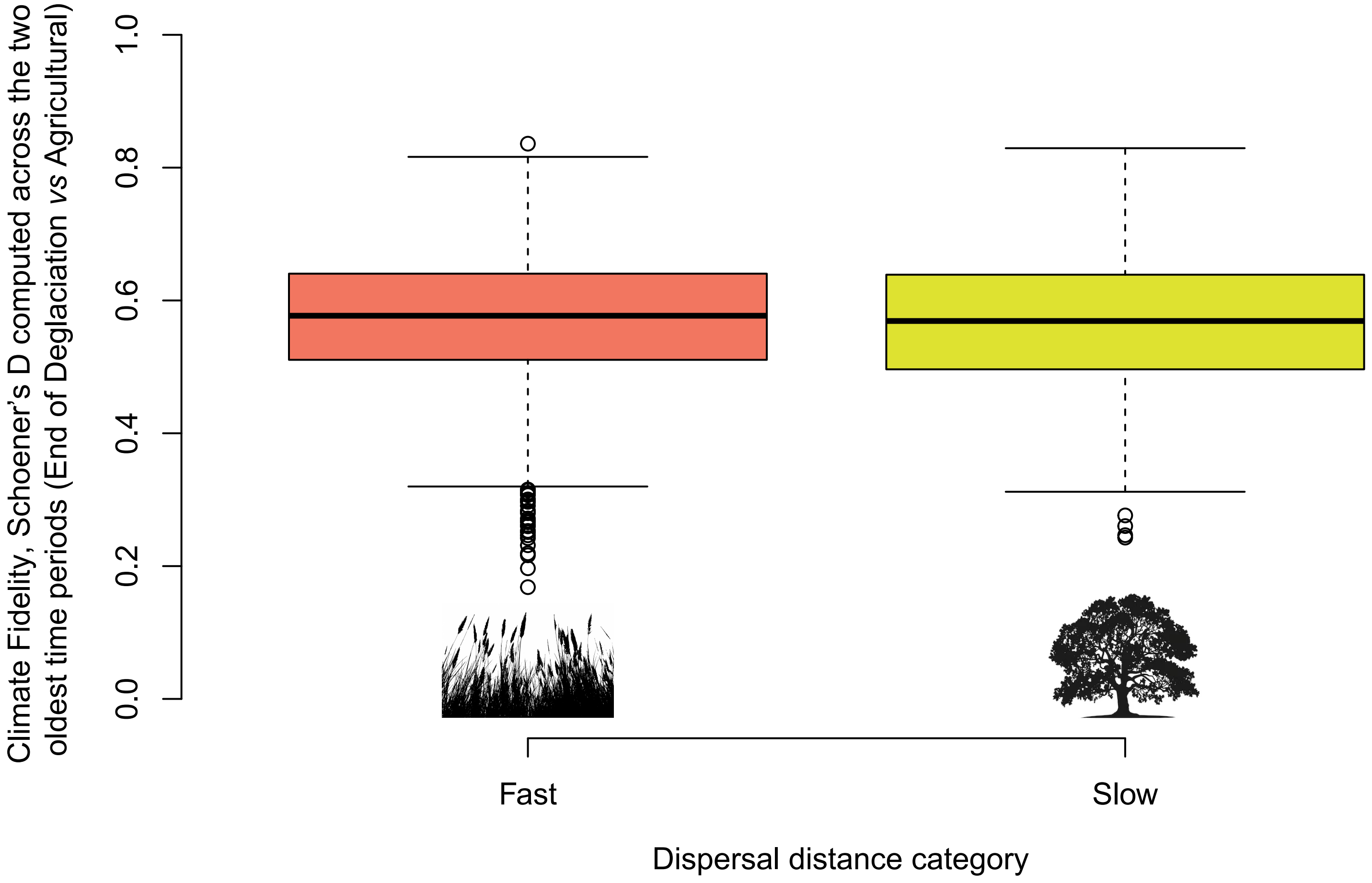

**Supplementary Figure 5**. Distribution of fast and slow moving plant climate fidelity computed on deglaciation and agricultural periods. Plant dataset is divided based on their dispersion potential following Wang et al. (2023). *Pinus*, *Quercus*, *Betula*, *Alnus*, *Fagus*, *Ulmus*, *Abies*, *Fraxinus* form the short-distance dispersal group, while *Picea*, *Tsuga*, Cupressaceae, *Salix*, Poaceae, Cyperaceae, *Artemisia* and Amaranthaceae form the long-distance dispersal group.

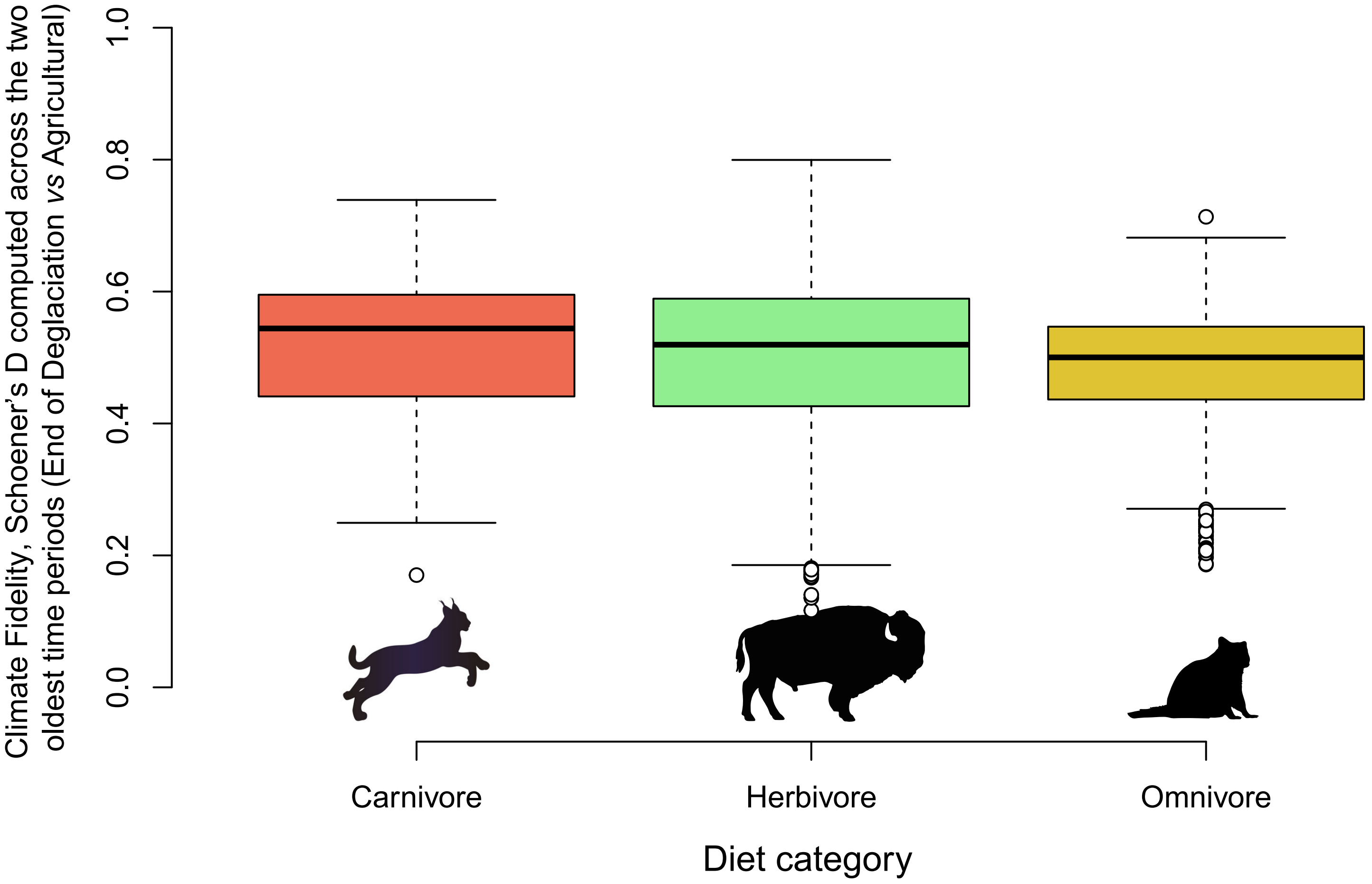

**Supplementary Figure 6.** Distribution of mammal climate fidelity computed on End of Deglaciation and Agricultural periods as a function of diet category.

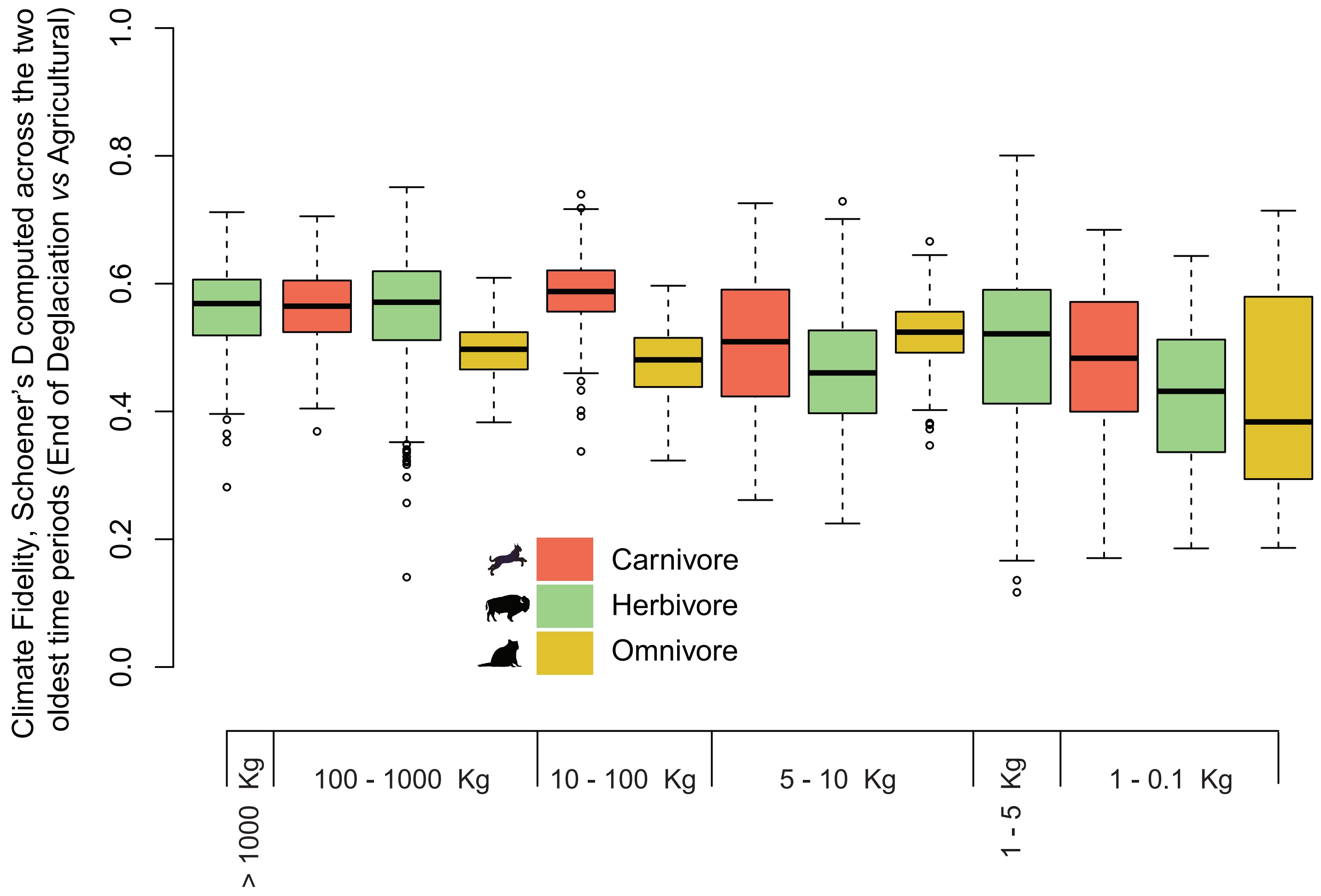

**Supplementary Figure 7.** Distribution of mammal climate fidelity computed on End of Deglaciation and Agricultural periods as a function of body-size (Schloss et al. 2012) and diet.

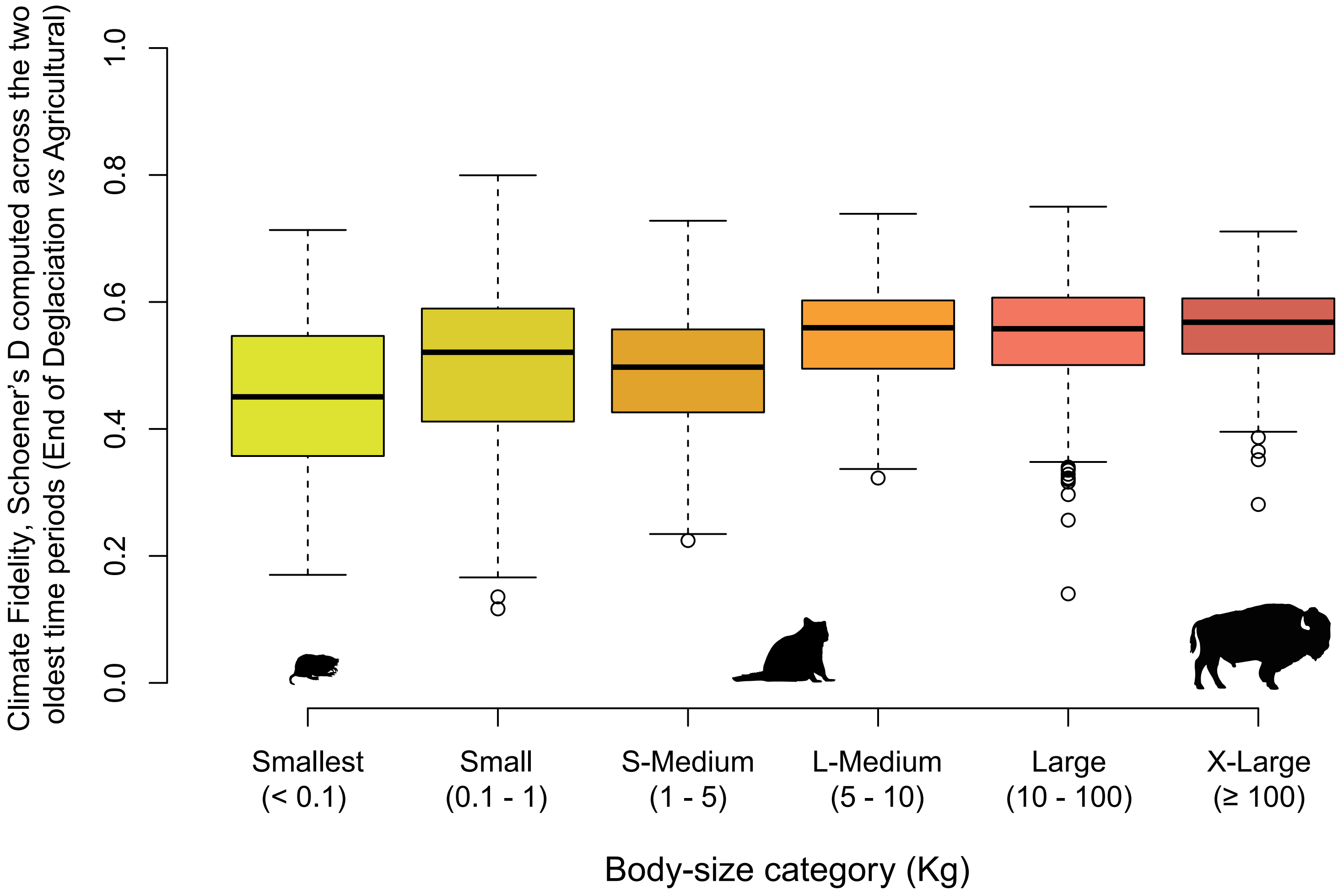

**Supplementary Figure 8.** Distribution of mammal climate fidelity computed on the End of Deglaciation and Agricultural periods as a function of body-size category following Schloss et al. (2012).

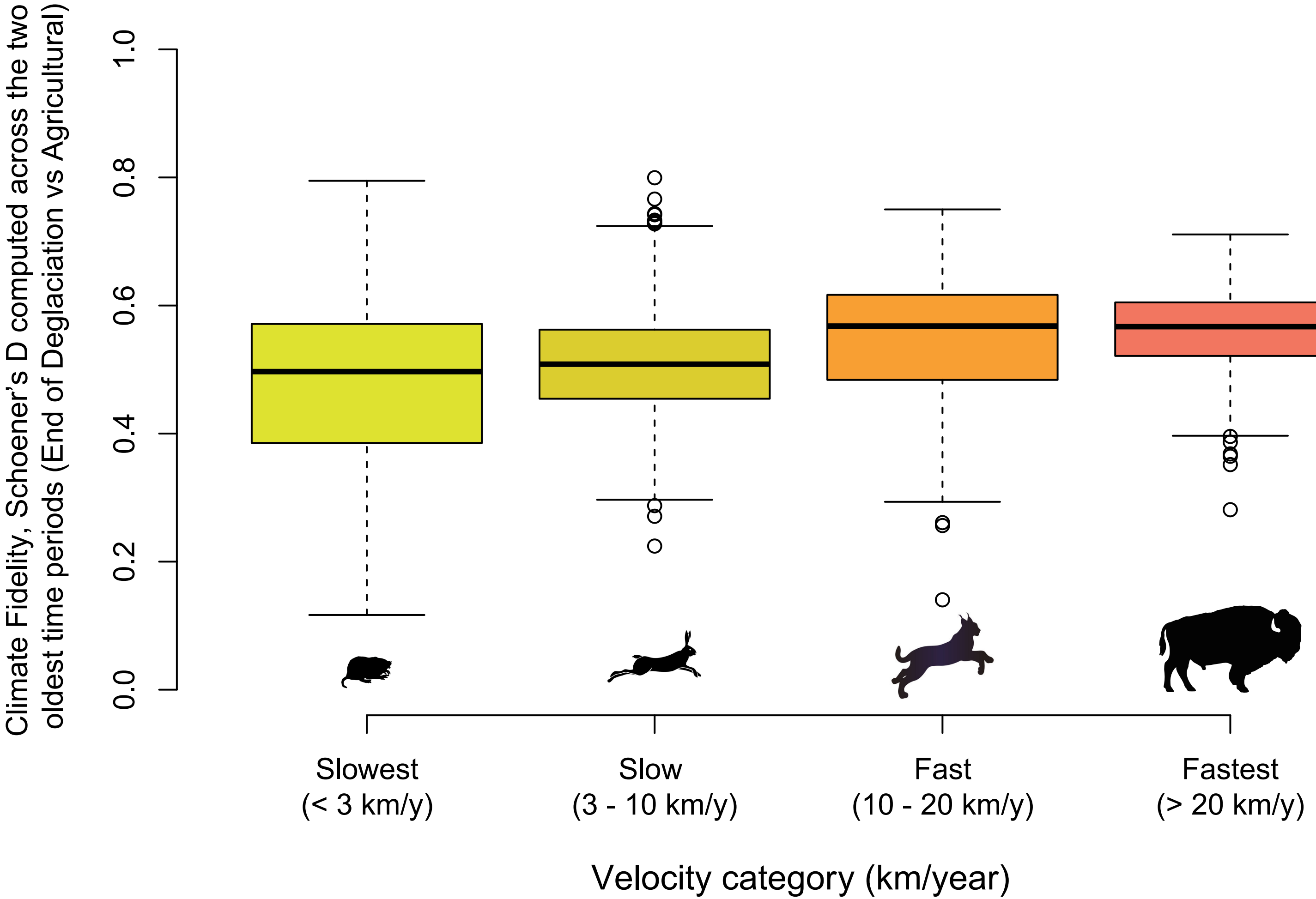

**Supplementary Figure 9**. Distribution of mammal climate fidelity computed on the End of Deglaciation and Agricultural periods as a function of velocity category following Schloss et al. (2012).

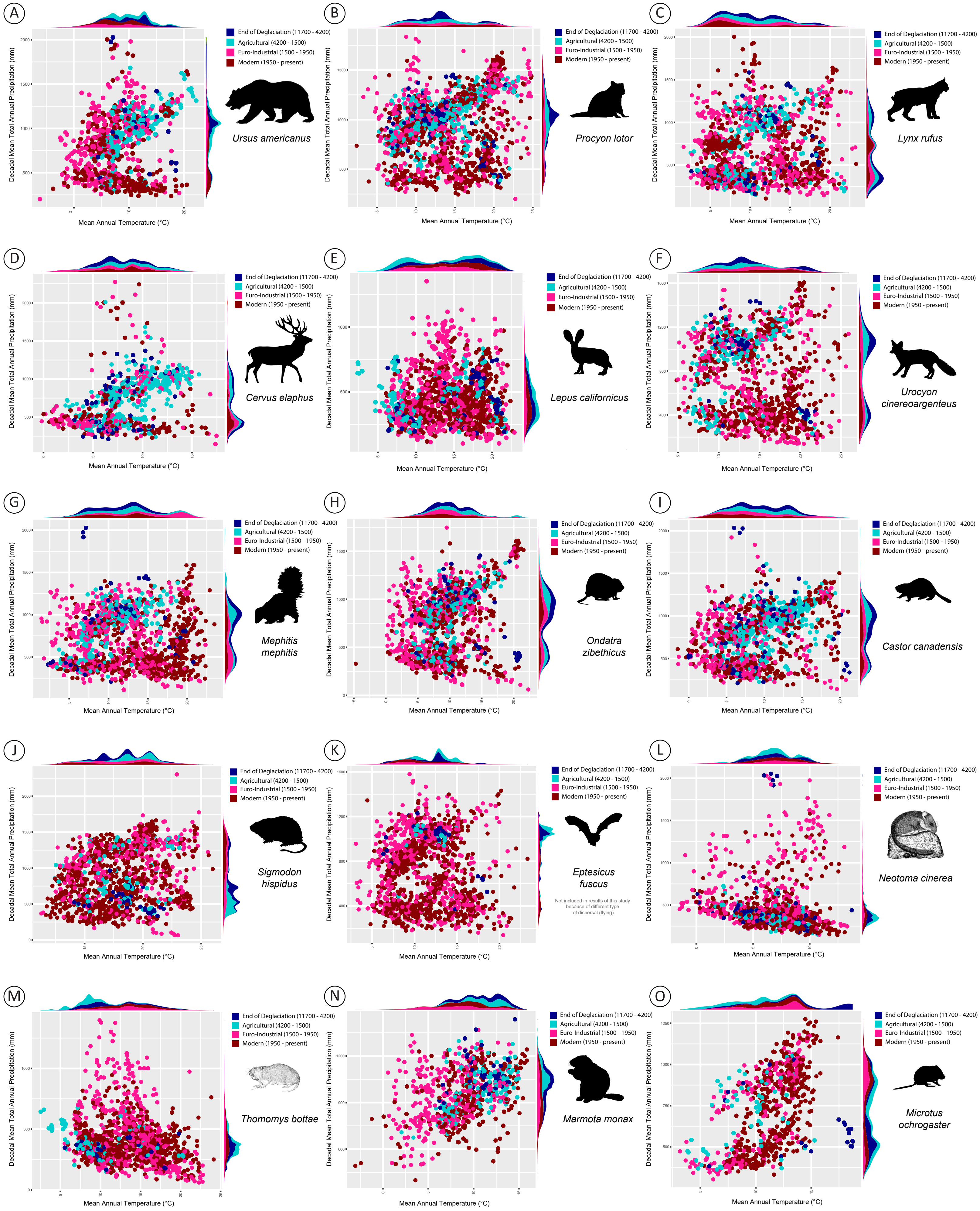

**Supplementary Figure 10.** Climatic niches of 15 mammal species. Every dot is an occurrence associated to a simulated value of decadal mean total annual precipitation (MAP) and mean annual temperature (MAT). Temporal overlap (climate fidelity) of niches are computed with 95% kernel-density.

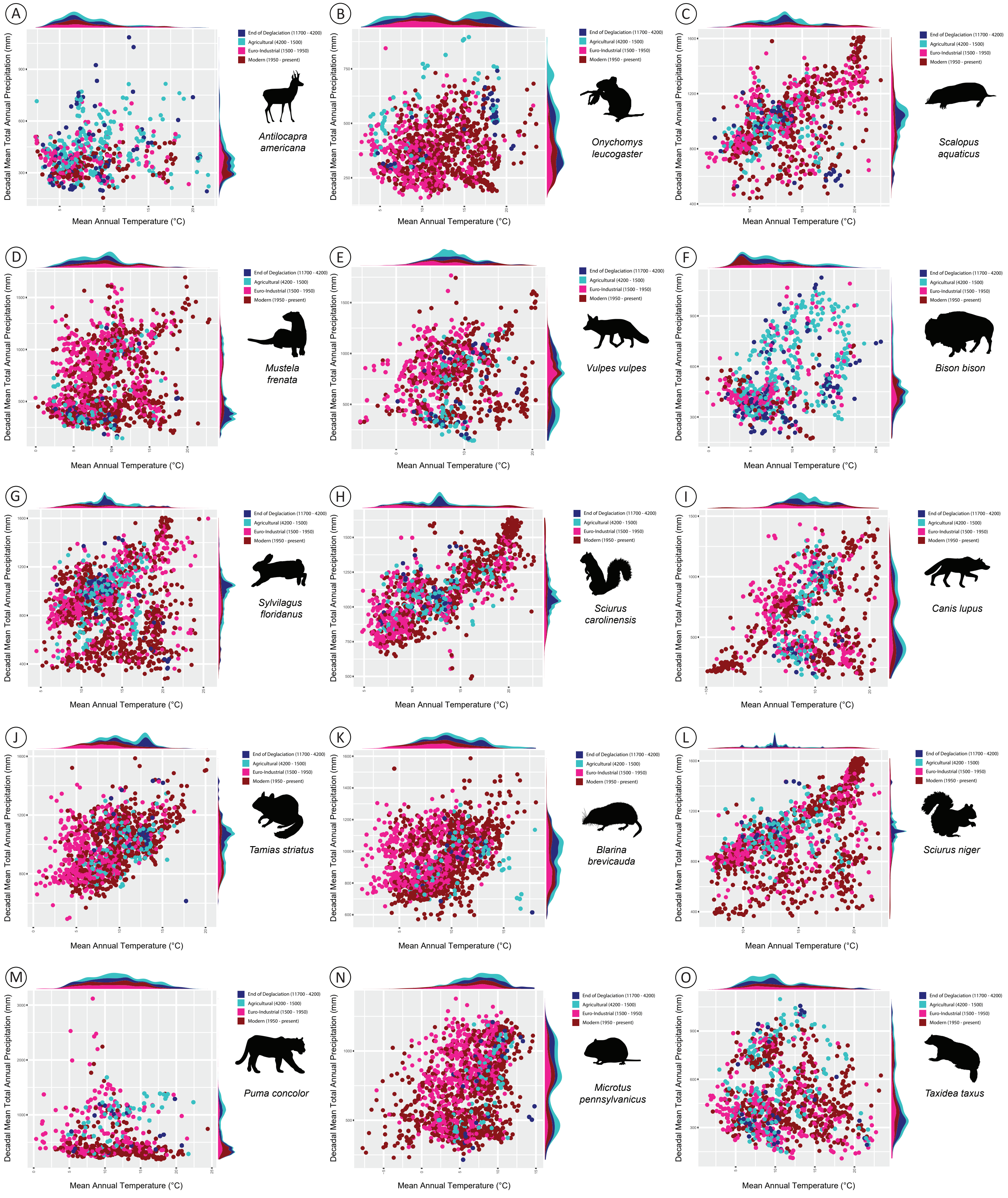

**Supplementary Figure 11.** Climatic niches of 15 mammal species. Every dot is an occurrence associated to a simulated value of decadal mean total annual precipitation (MAP) and mean annual temperature (MAT). Temporal overlap (climate fidelity) of niches are computed with 95% kernel-density.

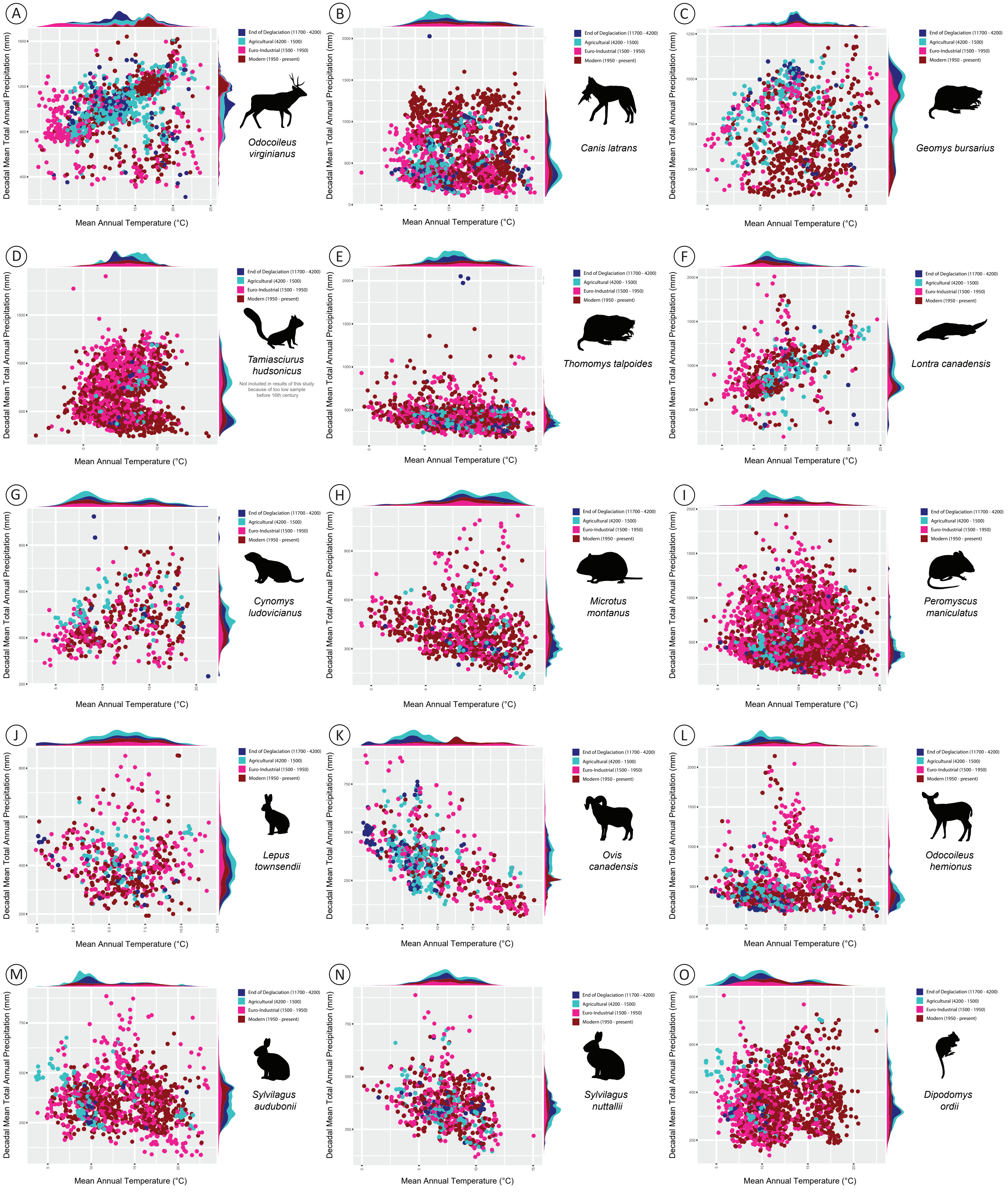

**Supplementary Figure 12.** Climatic niches of 15 mammal species. Every dot is an occurrence associated to a simulated value of decadal mean total annual precipitation (MAP) and mean annual temperature (MAT). Temporal overlap (climate fidelity) of niches are computed with 95% kernel-density.

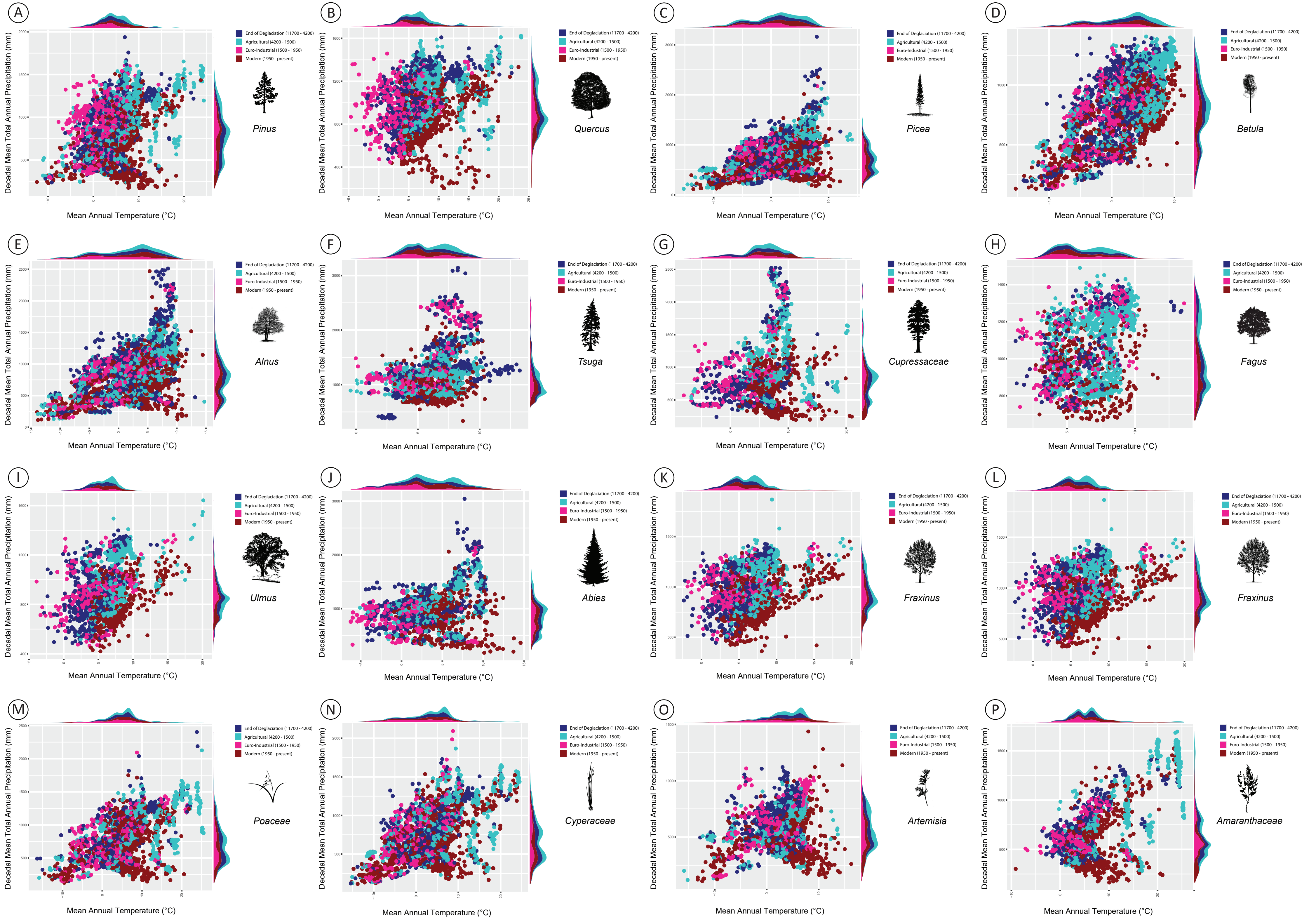

**Supplementary Figure 13.** Climatic niches of 16 plant taxa. Every dot is an occurrence associated to a simulated value of decadal mean total annual precipitation (MAP) and mean annual temperature (MAT). Temporal overlap (climate fidelity) of niches are computed with 95% kernel-density.

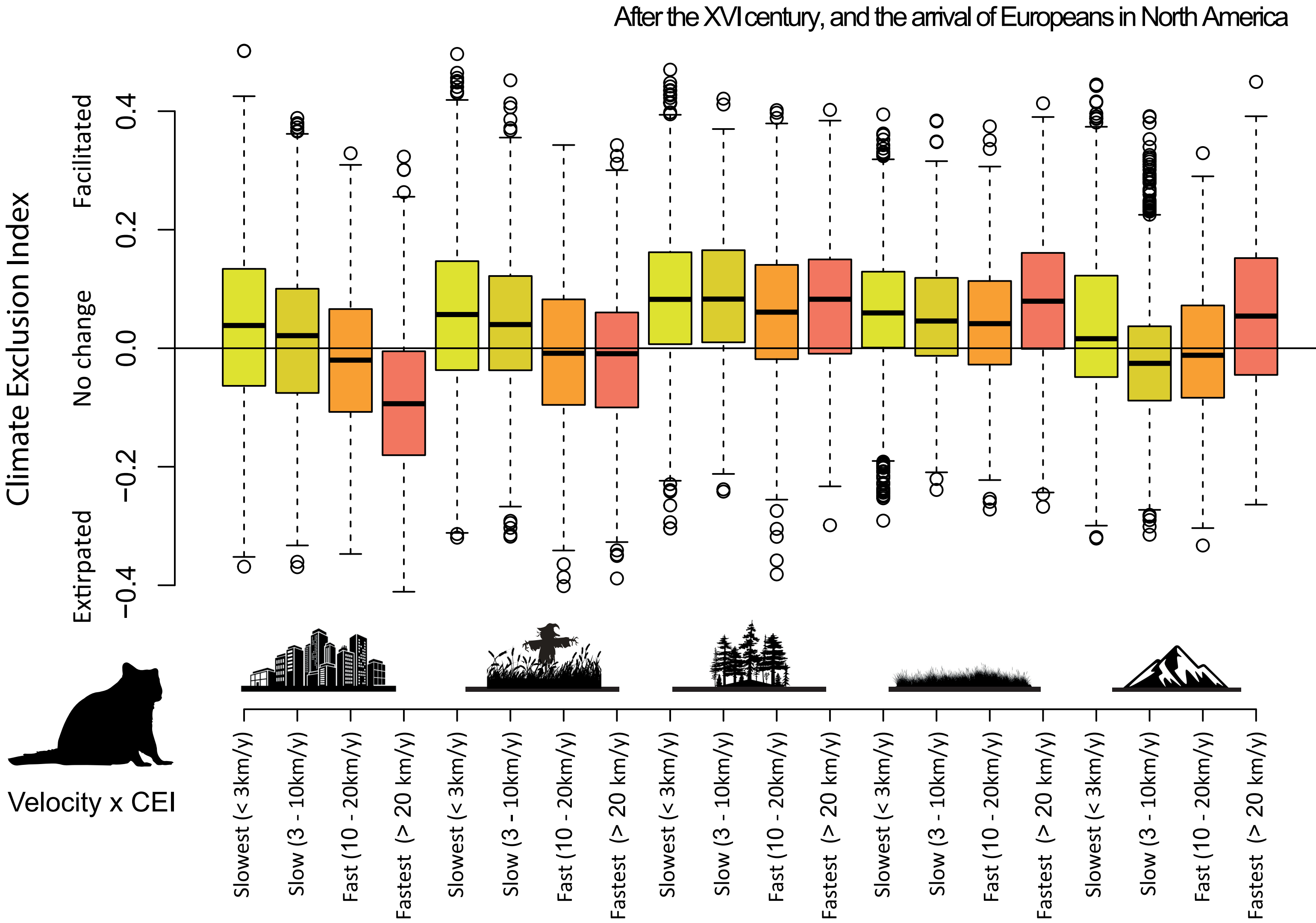

**Supplementary Figure 14.** Distribution of mammal Climate Exclusion Index (CEI) in function of velocity category (Schloss et al. 2012). A positive CEI shows that mammal climatic niches is overlapping more with landscape's climate after 1500 AD. Negative CEI shows extirpation. From left to right: Urban, Cropland, Forest, Grassland, Mountain

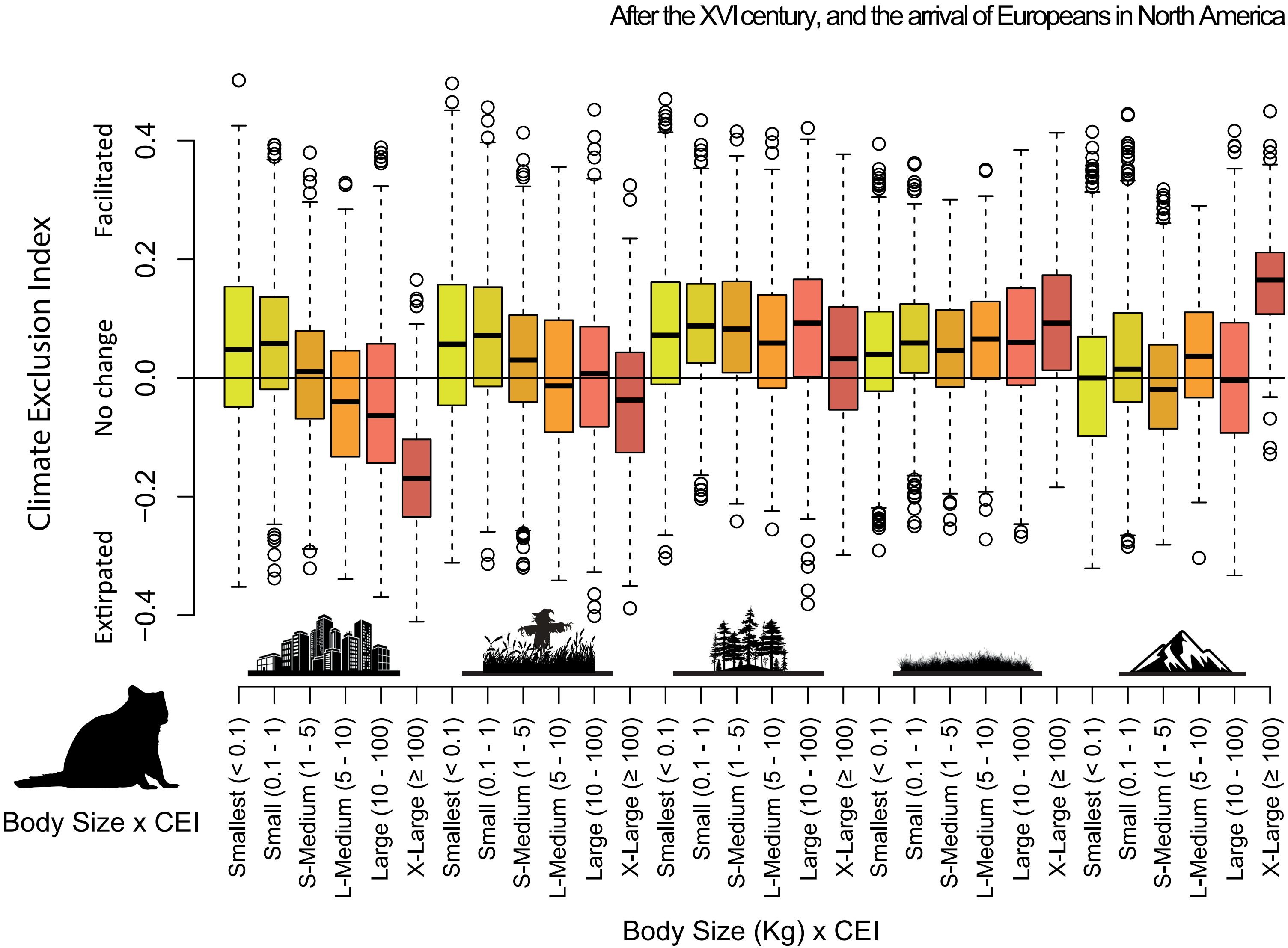

**Supplementary Figure 15**. Distribution of mammal Climate Exclusion Index (CEI) in function of body-size category (Schloss et al. 2012). A positive CEI shows that mammal climatic niches is overlapping more with landscape's climate after 1500 AD. Negative CEI shows extirpation. From left to right: Urban, Cropland, Forest, Grassland, Mountain

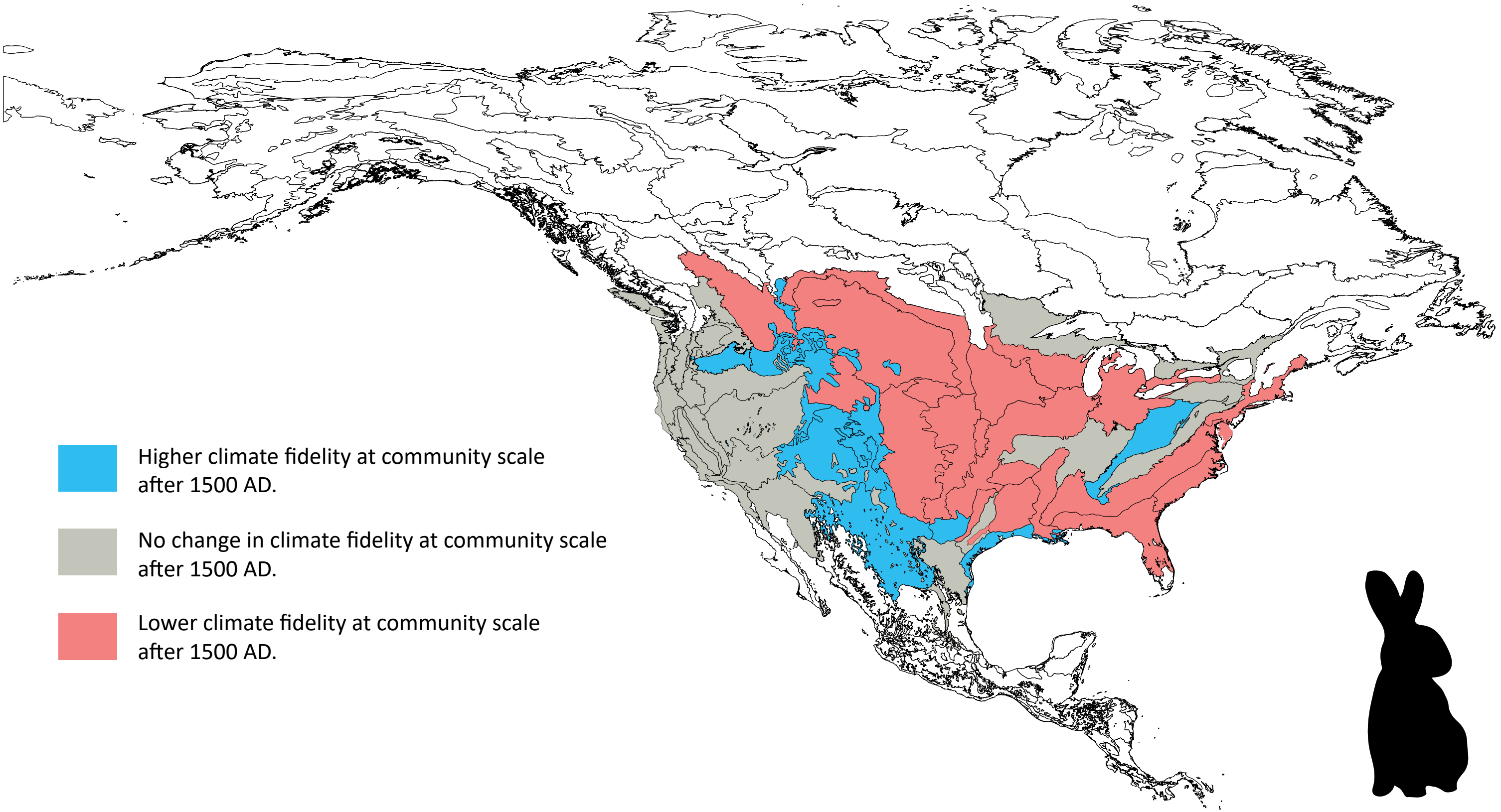

**Supplementary Figure 16.** Change in mammal climate fidelity (CF) at community-scale after the XVIth century. Blue regions have higher CF after 1500 AD, grey regions experienced no change and red regions have lower CF after 1500 AD. Large mammals associated with high CF have been extirpated from central plaints and east coast after the arrival of Europeans.

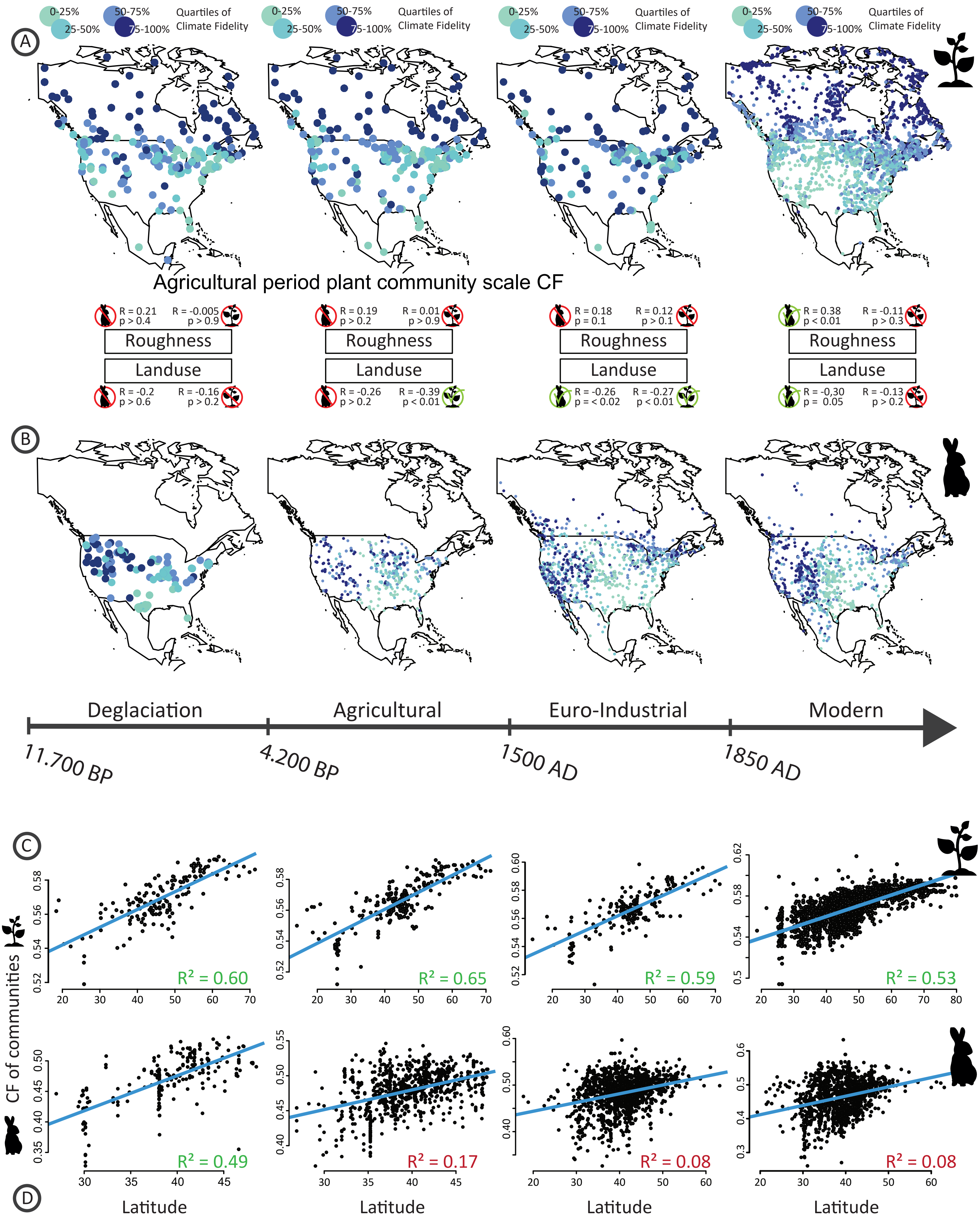

**Supplementary Figure 17.** Temporal variation of the spatial distribution of climate fidelity (CF) at community scale for plants and mammals. Communities are fossil or modern sites with more than 5 taxa. Left is deglaciation, right is modern. A) Spatial distribution of community-scale CF for plants; B) Spatial distribution of community-scale CF for mammals. $R^2$ and p-values of spatial correlation with roughness and landuse patterns are reported and higlighted by red (non-significant relationship) or green signs (significant relationship). C) Latitudinal distribution of community-scale CF for plants; D) Latitudinal distribution of community-scale CF for mammals. Green $R^2$ are significant (p-value < 0.05), red $R^2$ are non-significant (p-value > 0.05).

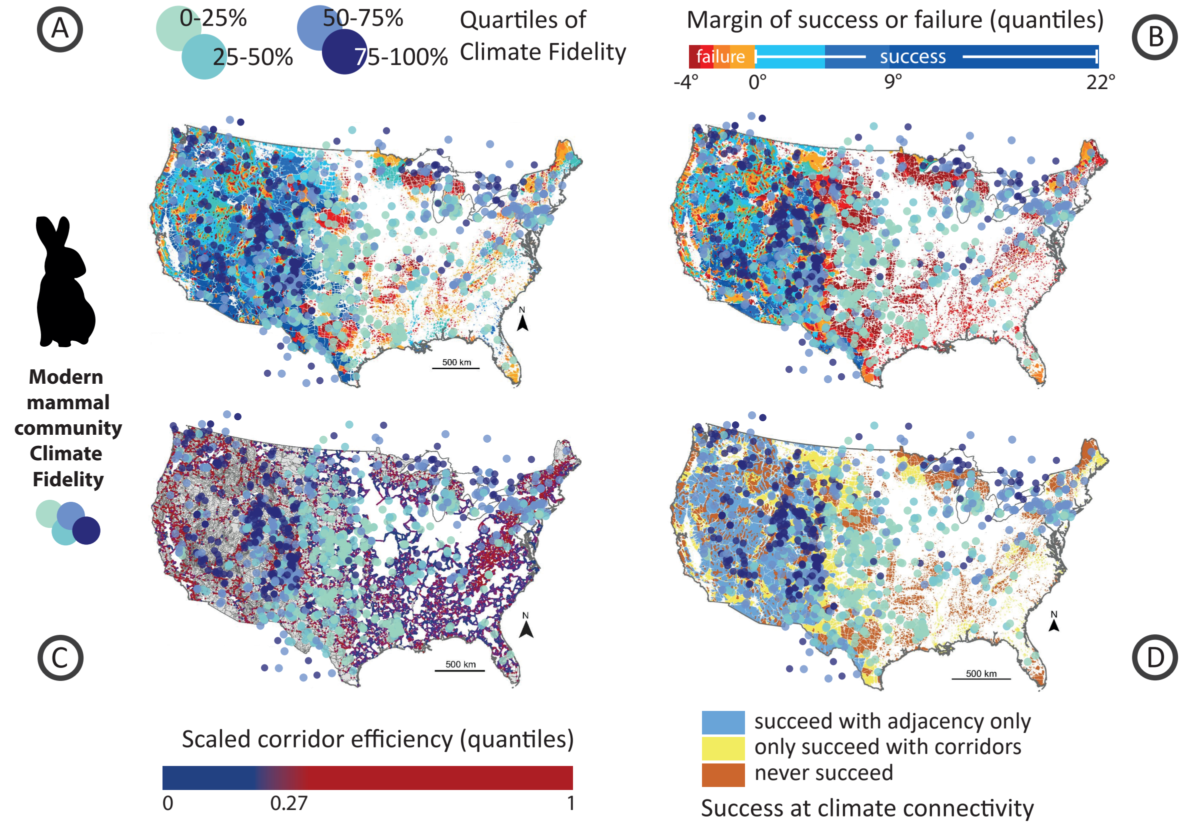

**Figure S18**. Comparison of the spatial distribution of climate fidelity (CF) at modern mammal communities scale with McGuire et al. 2016 connectivity maps. CF at community scale is the average of CF computed on the two oldest time-bins for every mammal present in modern communities, only communities with > 5 taxa are retained. Scale for A and B is above B map.
A) Margin of sucess or failure at achieving climate connectivity with corridors, B) and without corridors, C) Corridor efficiency scaled to the maximum value, D) Improvement due to corridors.

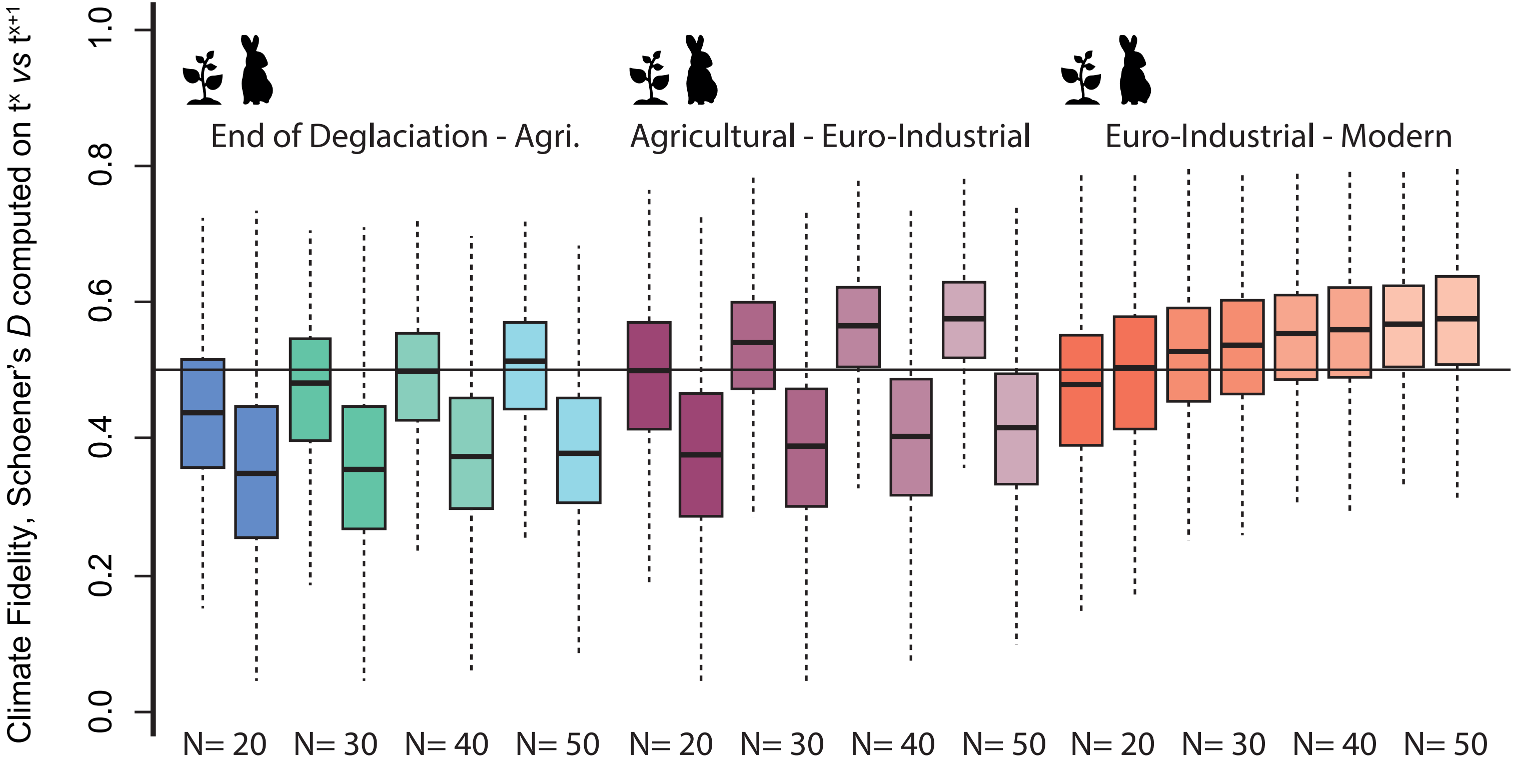

**Supplementary Figure 19.** Sensitivity of climate fidelity (CF) to sample size, from N = 50 to N = 20. Blue-green boxplots are computed between the End of Deglaciation and the Agricultural periods; Purple-pink boxplots are computed between the Agricultural and the EuroIndustrial periods; Orange boxplots are computed between the EuroIndustrial and Modern periods. For any sample size (e.g. 20), left boxplots depict plant CF, right boxplots depict animal CF

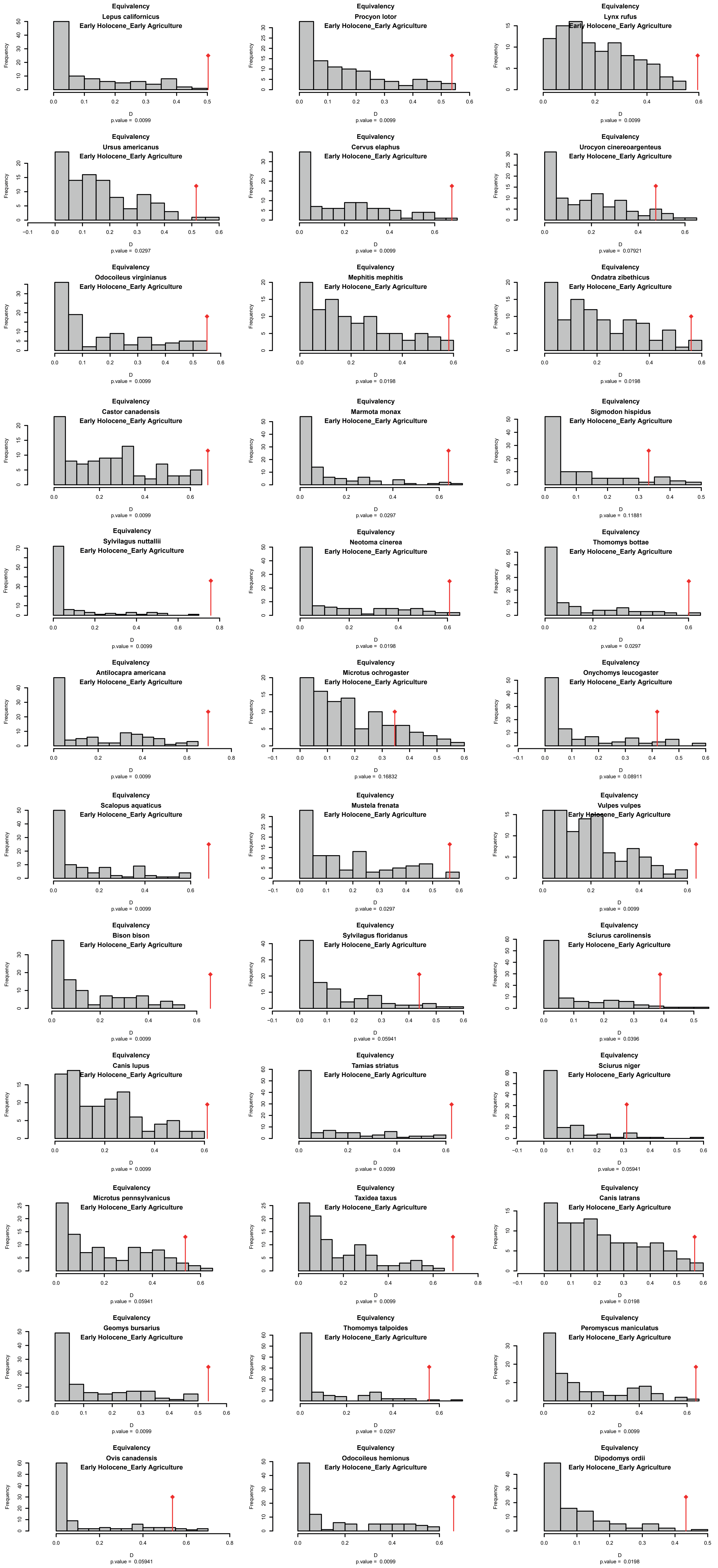

**Supplementary Figure S20.** Tests for niche conservatism (Equivalency, see Warren et al. 2008) of climatic niches associated with Schoener's *D* index computation (ecospat R package: Di Cola et al. 2017). Temporal comparison of mammalian climatic niches between the End of Deglaciation (11.700 BP - 4.200 BP, named Early Holocene here) and the Agricultural period (4.200 BP - 1500 AD, named Early Agriculture here).

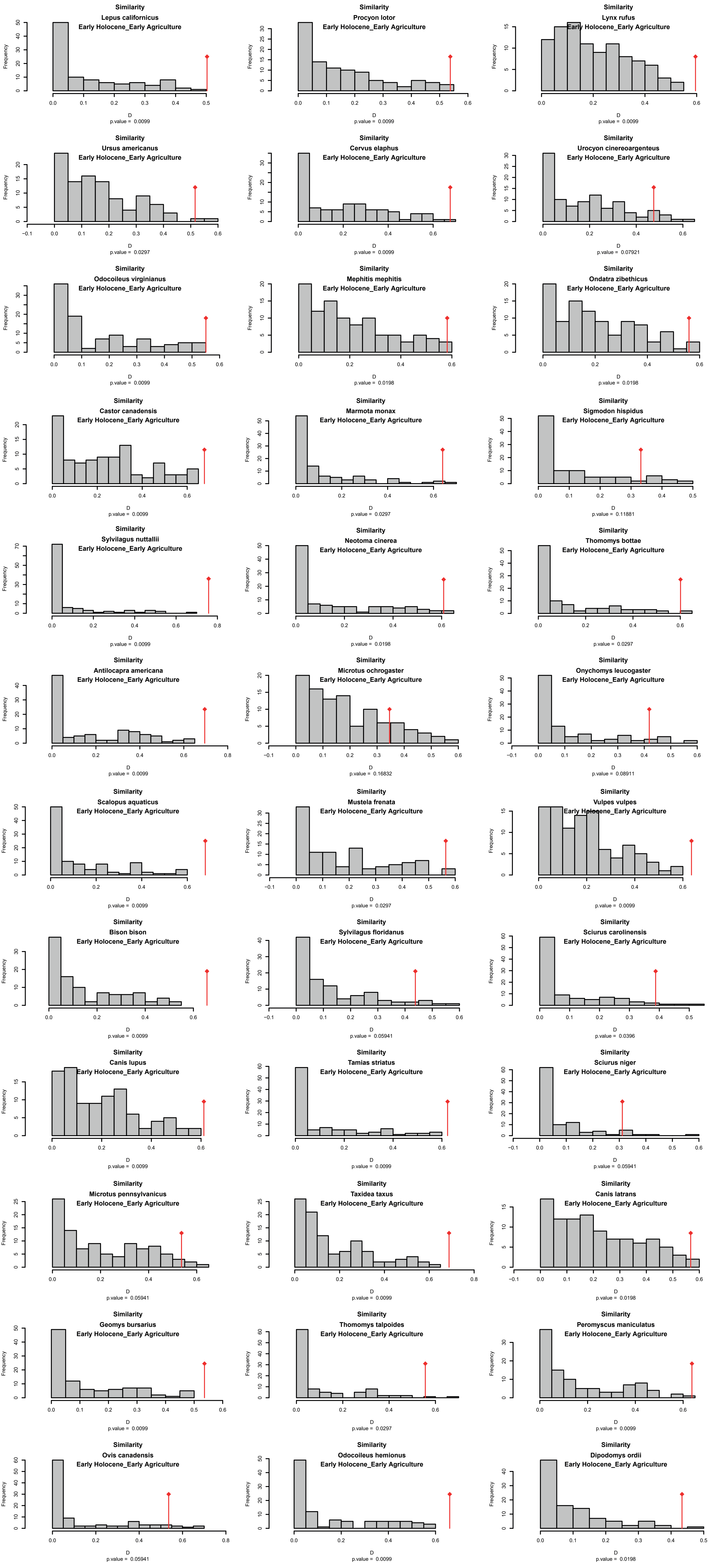

**Supplementary Figure S21**. Tests for niche conservatism (Similarity, see Warren et al. 2008) of climatic niches associated with Schoener's *D* index computation (ecospat R package: Di Cola et al. 2017). Temporal comparison of mammalian climatic niches between the End of Deglaciation (11.700 BP - 4.200 BP, named Early Holocene here) and the Agricultural period (4.200 BP - 1500 AD, named Early Agriculture here).

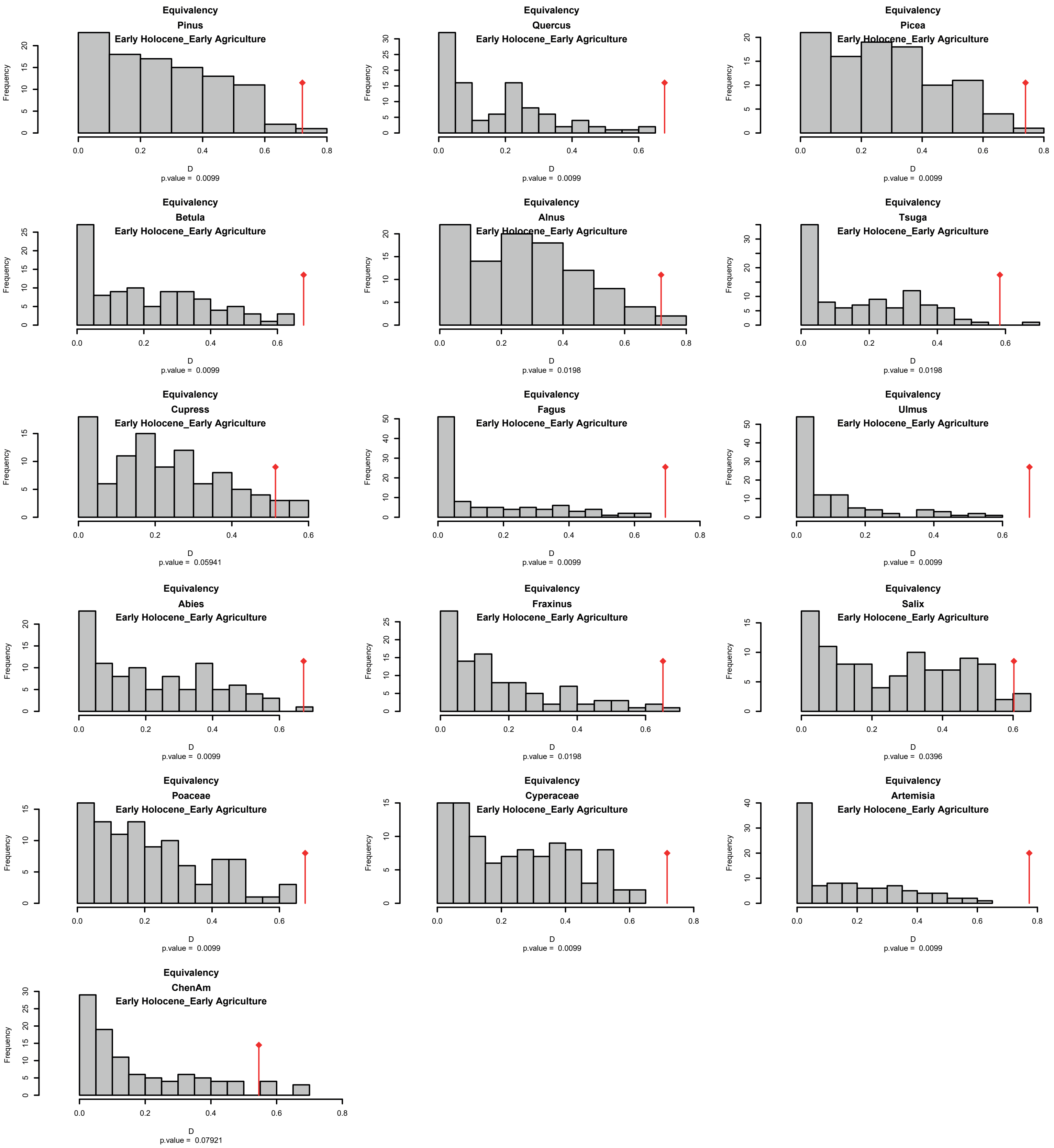

**Supplementary Figure S22**. Tests for niche conservatism (Equivalency, see Warren et al. 2008) of climatic niches associated with Schoener's *D* index computation (ecospat R package: Di Cola et al. 2017). Temporal comparison of plant climatic niches between the end of Deglaciation (11.700 BP - 4.200 BP, named Early Holocene here) and the Agricultural period (4.200 BP - 1500 AD, named Early Agriculture here).

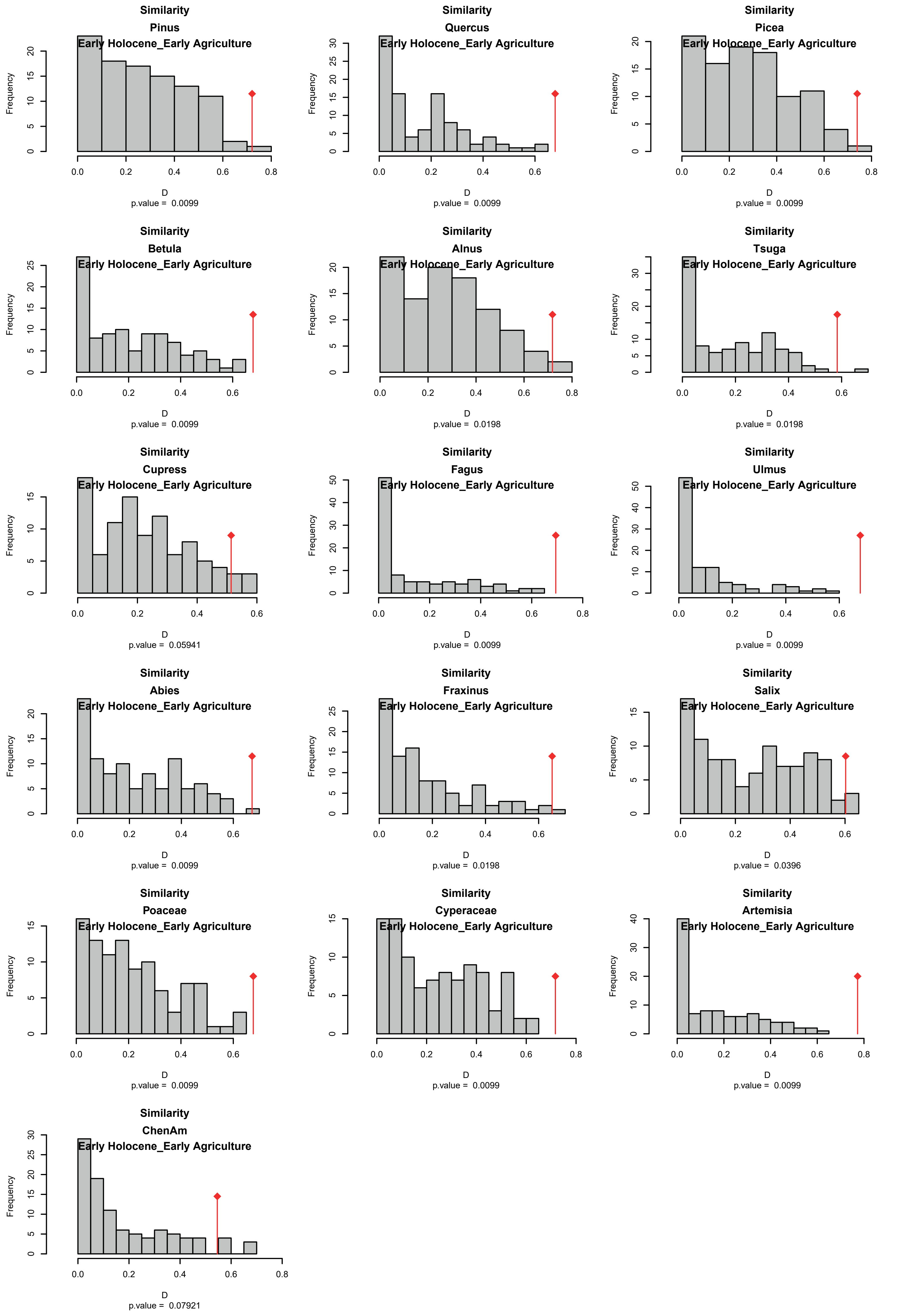

**Supplementary Figure S23**. Tests for niche conservatism (Similarity, see Warren et al. 2008) of climatic niches associated with Schoener's *D* index computation (ecospat R package: Di Cola et al. 2017). Temporal comparison of plant climatic niches between the End of Deglaciation (11.700 BP - 4.200 BP, named Early Holocene here) and the Agricultural period (4.200 BP - 1500 AD, named Early Agriculture here).

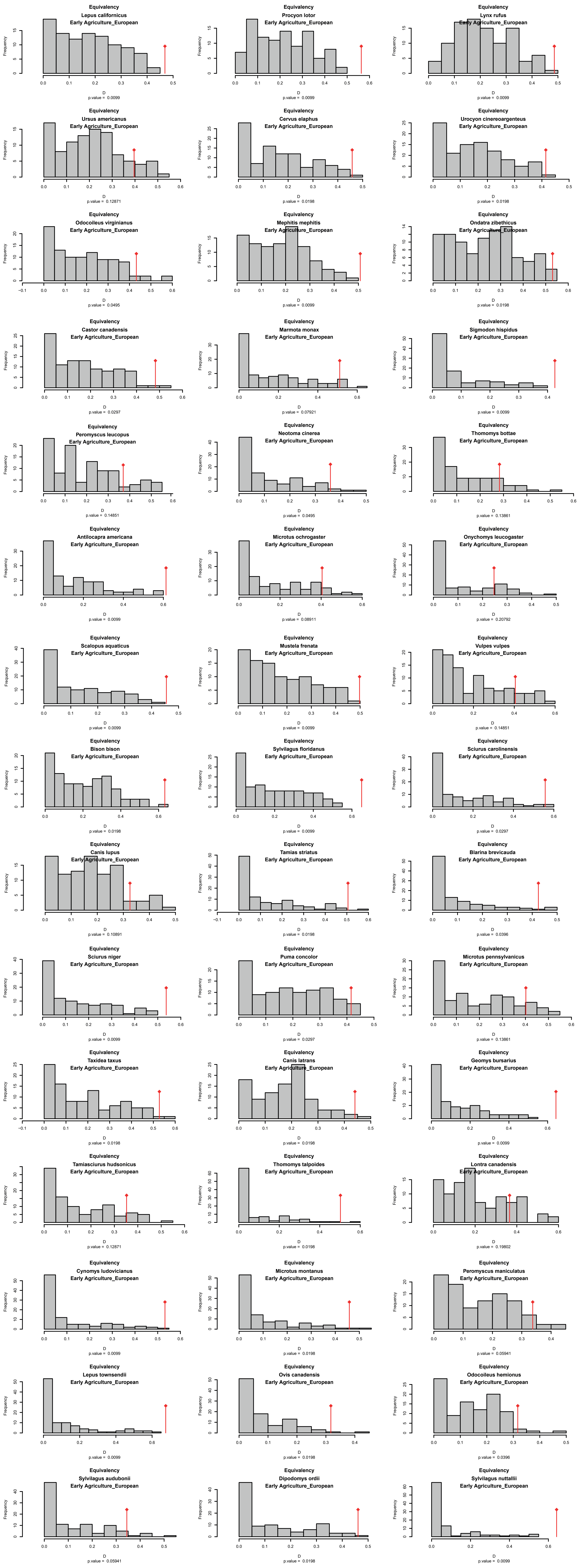

**Supplementary Figure S24**. Tests for niche conservatism (Equivalency, see Warren et al. 2008) of climatic niches associated with Schoener's *D* index computation (ecospat R package: Di Cola et al. 2017). Temporal comparison of mammalian climatic niches between the Agricultural period (4.200 BP - 1500 AD, named Early Agriculture here) and the EuroIndustrial period (1500 AD - 1950 AD, named European here).

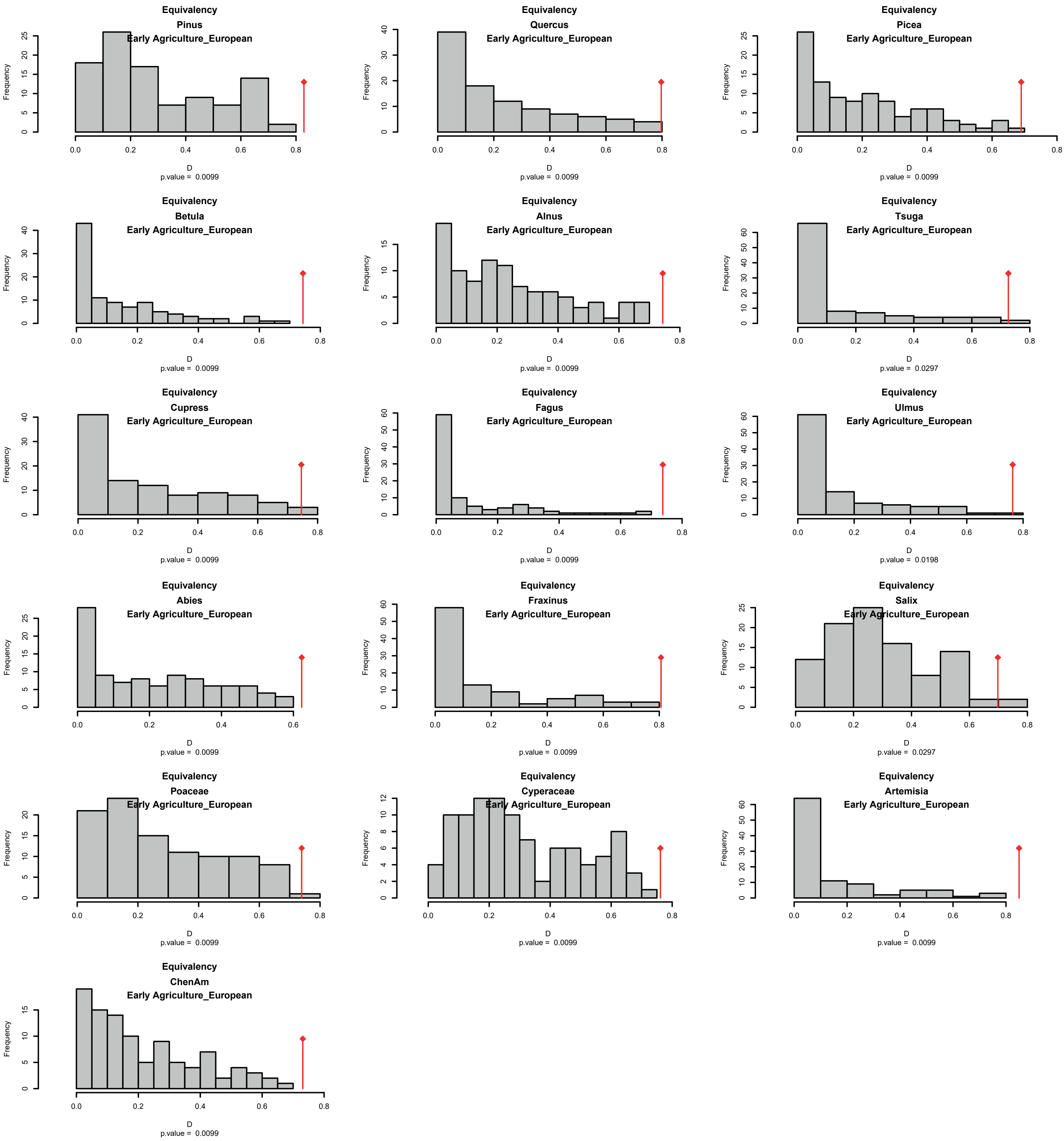

**Supplementary Figure S25**. Tests for niche conservatism (Equivalency, see Warren et al. 2008) of climatic niches associated with Schoener's *D* index computation (ecospat R package: Di Cola et al. 2017). Temporal comparison of plant climatic niches between the Agricultural period (4.200 BP - 1500 AD, named Early Agriculture here) and the EuroIndustrial period (1500 AD - 1950 AD, named European here).

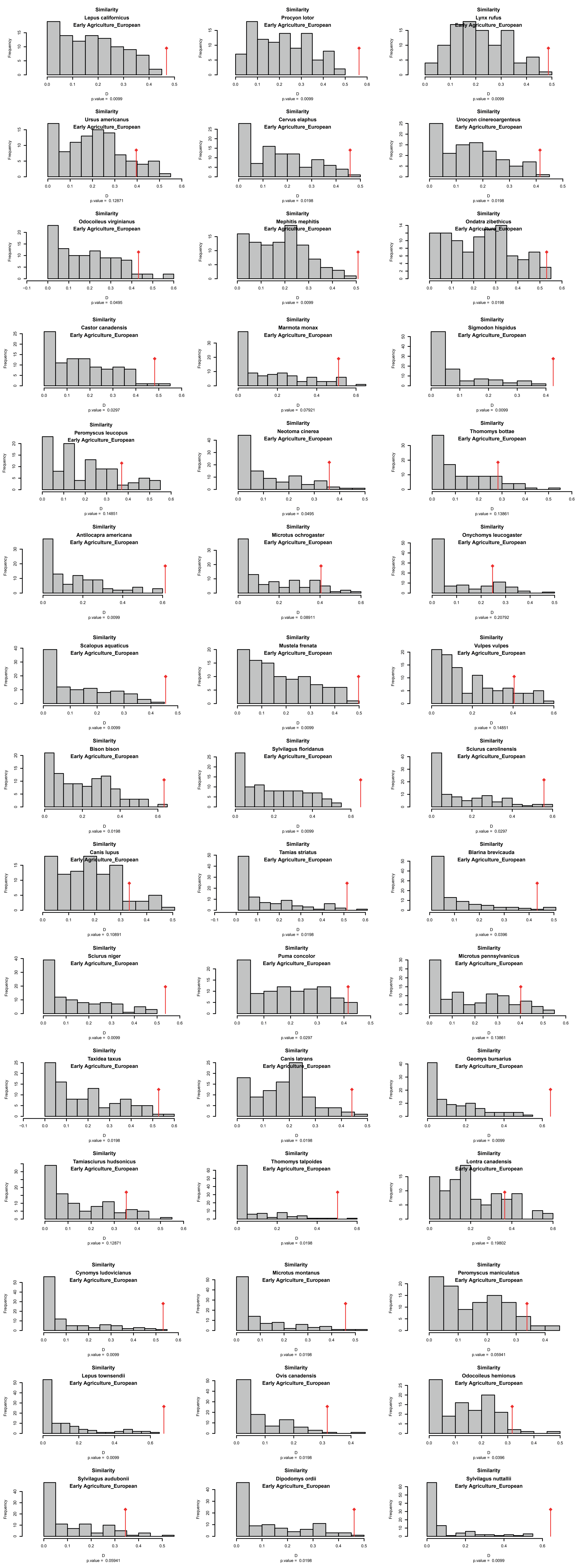

**Supplementary Figure S26.** Tests for niche conservatism (Similarity, see Warren et al. 2008) of climatic niches associated with Schoener's *D* index computation (ecospat R package: Di Cola et al. 2017). Temporal comparison of mammalian climatic niches between the Agricultural period (4.200 BP - 1500 AD, named Early Agriculture here) and the EuroIndustrial period (1500 AD - 1950 AD, named European here).

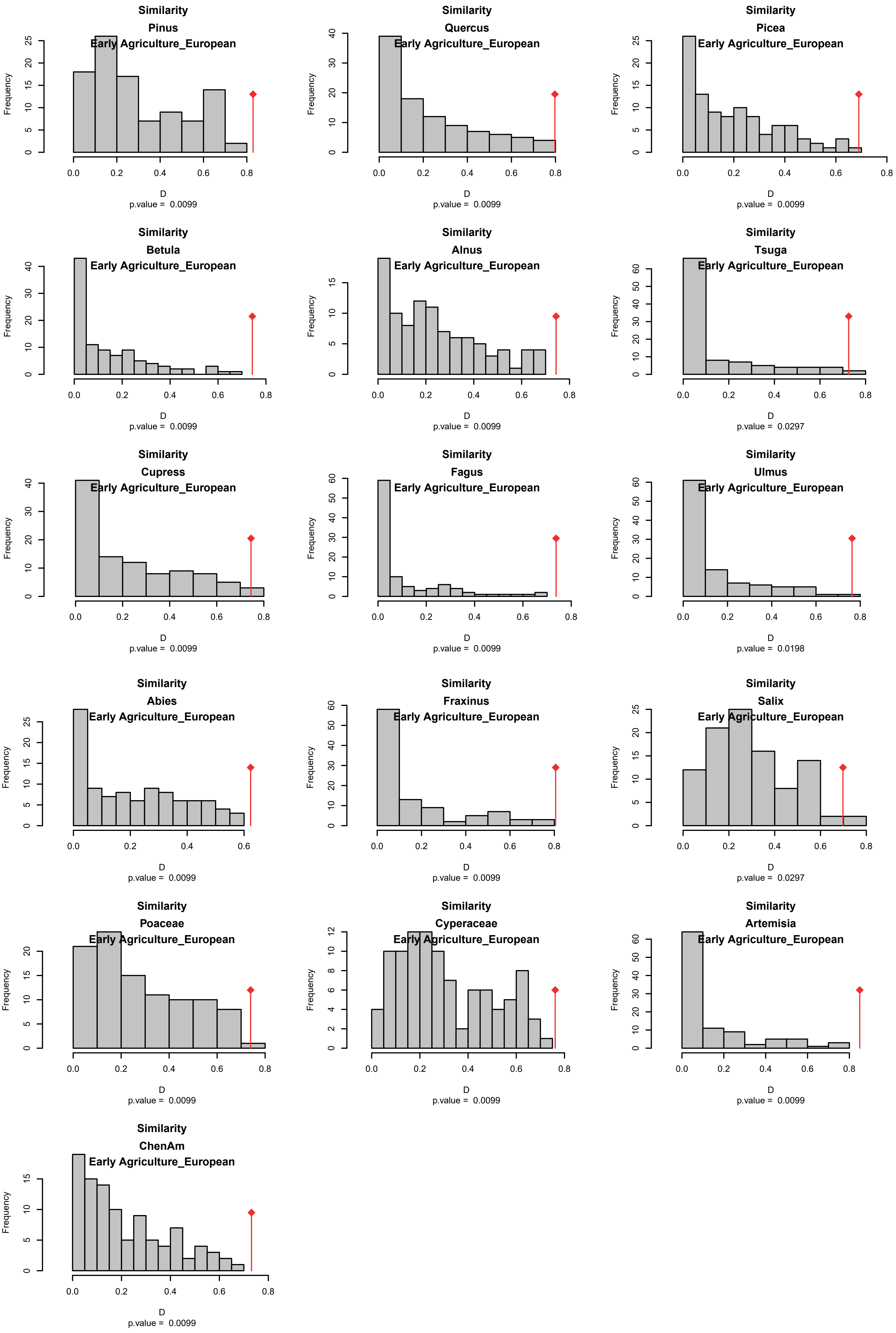

**Supplementary Figure S27**. Tests for niche conservatism (Similarity, see Warren et al. 2008) of climatic niches associated with Schoener's *D* index computation (ecospat R package: Di Cola et al. 2017). Temporal comparison of plant climatic niches between the Agricultural period (4.200 BP - 1500 AD, named Early Agriculture here) and the EuroIndustrial period (1500 AD - 1950 AD, named European here).

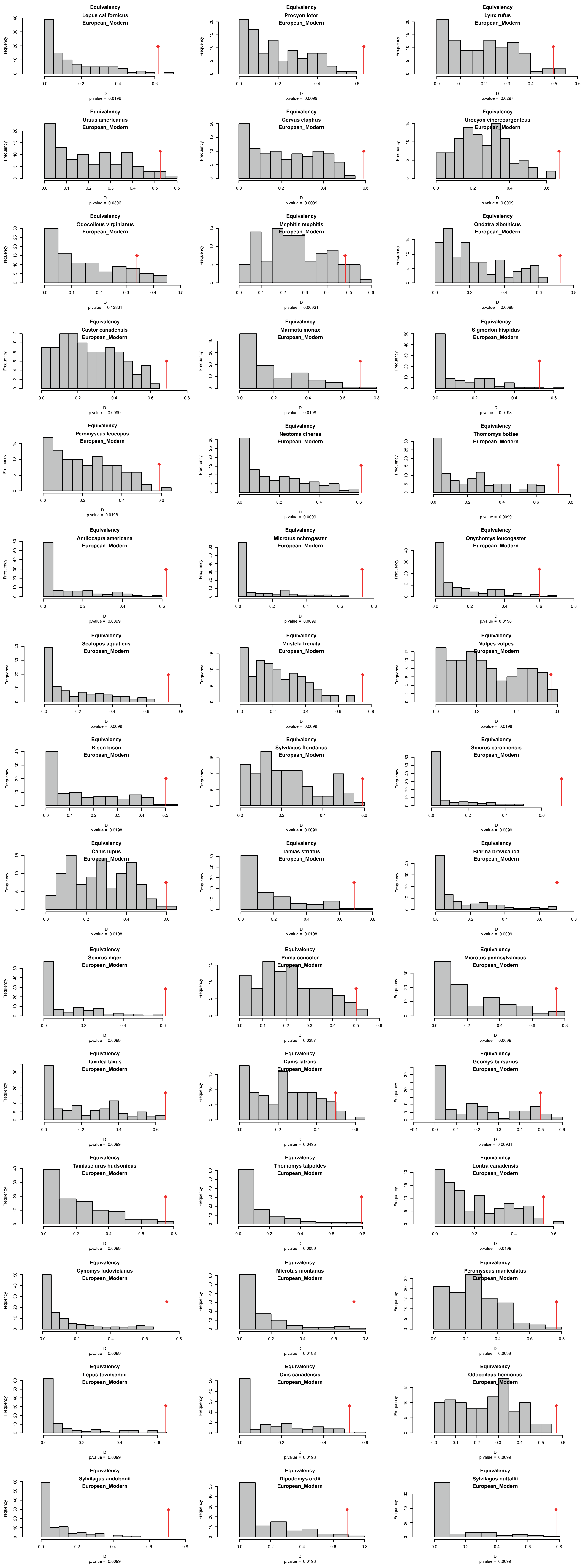

**Supplementary Figure S28**. Tests for niche conservatism (Equivalency, see Warren et al. 2008) of climatic niches associated with Schoener's *D* index computation (ecospat R package: Di Cola et al. 2017). Temporal comparison of mammalian climatic niches between the EuroIndustrial period (1500 AD - 1950 AD, named European here) and the Modern period (1950 - ).

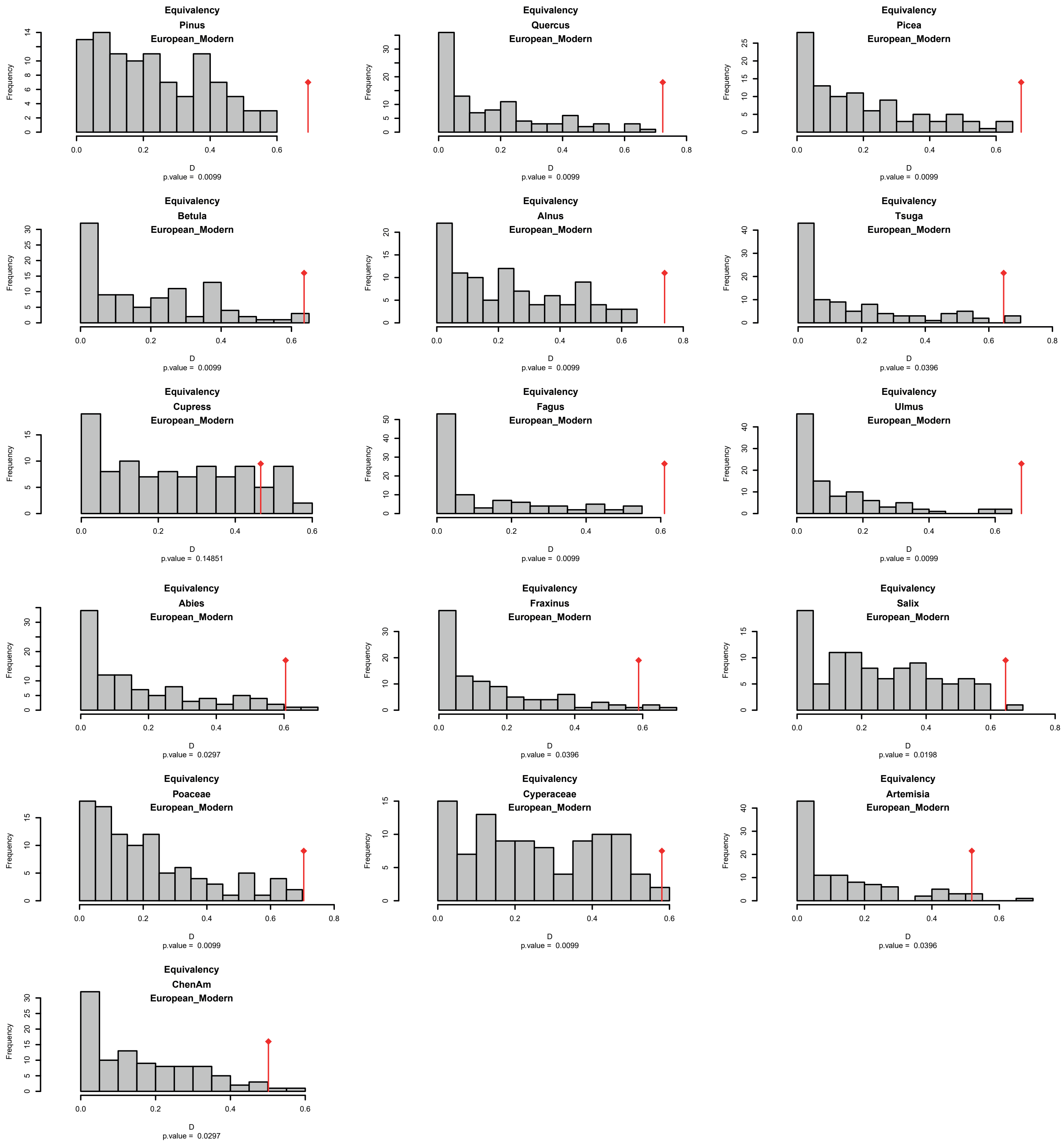

**Supplementary Figure S29**. Tests for niche conservatism (Equivalency, see Warren et al. 2008) of climatic niches associated with Schoener's *D* index computation (ecospat R package: Di Cola et al. 2017). Temporal comparison of plant climatic niches between the EuroIndustrial period (1500 AD - 1950 AD, named European here) and the Modern period (1950 - ).

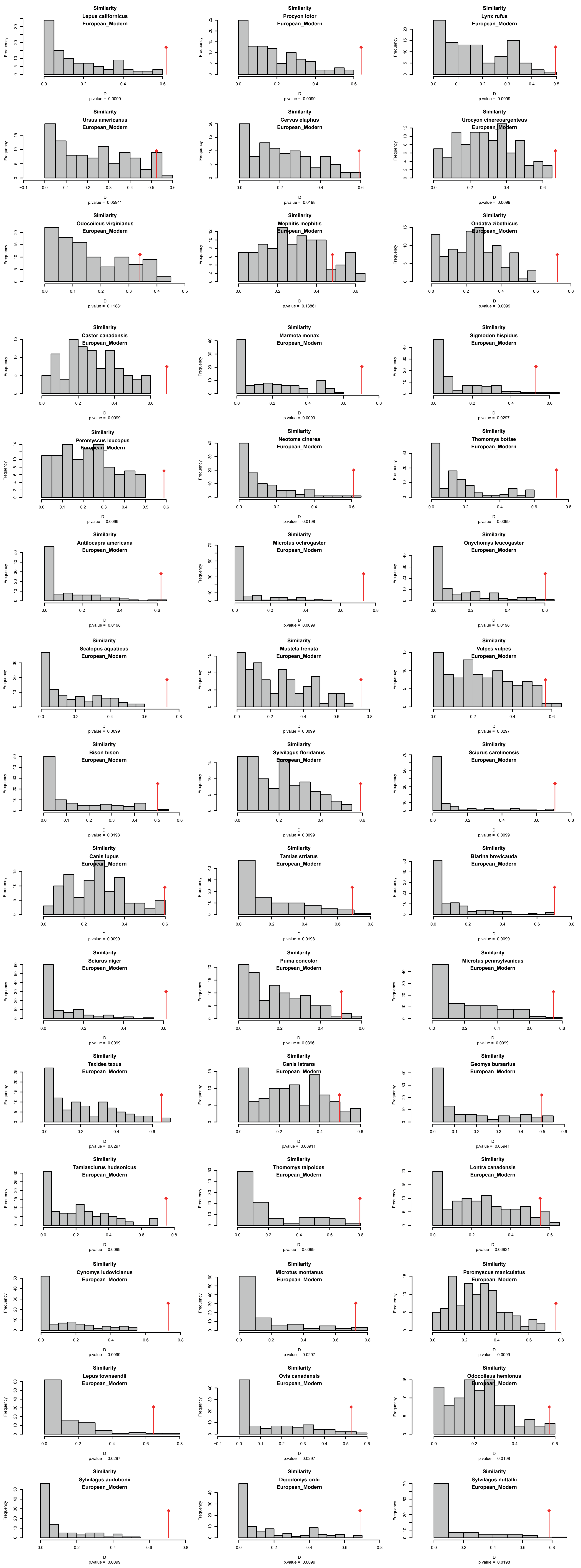

**Supplementary Figure S30**. Tests for niche conservatism (Similarity, see Warren et al. 2008) of climatic niches associated with Schoener's *D* index computation (ecospat R package: Di Cola et al. 2017). Temporal comparison of mammalian climatic niches between the EuroIndustrial period (1500 AD - 1950 AD, named European here) and the Modern period (1950 - ).

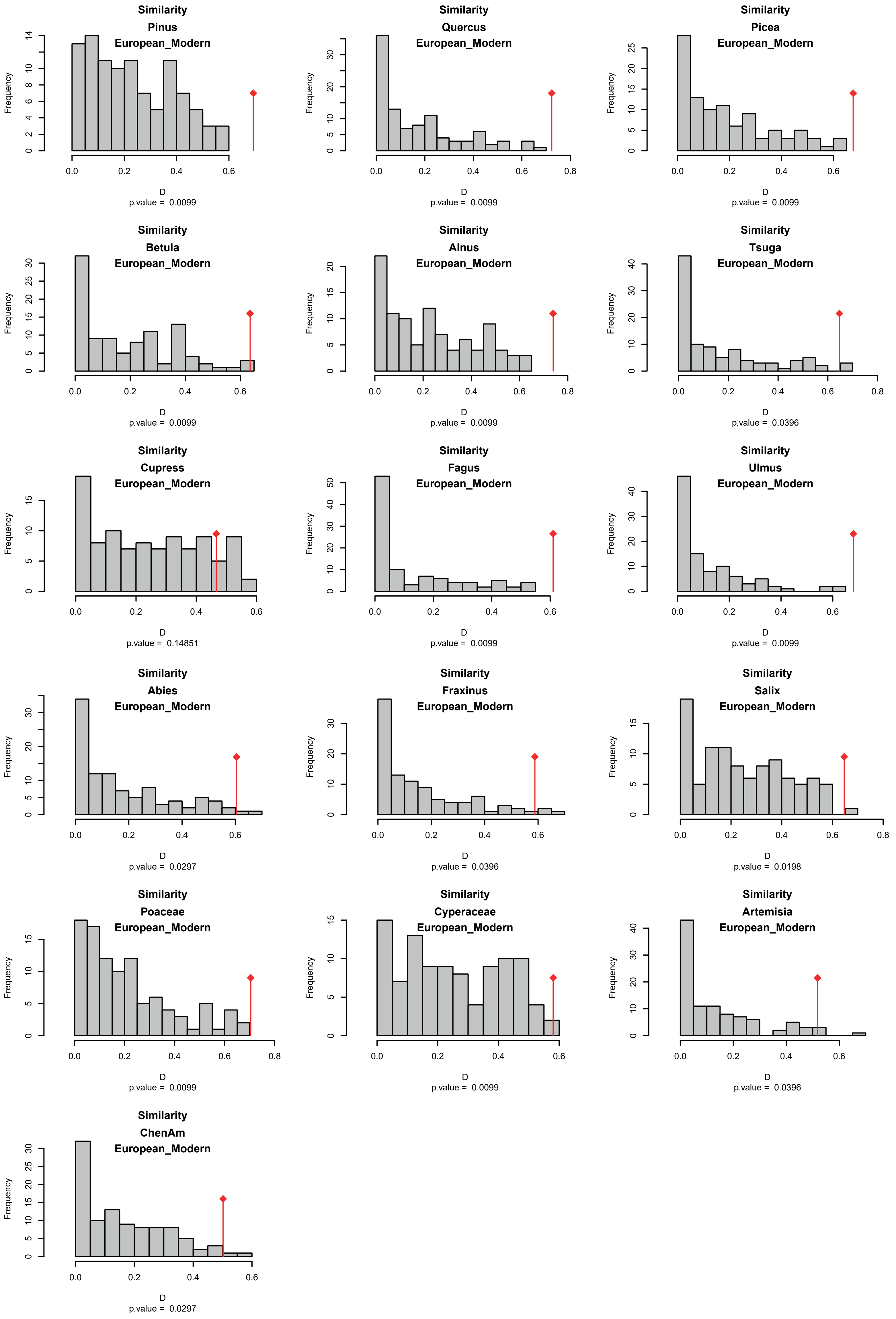

**Supplementary Figure S31**. Tests for niche conservatism (Similarity, see Warren et al. 2008) of climatic niches associated with Schoener's *D* index computation (ecospat R package: Di Cola et al. 2017). Temporal comparison of plant climatic niches between the EuroIndustrial period (1500 AD - 1950 AD, named European here) and the Modern period (1950 - ).

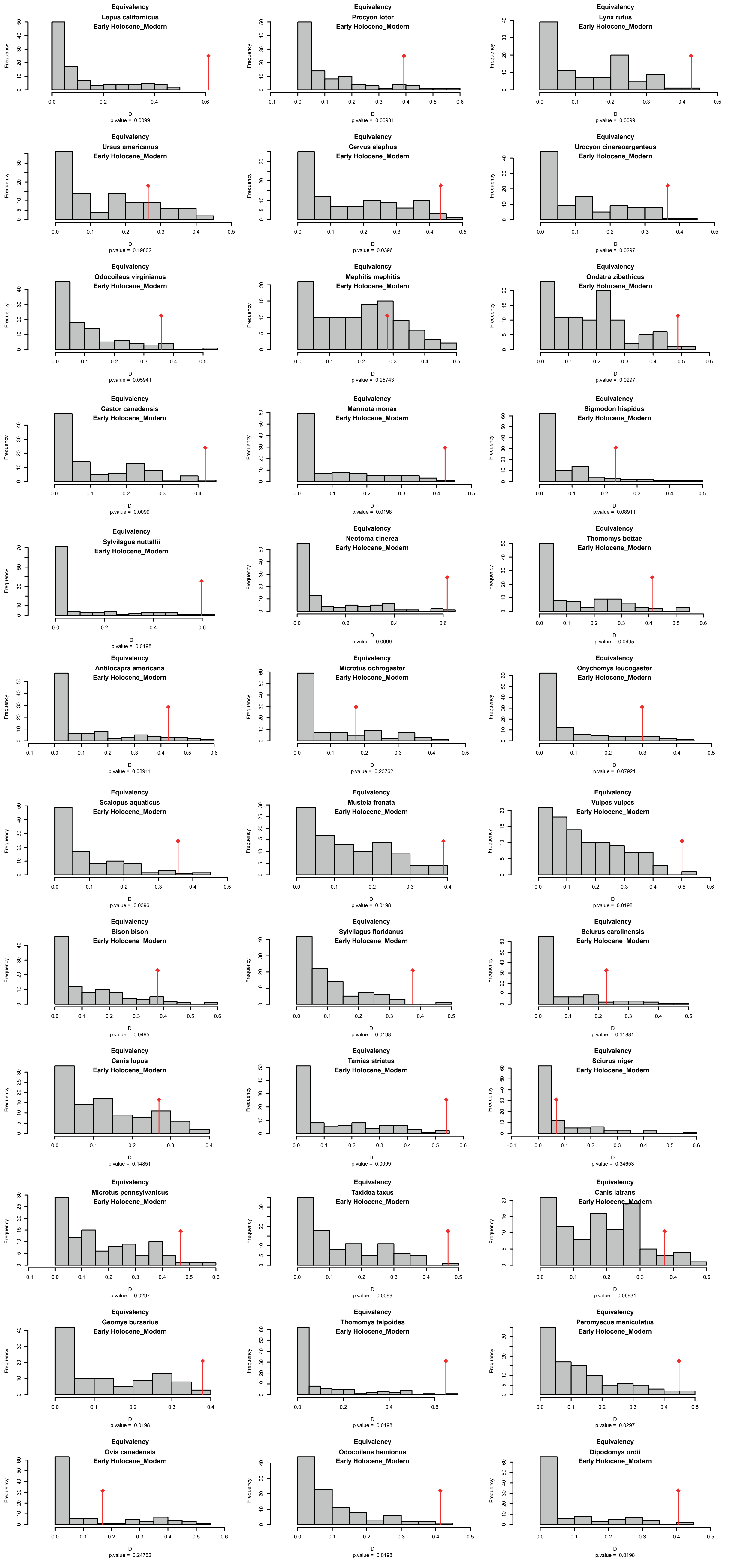

**Supplementary Figure S32**. Tests for niche conservatism (Equivalency, see Warren et al. 2008) of climatic niches associated with Schoener's *D* index computation (ecospat R package: Di Cola et al. 2017). Temporal comparison of mammalian climatic niches between the End of Deglaciation (11.700 BP - 4.200 AD, named Early Holocene here) and the Modern period (1950 AD - ).

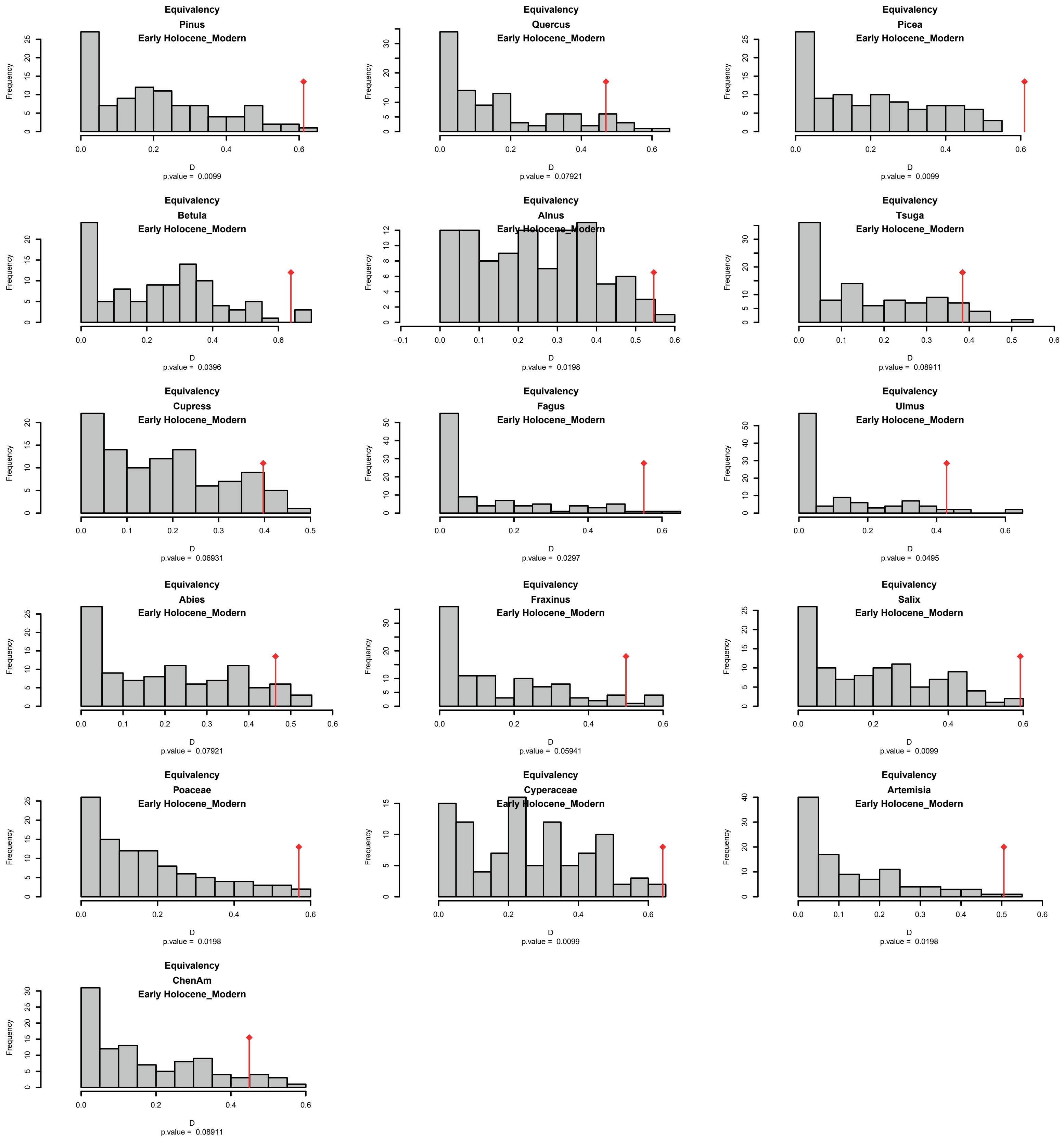

**Supplementary Figure S33**. Tests for niche conservatism (Equivalency, see Warren et al. 2008) of climatic niches associated with Schoener's *D* index computation (ecospat R package: Di Cola et al. 2017). Temporal comparison of plant climatic niches between the End of Deglaciation (11.700 BP - 4.200 AD, named Early Holocene here) and the Modern period (1950 AD - ).

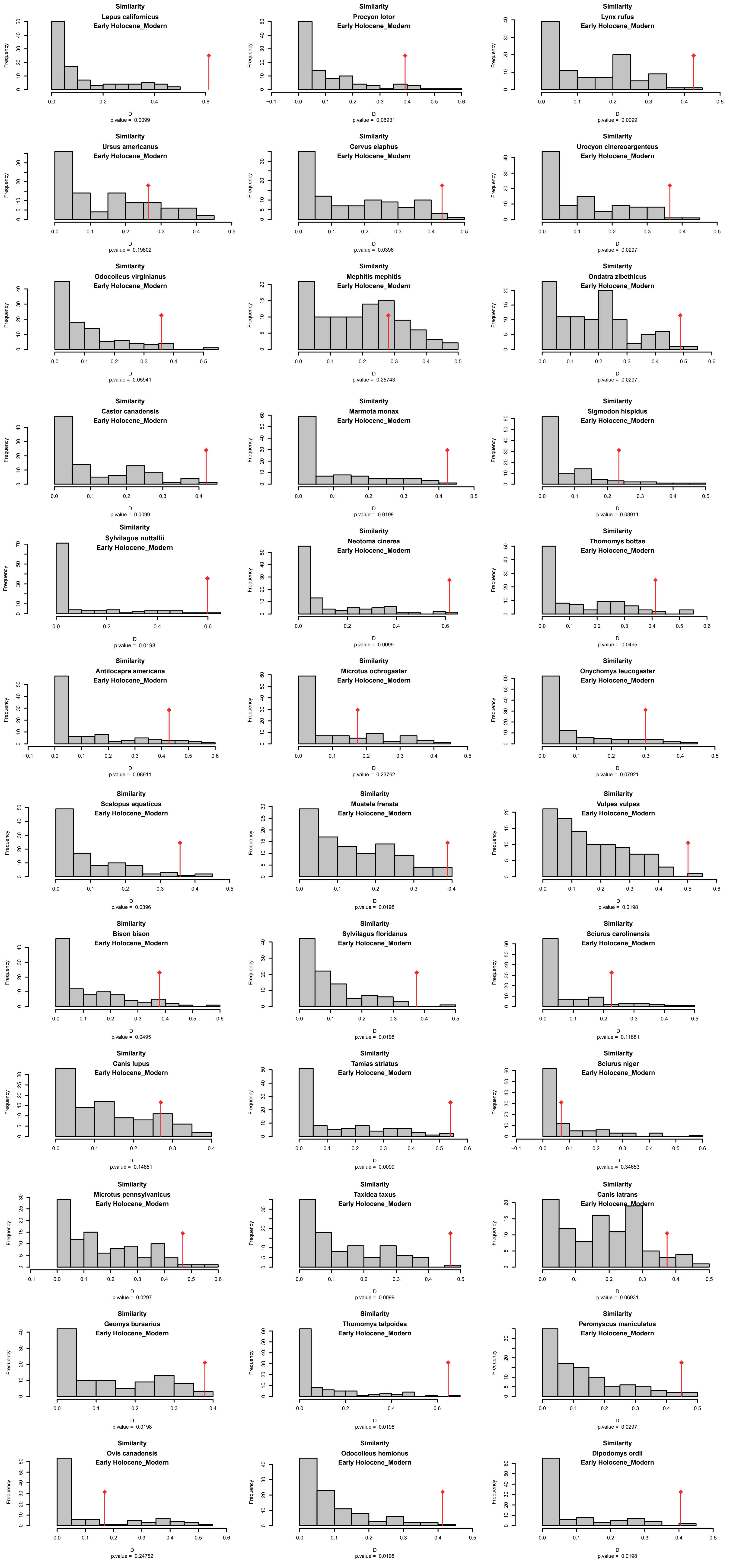

**Supplementary Figure S34**. Tests for niche conservatism (Similarity, see Warren et al. 2008) of climatic niches associated with Schoener's *D* index computation (ecospat R package: Di Cola et al. 2017). Temporal comparison of mammalian climatic niches between the End of Deglaciation (11.700 BP - 4.200 AD, named Early Holocene here) and the Modern period (1950 AD - ).

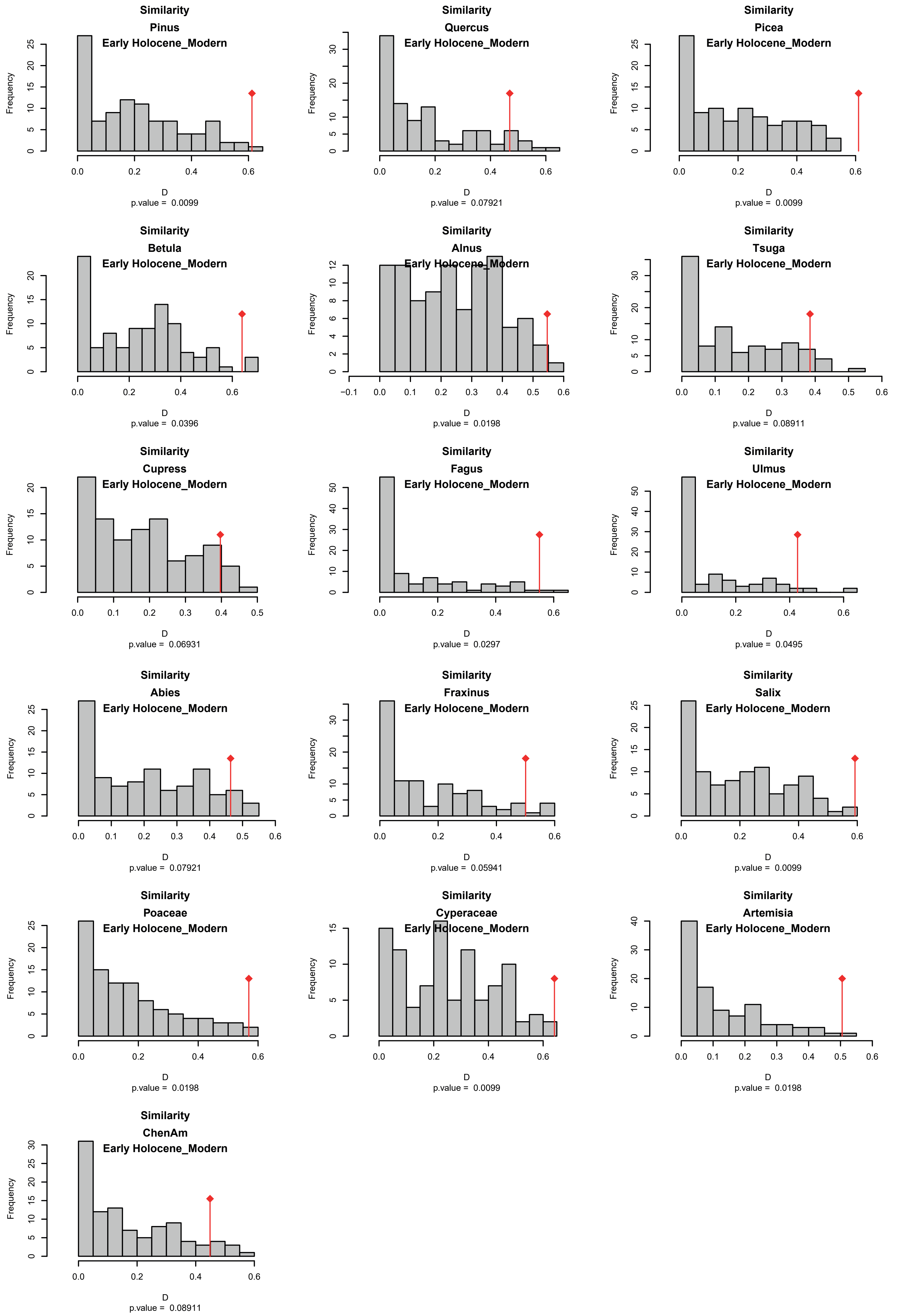

**Supplementary Figure S35.** Tests for niche conservatism (Similarity, see Warren et al. 2008) of climatic niches associated with Schoener's *D* index computation (ecospat R package: Di Cola et al. 2017). Temporal comparison of plant climatic niches between the End of Deglaciation (11.700 BP - 4.200 AD, named Early Holocene here) and the Modern period (1950 AD - ).

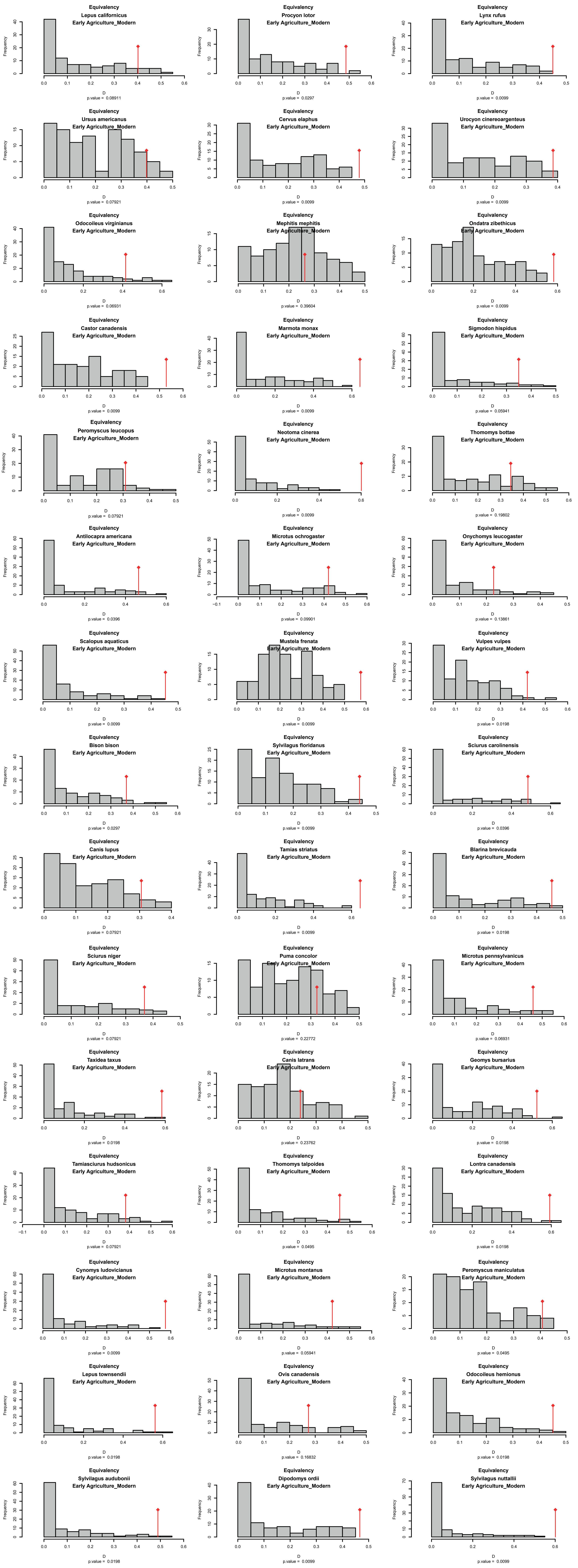

**Supplementary Figure S36.** Tests for niche conservatism (Equivalency, see Warren et al. 2008) of climatic niches associated with Schoener's *D* index computation (ecospat R package: Di Cola et al. 2017). Temporal comparison of mammalian climatic niches between the Agricultural period (4.200 AD - 1.500 AD, named Early Agriculture here) and the Modern period (1950 AD - ).

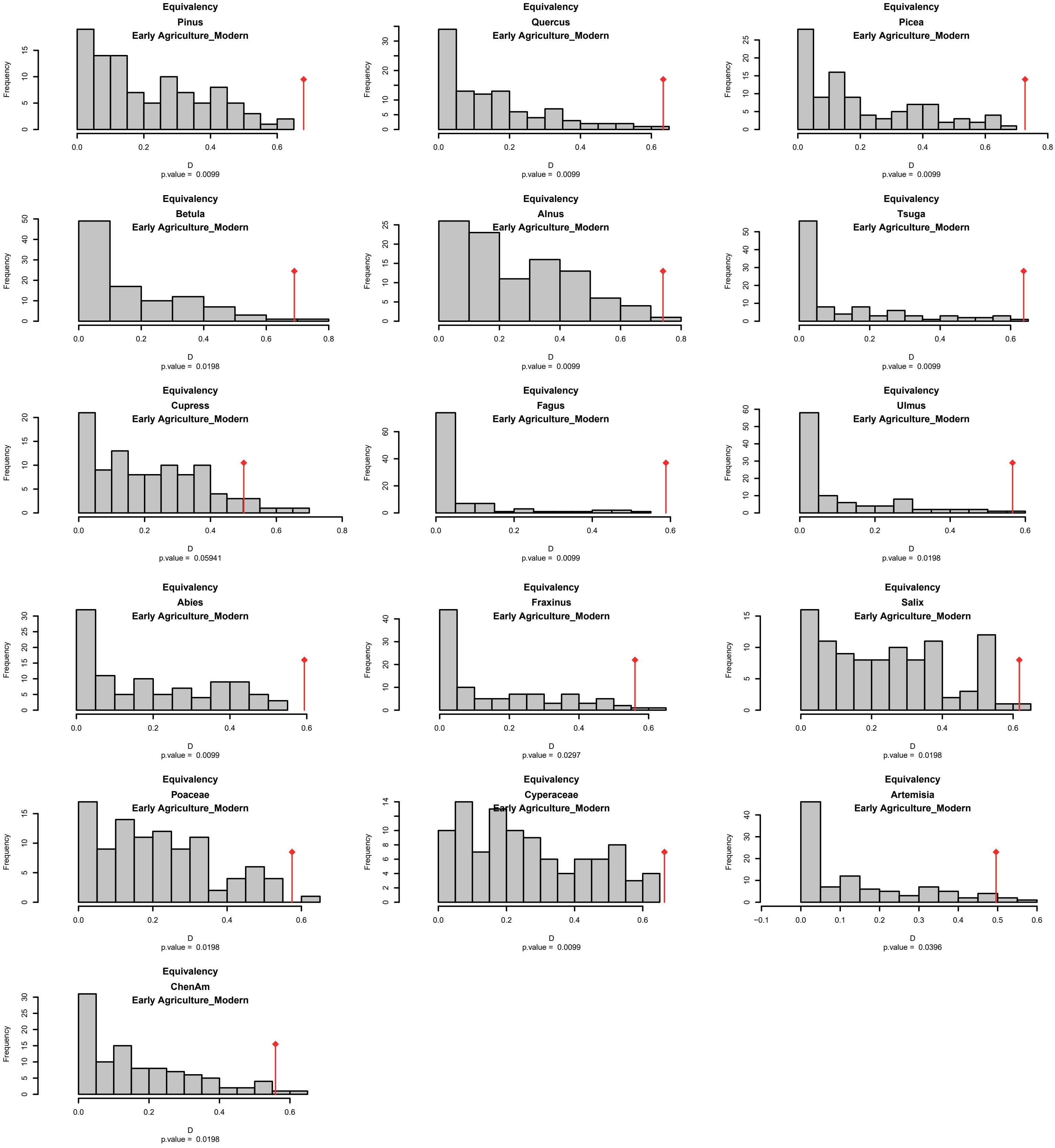

**Supplementary Figure S37**. Tests for niche conservatism (Equivalency, see Warren et al. 2008) of climatic niches associated with Schoener's *D* index computation (ecospat R package: Di Cola et al. 2017). Temporal comparison of plant climatic niches between the Agricultural period (4.200 AD - 1.500 AD, named Early Agriculture here) and the Modern period (1950 AD - ).

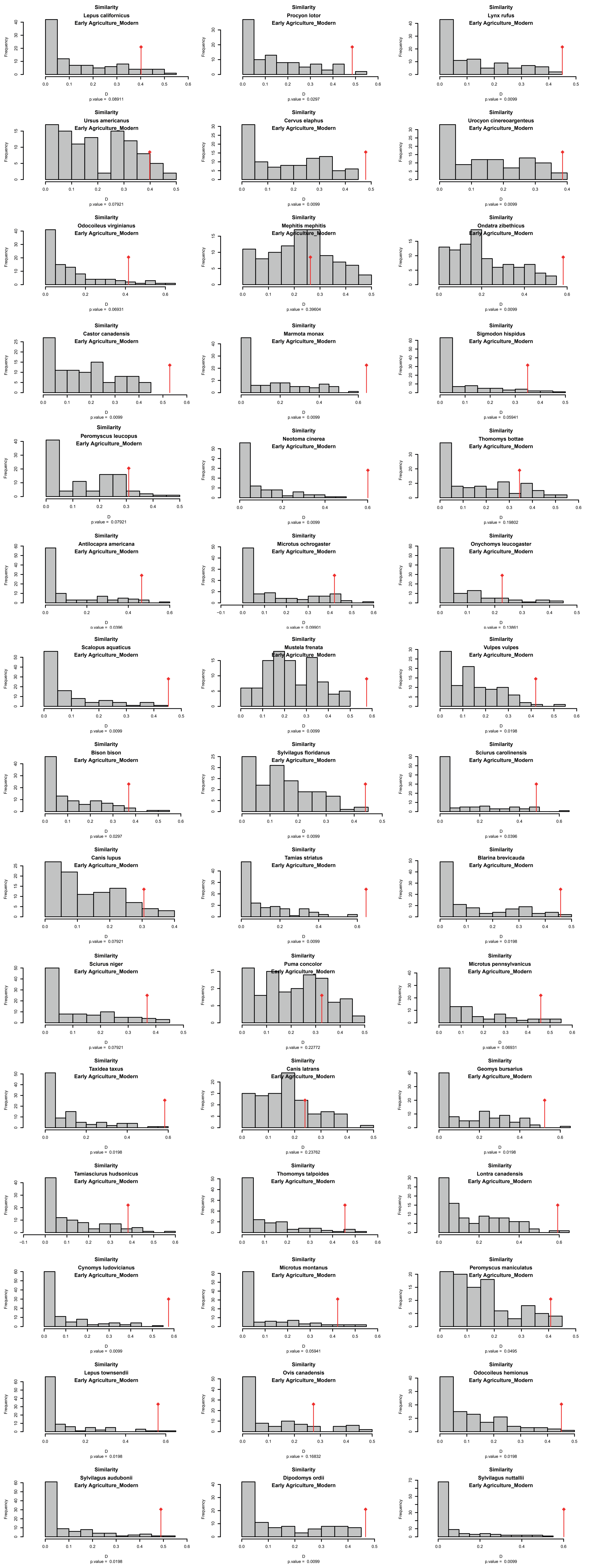

**Supplementary Figure S38.** Tests for niche conservatism (Similarity, see Warren et al. 2008) of climatic niches associated with Schoener's *D* index computation (ecospat R package: Di Cola et al. 2017). Temporal comparison of mammalian climatic niches between the Agricultural period (4.200 AD - 1.500 AD, named Early Agriculture here) and the Modern period (1950 AD - ).

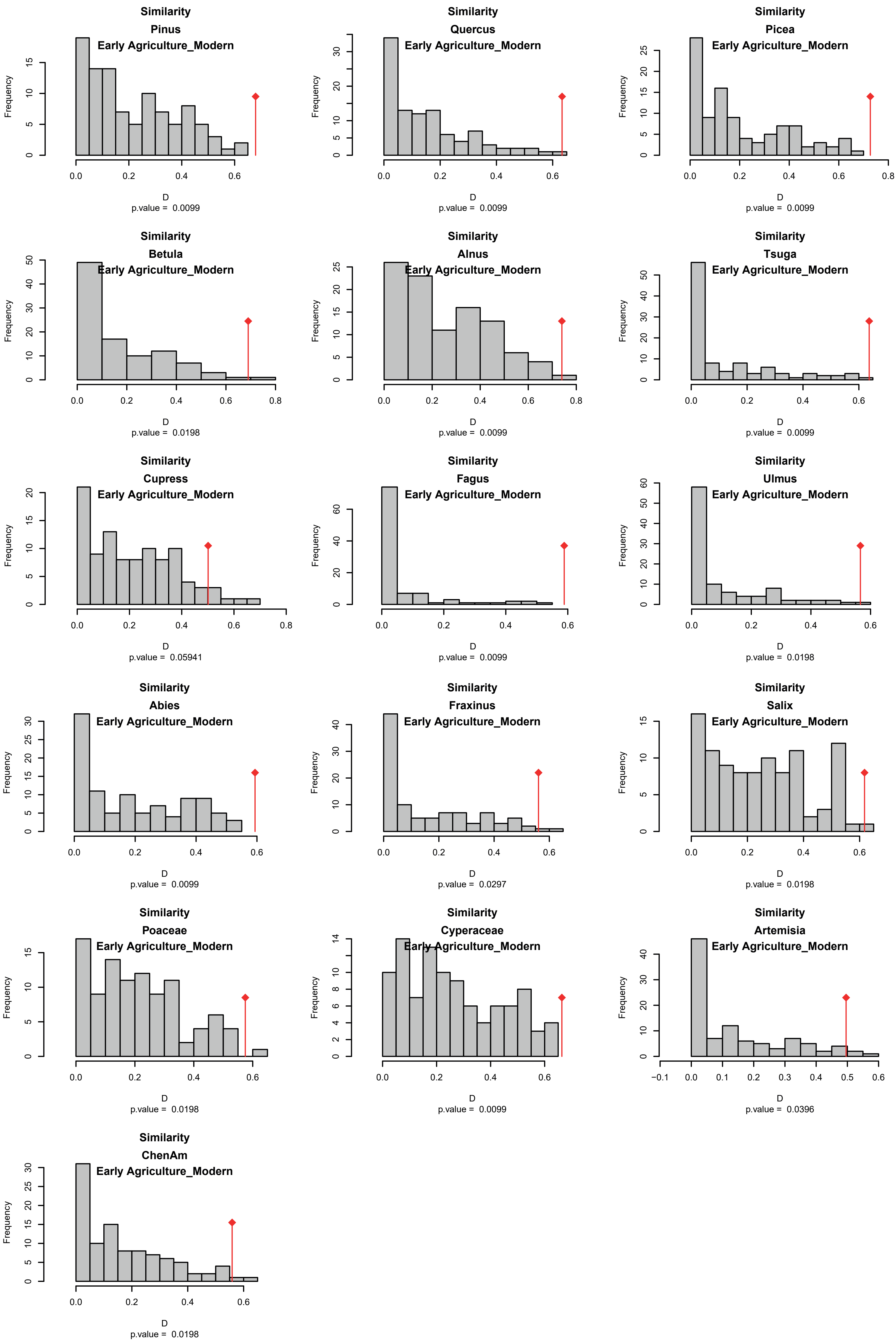

**Supplementary Figure S39**. Tests for niche conservatism (Similarity, see Warren et al. 2008) of climatic niches associated with Schoener's *D* index computation (ecospat R package: Di Cola et al. 2017). Temporal comparison of plant climatic niches between the Agricultural period (4.200 AD - 1.500 AD, named Early Agriculture here) and the Modern period (1950 AD - ).

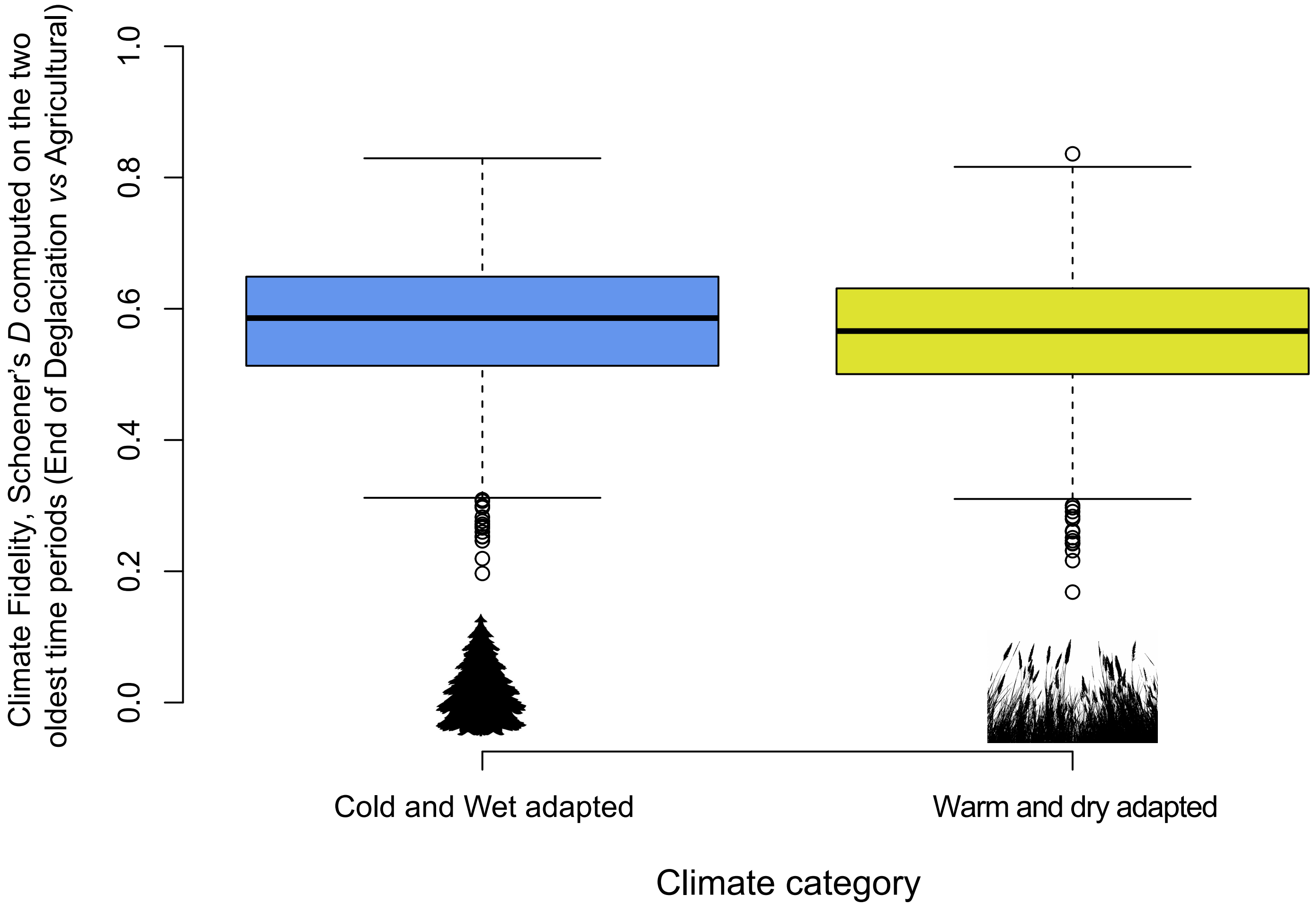

**Supplementary Figure 40**. Distribution of plant Climate Fidelity as a function of climate (see Wang et al. 2022).
